# Aberration-aware 3D localization microscopy via self-supervised neural-physics learning

**DOI:** 10.1101/2025.02.16.638552

**Authors:** Shuang Fu, Wei Shi, Eugene A. Katrukha, Xi Chen, Yue Fei, Ke Fang, Ruixiong Wang, Tianlun Zhang, Donghan Ma, Yiming Li

## Abstract

Single-molecule localization microscopy (SMLM) enables volumetric nanoscopy by retrieving 3D molecular positions from engineered 2D fluorescence patterns. However, achieving nanoscale resolution over large axial ranges in complex biological samples remains challenging due to optical aberrations and overlapped single molecules. Here, we introduce LUNAR, a blind SMLM framework that can precisely localize high-density molecules even with inaccurate and highly distorted PSFs. LUNAR employs a self-supervised neural-physics learning strategy that jointly optimizes a physical PSF model and a neural network to infer key molecular parameters, including 3D positions, photon counts, and aberrations. Through simulations and experiments, we show that LUNAR achieves superior robustness to aberrations, high-density performance across diverse PSFs over multiple frames. We showcase its capabilities through whole-cell nanoscopy of mitochondria, nuclear pores, and neuronal cytoskeletons at large imaging depths. By uniting deep learning with physical modeling, LUNAR provides a calibration-free solution for aberration-robust volumetric nanoscopy and establishes a general framework for adaptive, data-driven 3D imaging.

## Introduction

As a member of the super-resolution (SR) microscopy family, single-molecule localization microscopy (SMLM) has improved spatial resolution by an order of magnitude or more beyond the diffraction limit, while maintaining outstanding molecular specificity and high contrast^1,2^. Recent advancements have pushed its resolution down to Ångström scales^3–5^ and enabled whole-cell 3D SR imaging at high throughput ^6,7^, creating unprecedented opportunities in structural cell biology. In SMLM, blinking molecules are localized with a point spread function (PSF) model. The high precision permitted by SMLM requires accurate PSF modeling, which restricts most SMLM applications to sample of interest positioned just above the coverslip, with a limited depth-of-field (DOF).

When imaging deeper within the samples, mismatched refractive index (RI) between the sample medium and objective immersion medium compromises PSF calibration accuracy, degrading the localization accuracy rapidly with increasing imaging depth^8^. Approaches such as spline interpolation^9,10^, phase retrieval^11,12^ and in situ PSF modeling^13,14^ have sought to improve calibration accuracy. Yet, they all rely on isolated point emitters, either fluorescent beads or blinking molecules, that become scarce in dense or deep imaging conditions, especially for the broadened large-DOF PSF^15,16^. This poses significant challenges for accurate whole-cell longitudinal imaging at greater depth, where in situ PSF models are very different from bead-calibrated PSFs on coverslip and isolated emitters are difficult to obtain.

In recent years, deep learning (DL) methods have shown remarkable promise in SMLM tasks, including high-density molecule localization^15,17^, live-cell imaging^18^, aberration estimation^19^, and spatially variant PSF analysis^6^. However, these methods heavily rely on labeled training data and carefully calibrated PSF models. Any mismatch between training and experimental conditions reduces accuracy, limiting generalizability^20,21^, and leaving much of the information in raw data underutilized.

Unsupervised or self-supervised learning offers a promising alternative by learning representations directly from data, but applying these methods to single-molecule localization remains challenging. Although unsupervised/self-supervised learning has been applied to image denoising and deconvolution, these methods were usually designed to directly predict pixelated images with learning mechanisms based on prior knowledge such as spatial redundancy^22,23^, temporal continuity^24^, or fixed physical models^25,26^. Different from these image-to-image transformation tasks, DL-based single-molecule localization is a structured prediction task that requires inferring 3D point clouds from diffraction-limited 2D single-molecule images with nanometer precision. Previous attempt has been limited to simplified Gaussian PSF model which is only valid for few imaging modalities^27^. A data-driven yet physics-aware framework capable of learning accurate experimental PSFs and emitter parameters directly from raw SMLM data has remained elusive.

To address this gap, we developed LUNAR (Localization Using Neural-physics Adaptive Reconstruction), a blind SMLM method for high-density 3D localization under unknown aberrations. Unlike existing in situ PSF calibration tools like INSPR^13^, DL-AO^19^ and uiPSF^14^ which requires isolated emitters, LUNAR extracts information directly from raw SMLM data even in the presence of overlapped single molecules (**Supplementary Table 1**). By combining CNNs and Transformers, it leverages both spatial and temporal features of SMLM data, yielding superior accuracy and robustness across diverse PSFs, aberrations, and signal-to-noise (SNR) conditions, consistently outperforming state-of-the-art (SOTA) networks on all datasets we investigated, often by a large margin. We demonstrate that LUNAR enables high-fidelity volumetric nanoscopy of cellular organelles across extended axial ranges, providing access to structures previously inaccessible with conventional localization approaches.

## Results

### Principle of LUNAR

LUNAR is a deep generative model that jointly estimates the 3D positions and optical aberrations of fluorescent molecules directly from the intact SMLM frames, without the need to isolate individual emitters (**Fig. 1**). Inspired by variational autoencoders (VAEs)^28^, LUNAR employs an encoder-decoder architecture with intermediate outputs as latent variables. However, unlike typical VAEs that rely solely on neural networks, LUNAR integrates a neural network-based encoder with a physical model-based decoder (**Supplementary Fig. 1** and **Methods**), which we term neural-physics model. This design ensures that the generation process strictly follows physical laws, with intermediate outputs having explicit meanings. Since both the neural network and the physical model are learned together using the unlabeled experimental data, we call this method self-supervised neural-physics learning (**Fig. 1c,d** and **Supplementary Video 1**).

**Fig. 1.**
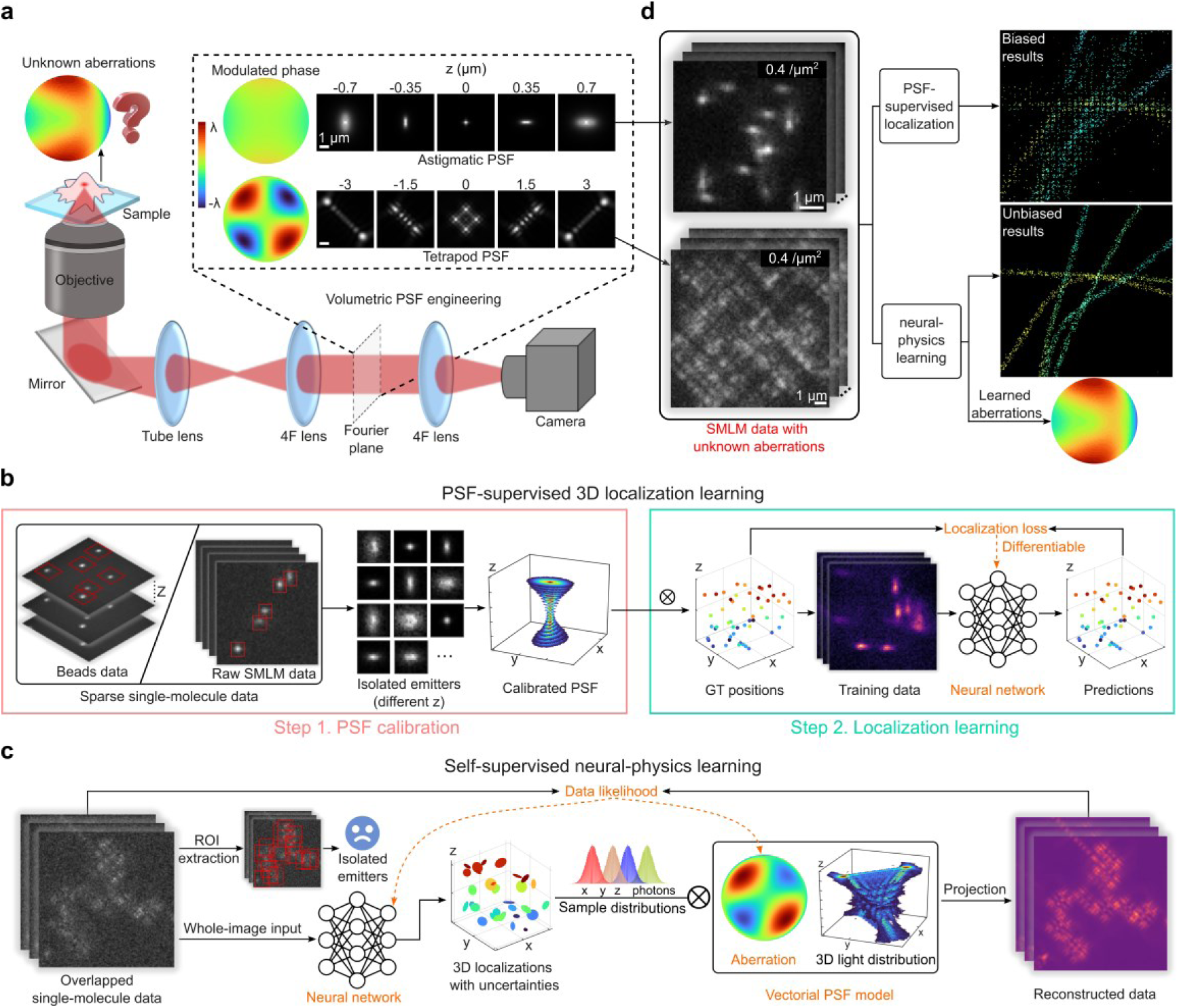
Concept of LUNAR for aberration-robust, high-density volumetric nanoscopy. **a**, 3D SMLM relies on precise estimation of emitter positions from engineered PSF shapes, but sampled-induced aberrations introduce unknown distortions and degrade localization accuracy. **b**, Traditional 3D localization learning methods require a pre-calibrated PSF model derived from sparse single-molecule data to supervise the network training. **c**, LUNAR overcomes this limitation by jointly learning a vectorial PSF model and a deep localization network in a self-supervised manner, eliminating the need for sparse emitters and PSF calibration. **d**, Compared to astigmatic PSF, large axial-range Tetrapod PSF exhibits greater overlap at the same emitter density. LUNAR’s neural-physics learning avoids bias and enables accurate localization in the presence of unknown aberrations and emitter overlaps. Scale bars, 1 μm (**a**, **d**).

The multiple-emitter localization task can be viewed as the inverse of the imaging process, solving the posterior distribution 𝑝_𝜃_(ℎ|𝑑) of emitter positions ℎ given the data 𝑑 (**Methods** and **Supplementary Note 1**). 𝜃 represents the generative parameters that need to be learned, and in this work, we set 𝜃 as a vector of Zernike coefficients to represent aberrations of the PSF model (**Methods** and **Supplementary Note 2**). However, this posterior is generally intractable due to the complexity involved in the imaging process. Therefore, we use the network encoder to learn an approximate posterior function 𝑞_𝜙_(ℎ|𝑑) , where 𝜙 denotes the neural network weights (**Supplementary Fig. 2** and **Methods**). Due to the unknown overlapped emitters and intractable posterior, directly maximizing the marginal likelihood 𝑝_𝜃_(𝑑) is infeasible in this context. Instead, we optimize its variational lower bound, which provides an indirect way to maximize the likelihood (**Methods**).

Simultaneously training a physical model and a neural network that approximates its inverse function is not trivial as the two components must be carefully coordinated during optimization to avoid instability. To efficiently train our neural-physics model, we developed a synchronized learning strategy inspired by the reweighted wake-sleep^29^ and expectation-maximization^30^ algorithms (**Supplementary Fig. 2** and **Methods**). The synchronized learning process alternates between two steps: physics learning and localization learning. In the physics learning step, the network encoder receives the raw SMLM frame as the input and outputs the prediction about the emitter posterior. The physics decoder then draws Monte Carlo samples from this posterior to generate synthetic data with importance weights. The reconstruction loss, which measures the probability of the raw data and synthetic data, is computed to update the physics decoder (**Methods**). In the localization learning step, the physics decoder samples the prior to simulate data with known molecule positions. These data are then used to train the network encoder through a probabilistic localization loss (**Methods** and **Supplementary Note 1**). By alternating between these two steps, the intractable likelihood is indirectly maximized, resulting in a refined physics decoder and the corresponding network encoder, which can then be used to analyze the aberrated, overlapped SMLM data. Conceptually, these two steps mirror the E-step and M-step of the EM algorithm, iteratively improving estimates of the latent variables (emitter positions and photon counts) and the model parameters (aberrations) to maximize the data likelihood.

The network encoder in LUNAR plays a crucial role by providing both the posterior for PSF learning and the localizations for SR image reconstruction. Therefore, we designed a decoupled network architecture (**Supplementary Fig. 3** and **Methods**) to effectively capture both the spatial and temporal context of emitting molecules. To capture spatial features, especially for complex or large PSFs, we introduced a feature extraction module (FEM) based on the U-Net^31^ and ConvNeXt architecture^32^ to facilitate precise feature extraction for PSFs with complex shape and various sizes. To exploit temporal information of blinking emitters across frames, we implemented a temporal attention module (TAM) based on the Transformer^33^ architecture. The self-attention mechanism in the Transformer allows spatial features to interact with each other across the time domain, outperforming traditional CNNs in multi-frame localization task (demonstrated in the following sections). Finally, a convolution-based output module is used to predict the position parameters with uncertainties (**Methods**).

### Comprehensive benchmarking of LUNAR

LUNAR is a 3D SMLM method that jointly learns a neural network for high-density localization and a physical PSF model to capture unknown aberrations. To quantitatively evaluate its performance, we comprehensively benchmarked LUNAR against other SOTA methods (i.e., INSPR^13^, uiPSF^14^, DeepSTORM3D^15^, DECODE^17^, FD-DeepLoc^6^) across multiple criteria, including generalization abilities, localization precision, and PSF learning accuracy.

#### LUNAR shows strong generalization across diverse imaging conditions

Generalization of DL models is a critical factor for practical application in biological research. In SMLM, exprimental conditions such as mismatches in PSF models, various emitter densities, and different SNRs during the long-time image acquisition are common. However , these diverse imaging conditions can significantly impact model performance^34^. To test networks’ robustness on PSF model mismatch, we introduce different levels of unknown aberrations to mimic real experimental conditions (**Supplementary Fig. 4,5** and **Supplementary Note 3**). The networks were trained without knowing the ground truth PSF model. LUNAR was benchmarked with 3 widely used DL based localization methods, DeepSTORM3D^15^, DECODE^17^, and FD-DeepLoc^6^ (**Supplementary Note 4**). As shown in **Fig. 2b** and **Supplementary Fig. 6**, LUNAR exhibits great generalization on different level of unknown aberrations, while the accuracy of PSF-supervised methods deteriorates quickly as the unknow aberrations increase. Although FD-DeepLoc incorporates a robust training strategy to mitigate aberration’s influence, LUNAR achieves more than 6-fold reduction in localization error (**Supplementary Fig. 6b**). It is worth noting that LUNAR could learn the aberrations and maintain high 3D efficiency (>80%) even in overlapped single molecule data for both astigmatic PSF (2 emitters/µm^2^) and 6 µm Tetrapod PSF (0.5 emitters/µm^2^) (**Fig. 2c** and **Supplementary Fig. 7**). Notably, the reported densities (emitters/µm^2^) correspond to per-frame “on”-state molecules, whereas the final reconstruction has substantially higher densities after integration over many frames.

**Fig. 2.**
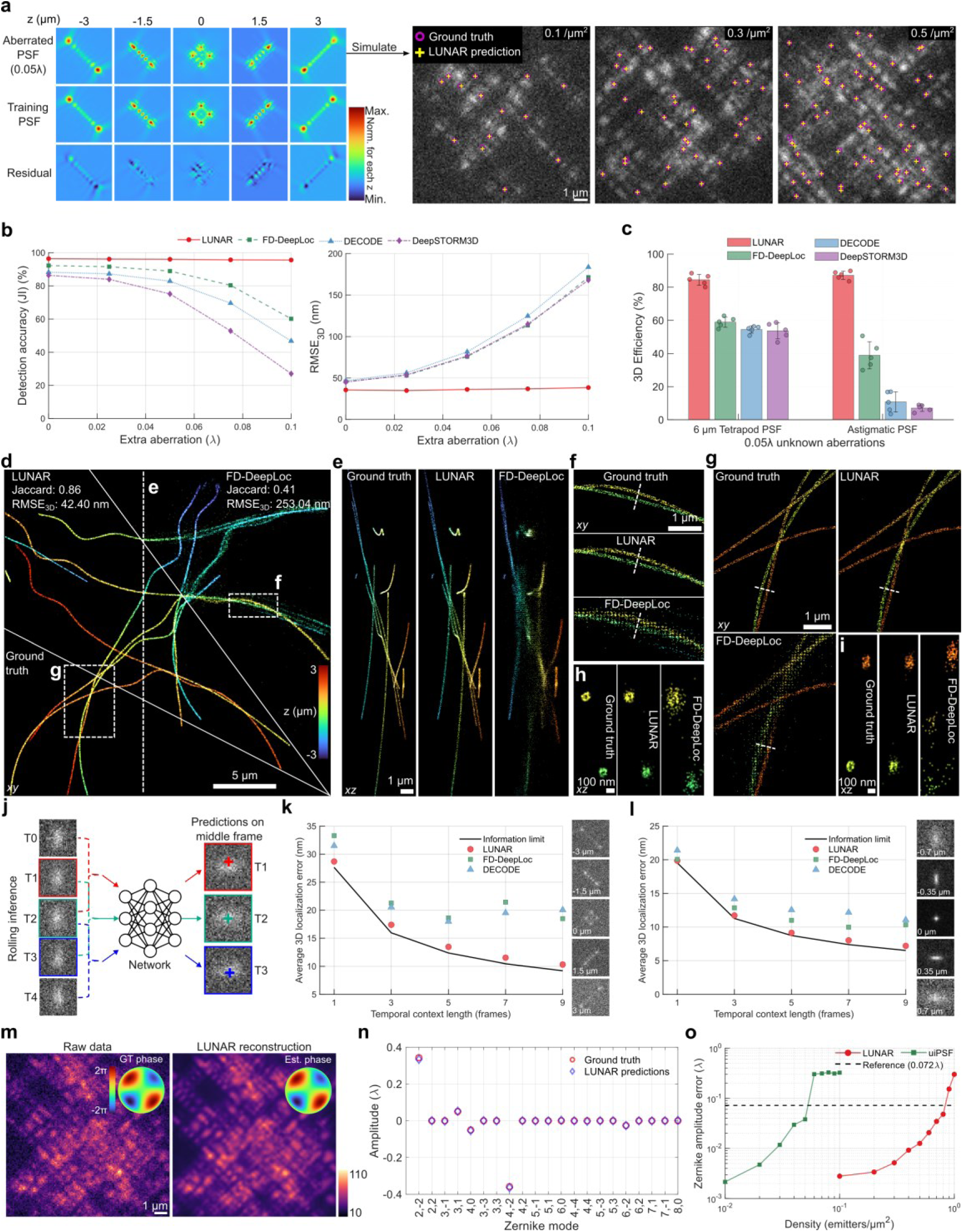
Comprehensive benchmarking of LUNAR. **a**, Example simulated images with unknown aberrations and varying emitter densities. Ground truth (purple) and LUNAR predictions (yellow) are shown for reference. **b**, Detection accuracy (left) and localization error (right) of different methods (LUNAR, FD-DeepLoc, DECODE, DeepSTORM3D) as a function of increasing unknown aberrations, shown for the 6 μm Tetrapod PSF at a density of 0.4 emitters/μm^2^. **c**, Average 3D efficiency across different emitter densities with 0.05λ unknown aberrations (n=5). Error bars indicate the standard deviation. **d**, SR microtubules reconstructed by LUNAR and FD-DeepLoc, with both networks were trained under mismatched conditions (PSF model, emitter density and photophysical model). **e**, 10 μm-width side-view reconstruction along the dashed line in **d**. **f** and **g**, Magnified views of regions denoted by dashed boxes in **d**. **h** and **i**, 800 nm-width side-view reconstructions along the dashed lines in **f** and **g**, respectively. **j**, Schematic of rolling inference using a 3-frame temporal context. In a batch of 5 frames, each frame except the first and last uses its neighbors as contextual input. Spatial features are temporally stored and integrated for temporal relationship computation. **k** and **l**, Average 3D localization error as a function of temporal context length for LUNAR, FD-DeepLoc and DECODE on the 6 μm Tetrapod PSF (**k**) and astigmatic PSF (**l**), respectively. The CRLB is shown as a reference for the information limit. Right insets are example test images at different z positions. Single molecules were simulated with 5,500 photons and 50 background photons per frame; 3,000 frames were simulated at each of 31 z-positions spanning the full axial range of the PSF. All networks were trained using the ground truth PSF model. **m**, Example raw data of the 6 μm Tetrapod PSF and LUNAR’s reconstruction. Insets show corresponding pupil aberrations. **n**, Comparison of Zernike amplitude estimates between ground truth and LUNAR predictions for the dataset shown in **m**. **o**, Zernike amplitude error as a function of increasing emitter densities for in situ PSF learning by uiPSF and LUNAR on the 6 μm Tetrapod PSF. Scale bars, 5 μm (**d**), 1 μm (**a**, **e-g**, **m**), and 100 nm (**h**, **i**).

We then investigated LUNAR’s ability to generalize across various background levels and photon budgets (**Supplementary Figs. 8** and **9**). To streamline training for practical use, we trained each network on a broad SNR range for a given PSF type and tested it on all datasets. Although performance degraded for all methods as background increased or photon counts decreased, LUNAR consistently outperformed the alternatives. Notably, background variation had a larger impact on the Tetrapod PSF than on the astigmatic PSF, especially for structured backgrounds with spatial frequencies higher than the PSF size (**Supplementary Fig. 8f**). This likely reflects the Tetrapod PSF’s broader footprint and lower effective SNR, which increases analysis difficulty. To quantify the photon budget required for acceptable performance, we used 50% 3D efficiency as a benchmark. For the 6 µm Tetrapod PSF, LUNAR exceeded this threshold with only 2000 photons, whereas the other networks required at least 3000 photons when PSF model is known (**Supplementary Fig. 9b,c**). For the astigmatic PSF, LUNAR required 1000 and 1500 photons under known and unknown PSF conditions, respectively, while the other networks did not surpass 50% 3D efficiency when residual aberrations existed.

Finally, we assessed LUNAR’s ability to generalize to complex sample dynamics, spanning a range of SNRs, emitter densities, and dye blinking behaviors. Networks that leverage temporal context (**Supplementary Fig. 3** and **10**) were trained using a simplified photophysical model with fixed emitter density and a broad SNR range (**Methods**). We then tested these models on datasets with varying emitter densities and photon budgets (**Supplementary Fig. 11**). To mimic the real experimental conditions, we employed the same photophysical model as the SMLM challenge^35^, with addition of a practical camera noise model (**Supplementary Note 3**). As shown in **Supplementary Fig. 12** and **13**, LUNAR exhibited stable performance and outperformed FD-DeepLoc for all tested PSFs regardless of training temporal context length and emitter density. Notably, FD-DeepLoc showed tenfold larger performance variance across trials compared to LUNAR, indicating its difficulty in converging under challenging conditions, especially for large-DOF PSFs. The performance gap is visually evident in the reconstructed SR images, where LUNAR produced well resolved hollow microtubules structures and less grid artifacts (**Fig. 2d-i**). Collectively, these results demonstrate LUNAR’s strong generalization under complex imaging conditions.

#### LUNAR approaches information limit for PSFs across wide range of spatiotemporal context

In SMLM, the localization precision required is normally one order of magnitude smaller than the pixel size. Although it has been shown that DL-based single-molecule localization could approach the theoretical minimum uncertainty (CRLB, **Supplementary Note 5**) for small astigmatic PSFs in a single frame^17^, it is still challenging to effectively extract the structural information for PSFs across a wide range of spatiotemporal context. To quantitatively evaluate this, we selected two PSFs: astigmatic PSF and 6 μm Tetrapod PSF, representing different sizes and DOFs. Single molecules were simulated across multiple frames (1/3/5/7/9) to investigate networks’ capability of utilizing temporal information for single-molecule localization.

Localization accuracy was quantified using the 3D root mean square error (RMSE_3D_, **Supplementary Note 6**), with RMSE_3D, avg_ averaged across the full axial range. As shown in **Fig. 2j-l** and **Supplementary Fig. 14**, all networks investigated could approach CRLB for small astigmatic PSFs in single-frame condition. However, if the single molecule appears in multiple frames, FD-DeepLoc and DECODE cannot approach CRLB anymore while the localization accuracy of LUNAR is still close to CRLB across the 1.4 μm axial range investigated for temporal context as high as 9 frames. For 6 μm Tetrapod PSF, which is normally 5 times larger than astigmatic PSF, the RMSE_3D, avg_ of LUNAR (28.7 nm) in a single frame is 14%, 10% and 16% better than that of DeepSTORM3D (32.8 nm), DECODE (31.5 nm) and FD-DeepLoc (33.3 nm), separately. Furthermore, only LUNAR consistently approaches the CRLB with increasing temporal context length (**Fig. 2k,l**). For example, under a temporal context length of 9 frames, the RMSE_3D, avg_ of LUNAR on astigmatic PSF is 7.2 nm, compared to 11.1 nm and 10.3 nm for DECODE and FD-DeepLoc, representing a ∼45% improvement in localization accuracy. The improvement further increased to ∼85% for 6 μm Tetrapod PSF (RMSE_3D, avg_ of LUNAR: 10.3 nm, DECODE: 20.1 nm, FD-DeepLoc: 18.5 nm). We attribute these improvements to the advanced design of LUNAR network, which includes the larger receptive field provided by the U-NeXt and the enhanced ability to capture temporal relationships through the Transformer.

#### Comparison of in situ PSF learning capability of different algorithms

To evaluate how accurate LUNAR can learn in situ PSF from raw data, we first tested it on sparse bead stacks, which offer bright, well-isolated emitters with known axial distance between frames. For comparison, we also evaluated two SOTA in situ PSF learning methods, uiPSF^14^ and INSPR^13^ (**Supplementary Note 4**). Each bead was treated as an individual molecule without using its axial position as prior information. Different PSF learning methods were used to retrieve axial positions of beads, which were then compared to the ground truth objective stage positions. As shown in **Supplementary Fig. 15**, the bead axial positions returned by likelihood-based methods (LUNAR and uiPSF) agreed well with the objective stage positions for both Tetrapod and astigmatic PSFs. In contrast, INSPR was only able to learn the astigmatic PSF and failed on the more complex Tetrapod PSF. We attribute this limitation to its original design for smoothly varying PSFs and the loss of intricate details during the averaging step for PSF retrieval. Next, we evaluated the performance of LUNAR and uiPSF on simulated data as a function of emitter density. Here, the aberration estimation error of 0.072λ was used as a baseline^36^. As shown in **Supplementary Fig. 16**, the aberration estimation error of uiPSF exceeds this baseline for emitter densities of 0.4 and 0.05 emitters/μm² for astigmatic PSF and 6 μm Tetrapod PSF separately. LUNAR could directly learn from the overlapped emitter patterns and process the whole image without single molecule isolation (**Fig. 2m,n**). As a result, LUNAR maintains high PSF model accuracy for emitter densities of 2.6 (astigmatic PSF) and 0.8 (6 μm Tetrapod PSF) emitters/μm² (**Fig. 2o** and **Supplementary Fig. 16c**). To maintain excellent localization performance (>85% efficiency), densities below 0.3 emitters/μm² and 1.6 emitters/μm² for Tetrapod and astigmatic PSF are recommended respectively (**Supplementary Fig. 16d**). LUNAR’s ability to extract PSF model from overlapped single-molecule data makes it a powerful tool to perform 3D nanoscopy in complex imaging conditions as we demonstrate in the following sections with a variety of biological samples.

### LUNAR allows precise 3D nanoscopy with poor PSF calibrations

In typical SMLM laboratories, preparing fluorescent beads for PSF calibration is a routine task. Without regular calibration, it becomes difficult to account for system changes, which can introduce localization errors. Therefore, frequent PSF calibration is essential to maintain accurate and matched PSF models, although this process can be time-consuming and costly.

To illustrate how LUNAR tackles this problem on real biological samples, the widely used reference structures (i.e., microtubules and nuclear pore complex) were imaged. We started with 3D astigmatic imaging of microtubules. For network training, we only introduce astigmatism to the PSF model. Different from the PSF-supervised methods (i.e., FD-DeepLoc, DECODE, DeepSTORM3D), LUNAR was able to learn additional aberrations besides astigmatism (**Supplementary Fig. 17g,h**), enabling accurate reconstruction of the hollow microtubule at different z positions (**Fig. 3c,d**). In contrast, PSF-supervised methods tended to produce more spread localizations and artifacts, especially at large defocus regions where the PSF is deviated from the PSF used for training (**Fig. 3d**).

**Fig. 3.**
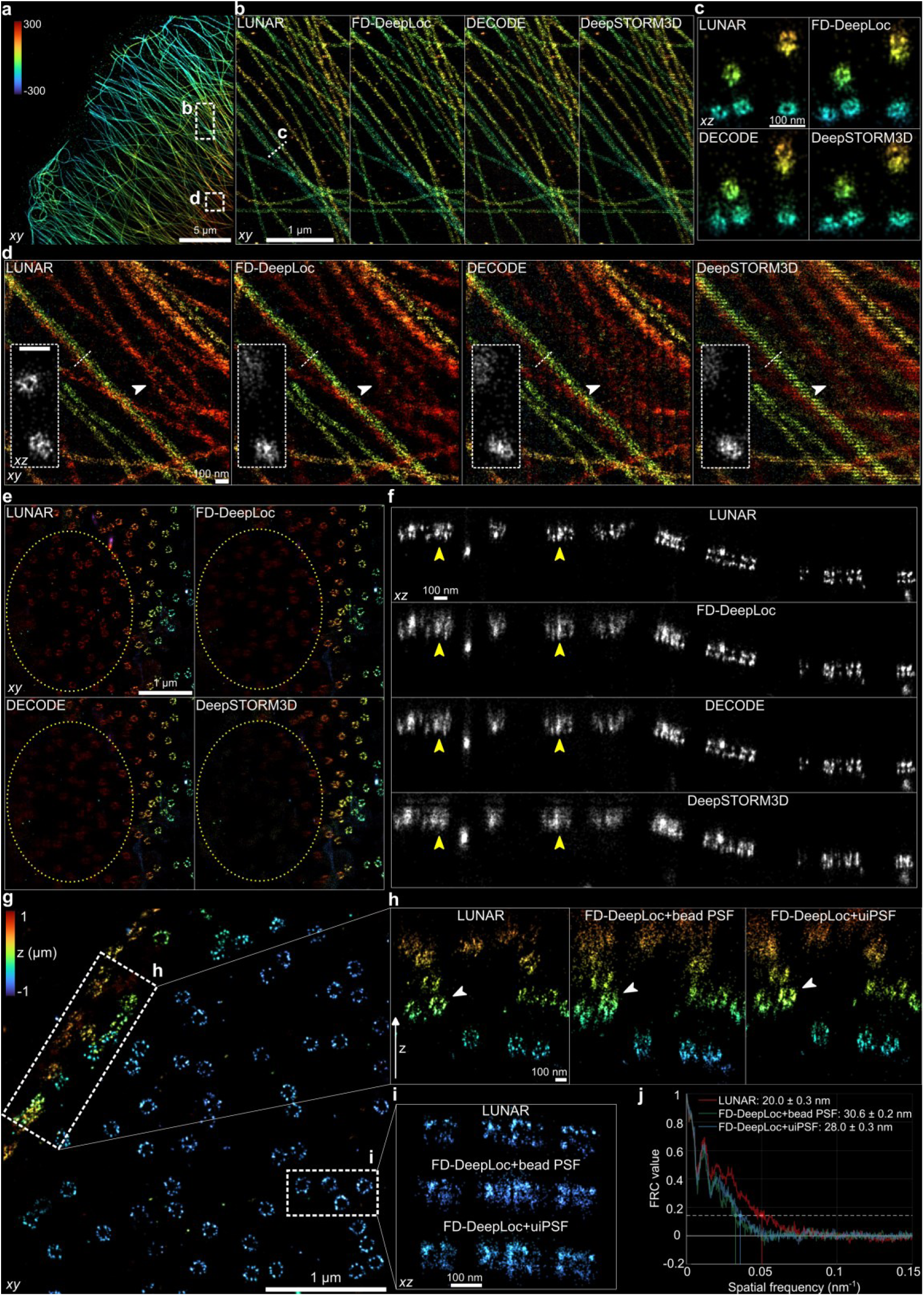
LUNAR enables 3D nanoscopy with poor PSF calibrations. **a**, Super-resolution image of immunolabeled microtubules (β-tubulin, DNA-PAINT, COS-7 cells) reconstructed by LUNAR. **b**, Magnified view of the focus region denoted by the dashed box **b** in **a**, reconstructed by LUNAR, FD-DeepLoc, DECODE and DeepSTORM3D, respectively. **c**, 150 nm-width side-view reconstructions along the dashed lines in **b**. **d**, Magnified view of the defocus region denoted by the dashed box **d** in **a**, reconstructed by different algorithms. Insets are side-view projections along dashed lines. White arrows indicate structures resolved only by LUNAR and FD-DeepLoc. **e**, Representative images of NPCs (Nup96-SNAP, AF647, U2OS cells) collected using an astigmatic PSF and reconstructed by different algorithms. Images are intensity-coded by number of localizations. Yellow dashed circles highlight defocus regions where NPCs are more clearly resolved by LUNAR. **f**, Representative side-view reconstructions by different algorithms. Yellow arrows indicate double-ring structures resolved only by LUNAR. **g**, Representative image of NPCs collected using the DMO PSF and reconstructed by LUNAR. **h** and **i**, Side-view reconstructions of the regions denoted by the dashed boxes in **g**. White arrows indicate ring-like NPC profiles in the side-view of cell nucleus equator. **j**, FRC resolution analysis of the region in **g**. Reported values are mean ± s.d., computed using a threshold of 1/7 and the variance of FRC values. Scale bars, 5 μm (**a**), 1 μm (**b**, **e**, **g**), and 100 nm (**c**, **d**, **f**, **h**, **i**).

We then imaged NPC with a well calibrated bead PSF, where PSF-supervised methods were able to resolve the lateral ring and axial double-ring profiles of NPCs near the focal plane (**Supplementary Fig. 18d**). However, in the upper focal plane that is far from the coverslip, all PSF-supervised methods showed more blurred reconstructions and less detected molecules (**Fig. 3e,f** and **Supplementary Fig.18e**). This is probably due to increased sample-induced aberrations with increase of imaging depth. In contrast, LUNAR reconstructed clear double-ring structures across the entire DOF. Further quantitative analysis of NPCs’ double ring geometry showed that LUNAR was able to correct the z-distortions due to sample-induced aberrations (**Supplementary Fig.18f-h**).

Next, we investigated the performance of the DL-based methods when the PSF model was poorly calibrated. The PSF calibration was performed several weeks before the imaging experiments, potentially leading to PSF model mismatch between calibration and SMLM data. As a result, PSF-supervised learning method (FD-DeepLoc) reconstructed distorted ring structure in the side view of the plane near the equator (**Fig. 3g-i**). LUNAR could iteratively correct the deviated PSF model through its physics decoder during training (**Supplementary Fig. 19** and **Supplementary Video 2)**. For comparison, we also combined FD-DeepLoc with an in situ PSF learning method, uiPSF^14^. When trained using in situ PSF calibrated by uiPSF, FD-DeepLoc was able to resolve NPC ring profiles in the side view reconstruction. However, the reconstructions were more blurred and exhibited lower Fourier ring correlation (FRC) resolution^37^compared to that of LUNAR (**Fig. 3j**), indicating the superior PSF modeling and localization capability of LUNAR compared to the conventional PSF-supervised learning methods.

### Blind whole-cell longitudinal super-resolution imaging with high fidelity

Large-DOF PSFs enables imaging of the whole cell in a single cycle without scanning^16,38,39^ (**Fig. 4a,b**). However, large-DOF PSFs often exhibit laterally extended shapes with reduced SNR. Furthermore, the high probability of overlapping emitter patterns poses significant challenges for computational analysis^15^. Although DL methods have demonstrated great capability in analyzing such data, they have predominantly relied on PSF-supervised learning using pre-calibrated PSF models. A major obstacle for precise large-DOF SMLM imaging is that it is difficult to estimate accurate PSF model from the overlapped single-molecule data.

**Fig. 4.**
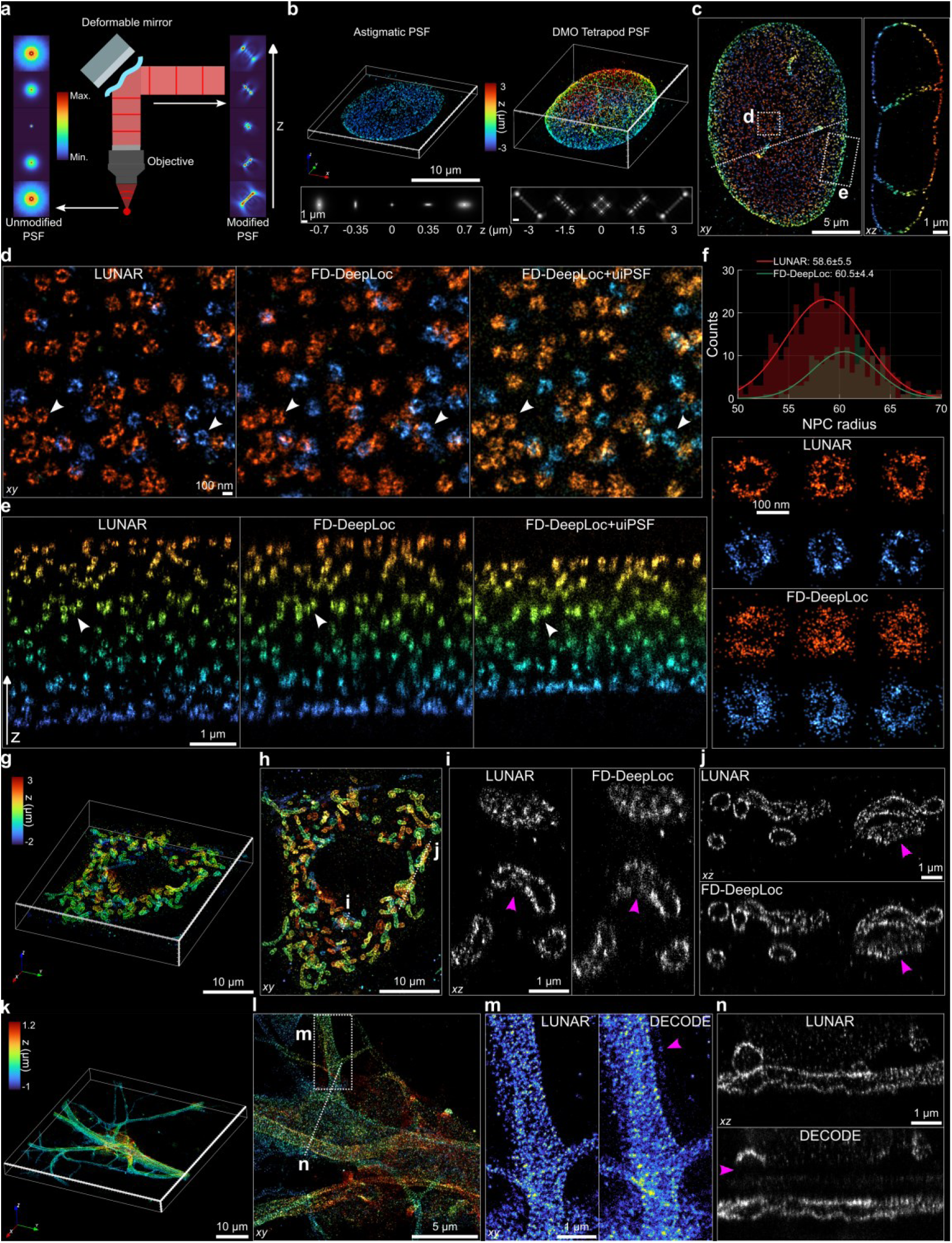
Whole-cell imaging of NPCs, mitochondria and neurons using large-DOF PSF engineering. **a**, Schematic of deformable mirror-based modulation for large-DOF PSF engineering. **b**, DOF comparison between the astigmatic PSF and the 6 μm DMO Tetrapod PSF when imaging the NPCs. **c**, Whole-cell reconstruction of NPCs (Nup96-SNAP, AF647, U2OS cells) by LUNAR. Right: top-view. Left: 1000 nm-wide side-view along the dashed line in the right panel. **d** and **e**, Magnified views (**d**) and side-view reconstructions (**e**) of the regions denoted by the dashed boxes in **c**, reconstructed by LUNAR, FD-DeepLoc and FD-DeepLoc+uiPSF. White arrows highlight superior resolution of NPCs at both the lateral and axial surfaces of the nuclear envelope achieved by LUNAR. **f**, NPC radius fitting for the top half of the nuclear envelope. Top inset: histograms of valid fitted radii from LUNAR and FD-DeepLoc reconstructions. Sample size: *n*_LUNAR_=515, *n*_FD-DeepLoc_=191 NPCs. Bottom inset: example NPC candidates used for radius fitting. LUNAR was trained using a theoretical Tetrapod PSF model while FD-DeepLoc was trained using calibrated PSFs. **g** and **h**, 3D perspective (**g**) and top-view (**h**) of whole-cell mitochondrial reconstruction by LUNAR (Tom20, AF647, COS-7 cells). **i** and **j**, 500 nm-width side-view reconstructions along the dashed lines in **h**, reconstructed by LUNAR and FD-DeepLoc. Magenta arrows indicate punctate protein aggregates more clearly resolved by LUNAR. **k** and **l**, 3D perspective (**k**) and representative top-view (**l**) of whole-cell neuronal reconstruction by LUNAR (βII-spectrin, AF647, mESC-derived neuronal cells). **m** and **n**, Magnified view (**m**) and 1000 nm-width side-view (**n**) of the regions denoted by the dashed box and line in **l**, comparing LUNAR and DECODE. Magenta arrows denote reconstruction artifacts and missing structures. Scale bars, 10 μm (**b** top, **g**, **h**, **k**), 5 μm (**c** left, **l**), 1 μm (**b** bottom, **c** right, **e**, **i**, **j**, **m**, **n**), and 100 nm (**d**, **f**).

Here, we demonstrate precise large-DOF SR imaging by blind data analysis with LUNAR. We first imaged whole-cell NPCs due to their well-defined geometry (**Fig. 4c-f** and **Supplementary Video 3**). Compared to FD-DeepLoc, LUNAR reconstructed sharper images with more punctate localizations, as indicated by the white arrows in **Fig. 4d**. The difference is more profound in the edge areas of the cell nucleus where the ring structure of NPC can be only observed in the side-view images. As shown in **Fig. 4e**, LUNAR provided superior axial reconstructions with clear ring structure on the side view of nuclear envelope over a large axial range, whereas FD-DeepLoc could only resolve disk-like structures. We further trained FD-DeepLoc with an in situ PSF model extracted by uiPSF^14^. However, the side-view image showed strong compression artifact (**Fig. 4e**), likely due to the mismatched PSF model estimated by uiPSF using the contaminated emitter patterns (**Supplementary Fig. 20b**). With improved image quality by LUNAR, we were able to extract more NPCs for quantitative analysis (**Fig. 4f**). Under identical rejection criteria in the automated NPC geometry pipeline in SMAP^40^ (**Supplementary Note 7**), LUNAR retained 515 valid NPCs for ring radius fitting after cleanup, compared to just 191 for FD-DeepLoc. The radii measured by LUNAR aligned closely with previously reported values^41^, whereas dispersed localizations in FD-DeepLoc resulted in slightly larger radius. These results underscore the high fidelity of the images reconstructed by LUNAR for large-DOF single-molecule localization, enabling statistical analysis of super-resolved images with more reliable data.

We next imaged whole-cell mitochondria labeled with TOM20-AF647 (**Fig. 4g**). As shown in **Supplementary Fig. 20c**, LUNAR could reduce the residual error between the raw single-molecule data and the learned reconstruction, indicating the superior model learning capability of LUNAR over FD-DeepLoc. With the help of accurately inferred PSF model and precise large-DOF single-molecule localization, LUNAR clearly reconstructed sharp punctate clusters across the whole cell in the side-view visualization. In contrast, FD-DeepLoc produced blurrier images in which these structures were less distinguishable (**Fig. 4i,j**). We further extended LUNAR to image βII-spectrin in cultured rat neurons (**Fig. 4k-n** and **Supplementary Fig. 21**). Unlike previous observation using astigmatic PSF which is limited to a DOF of a few hundred nanometers^42^, the combination of LUNAR and large-DOF PSF engineering allowed us to visualize the full 3D organization of βII-spectrin across a large axial range without scanning. Through axially separating the reconstruction of densely packed neurites, we were able to clearly resolve the membrane-associated periodic skeletons (MPS) along the whole axon at different depths (**Supplementary Fig. 21d**). In contrast, PSF-supervised methods (FD-DeepLoc and DECODE) exhibited more diffuse localizations and incomplete axial profiles (**Fig. 4m,n** and **Supplementary Fig. 21e**) with lower FRC resolutions and grid artifacts (**Supplementary Fig. 21f**).

### LUNAR enables versatile enhancement of SMLM data across imaging modalities

Sample-induced aberrations often lead to irreversible information loss^19^, which can be recovered through adaptive optics (AO)^43,44^. As a computational framework designed to fully exploit the information contained in the acquired data, LUNAR is compatible with different imaging modalities, enhancing reconstructions from both acquisition and post-processing perspectives. In this section, we validate LUNAR on SMLM datasets acquired using three imaging modalities, demonstrating its broad applicability and great potential for biological investigations.

We first reanalyzed a published imaging dataset acquired from rat brain slices using AO to enhance the SNR of the raw data^44^ (**Fig. 5a-h** and **Supplementary Video 4**). Neurons were labeled with βIV-spectrin, a protein normally located at the axon initial segment (AIS). Notably, imaging was performed at a depth of approximately 50 μm within the tissue, where strong spherical aberrations degraded the elliptical shape of astigmatic PSF^43^. REALM successfully corrected these aberrations using AO, enabling proper 3D reconstruction of the AIS using the classical Gaussian fitting. However, Gaussian PSF model cannot fully describe the in situ PSF inside the complex imaging environments, potentially limiting reconstruction quality. In contrast, LUNAR directly learned the precise PSF from the raw single-molecule data. Leveraging the accurate PSF model inferred and advanced localization network, LUNAR reconstructed sharper structural details and fewer distortions, particularly in side-view cross-sections (**Fig. 5c,d**). The MPS was clearly observed on both the top and bottom of the axon. In side-view projections, we could directly observe the ring-like structures wrapping around the axon circumference and evenly spaced along the axon (**Fig. 5e,f**). The line profiles and their autocorrelation analysis further confirm that LUNAR yields sharper and more structurally informative reconstructions (**Fig. 5g,h**). These results demonstrate that the combination of AO and LUNAR could enable high-quality SR imaging of neuron cytoskeletons inside the tissues, offering new insights into the complex structures of the brain.

**Fig. 5.**
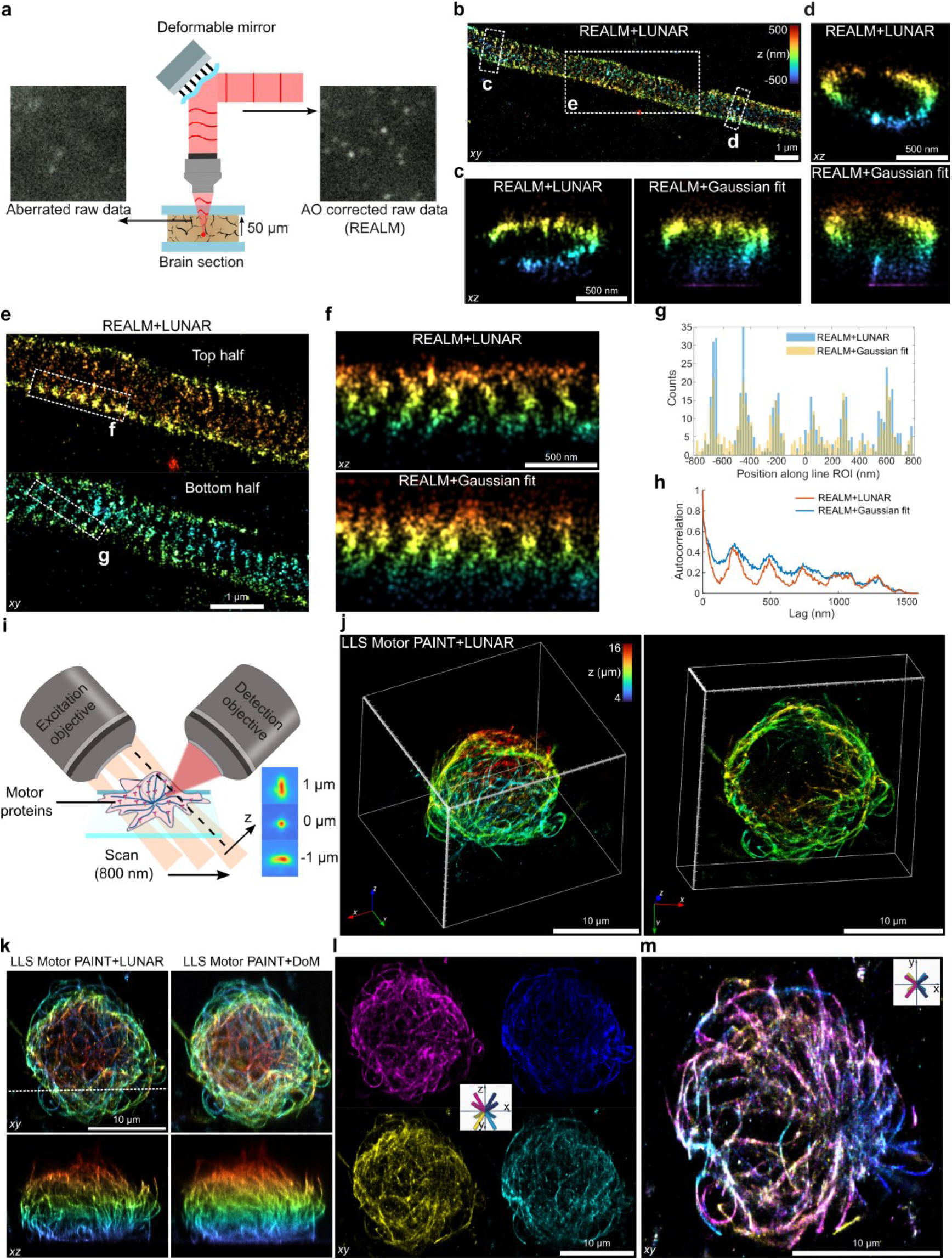
LUNAR enables versatile enhancement of SMLM datasets across diverse modalities. **a**, Schematic of the SMLM dataset collected using an AO system (REALM) for aberration correction at a depth of 50 μm in a rat brain slice. **b**, Overview SR image of an axon (βIV-spectrin, AF647, rat brain slice) reconstructed by LUNAR. **c** and **d**, Side-view reconstructions of the regions indicated by dashed boxes in **b**, using LUNAR and the original published results (Gaussian fit). **e**, Magnified views of the top and bottom halves of the axon in the region indicated by dashed box **e** in **b**. **f**, Comparison of LUNAR and Gaussian fit for the side-view of the region denoted by dashed rectangle **f** in **e**. **g** and **h**, Localization profiles (**g**) of the region denoted by the dashed rectangle **g** in **e**, and corresponding autocorrelations (**h**). **i**, Schematic of the SMLM dataset collected using LLS motor-PAINT to track whole-cell kinesins on microtubules, acquired with an 800 nm scanning step and 57 sections. **j**, 3D perspective of the whole-cell kinesin tracks (DmKHC (1–421)-SNAP-6xHis, JF646, Jurkat cells) reconstructed by LUNAR (localization learning). The right panel shows an alternative view of a truncated volume, with the central dark region corresponding to the nucleus. **k**, Top-view (top row) and side-view (bottom row, along the dashed line in top) reconstructions using LUNAR (left) and the original published results (temporal median filtering + DoM, right). **l**, Microtubule orientation maps derived from kinesin velocity vectors, with color-coded direction. The 3D compass indicates orientation-specific colors. **m**, Sum projection of the bottom 4 μm (near the coverslip) from the top-view, revealing a radial microtubule network originating from the centrosome near the coverslip, with plus-ends oriented outward and upward around the nucleus. Scale bars, 10 μm (**j**-**m**), 1 μm (**b**, **e**), 500 nm (**c**, **d, f**).

Next, we applied LUNAR to a motor-PAINT^45^ dataset imaged with lattice light-sheet (LLS) microscopy. LLS motor-PAINT^46^ was used to investigate microtubule organization and orientation across the entire cellular volume. The whole cell was imaged by scanning a thin light sheet, with in-sheet axial localization achieved by astigmatism (**Fig. 5i**). Fluorescently labeled kinesins bind randomly to the microtubule and run over it for hundreds of nanometers, resulting in dense blinking events (**Supplementary Fig. 22**). Here, we employed LUNAR in localization learning mode, using a bead-calibrated PSF immobilized in samples. As shown in **Fig. 5j**, whole-cell kinesin tracks wrapped around the nucleus across ∼14 µm axial range can be clearly visualized. Compared to the conventional Gaussian fitting based DoM software^47^, LUNAR reconstructs crisper microtubule tracks, especially in side-view projections (**Fig. 5k**). From the tracking results, we can visualize microtubule orientations by assigning colors to localizations based on the direction of the velocity vector (**Fig. 5l**). Analysis of the bottom layers (**Fig. 5m**) and 3D volumes (**Fig. 5j**) reveals that microtubules in Jurkat cells predominantly originate from the centrosome near the coverslip, with plus-ends oriented outward and wrapped around the nucleus.

Finally, we demonstrated LUNAR’s capability in multicolor imaging. As LUNAR is currently designed for single-channel data, we sequentially applied the network to each channel of a public multicolor dataset^48^, and then registered the resulting localizations to assign the colors using the ratiometric method^49^. As shown in **Supplementary Fig. 23a-d**, LUNAR clearly resolved the WGA in 3D, a protein located at the center of the NPC double-ring labeled by Nup107-SNAP-AF647. Owing to LUNAR’s superior detection and localization ability, it yielded more localizations and delivered higher-quality reconstructions than the cubic-spline method implemented in SMAP. A similar improvement was observed in dual-color imaging of mitochondria and microtubules (**Supplementary Fig. 23e-g**), where the contacts between the mitochondria outer membrane protein Tom20 and β-tubulin were readily discernible. However, LUNAR’s current single-channel design limits its ability to fully exploit information distributed across channels in multicolor experiments^50^, future work could extend the framework to multi-channels with self-calibrating capabilities.

## Discussion

In this work, we proposed LUNAR, a blind single-molecule localization framework that jointly learns a neural network and a physical PSF model to enable aberration-robust, high-density 3D nanoscopy. By embedding physical modeling directly into the learning process, LUNAR departs from conventional PSF calibration strategies and supervised deep learning approaches, addressing long-standing challenges of calibration mismatch, aberration distortion, and overlapping emitter patterns.

Our benchmarking demonstrates that LUNAR consistently outperforms existing state-of-the-art algorithms under diverse imaging conditions. Unlike PSF-supervised networks, which degrade rapidly with uncalibrated or distorted PSFs, LUNAR maintains high precision by iteratively refining its internal neural-physics model. Notably, localization accuracy approaches the Cramér–Rao lower bound across both compact and extended PSFs, even when molecules appear over multiple frames. This level of robustness is particularly significant for large-DOF imaging, where PSFs are broadened, SNR is reduced, and emitter overlap is unavoidable.

Applications to biological datasets further validate the advantages of this approach. LUNAR reconstructed detailed subcellular structures such as nuclear pores and mitochondria across extended axial ranges, where conventional calibration-based methods produced blurred or distorted images. The ability to extract accurate PSF models directly from raw data also allowed LUNAR to adapt to imperfect or outdated calibrations, reducing dependence on bead-based procedures. Moreover, compatibility with adaptive optics, lattice light-sheet and multicolor imaging illustrates its flexibility and potential to integrate with diverse hardware innovations. Together, these results suggest that LUNAR can substantially lower experimental barriers while extending the reach of super-resolution imaging into deeper and more heterogeneous environments.

At the same time, limitations should be acknowledged. For highly ill-posed situations LUNAR may converge to a local minimum, which can be solved by incorporating appropriate priors^51^ (**Supplementary Fig. 24** and **Supplementary Note 8**). To ensure reliable results, users should observe specific operational boundaries regarding density and signal quality. Specifically, LUNAR maintains high PSF learning accuracy for emitter densities up to 2.6 and 0.8 emitters/μm² for astigmatic PSF and Tetrapod PSF, respectively. For scenarios involving unknown aberrations, we recommend photon counts exceeding 1500 per emitter, while the spatial frequencies of the background should ideally be larger than the PSF size. Additionally, the current implementation is computationally more intensive than traditional fitting-based or supervised deep learning methods, which may limit throughput and real-time applications. While LUNAR’s neural-physics design captures aberrations effectively, additional developments are needed to handle more complex optical phenomena such as scattering in thick tissues or multimodal PSF behaviors. Finally, accessibility could be improved through streamlined software interfaces and integration with existing analysis pipelines, ensuring broader adoption by the biological imaging community.

In conclusion, LUNAR advances the field of single-molecule localization microscopy by uniting data-driven learning with physical modeling. This integration not only enhances robustness to aberrations and density but also frees researchers from frequent calibrations, enabling high-fidelity volumetric reconstructions in previously inaccessible contexts. Looking ahead, extending the neural-physics framework to other imaging modalities and scaling its computational efficiency will further expand the scope of aberration-robust super-resolution imaging for cell biology.

## Methods

### Derivation of general objective

The overarching goal of LUNAR is to maximize the likelihood of measured SMLM data 𝑑 by simultaneously learning the localization network and the aberrated PSF model. Mathematically, the objective can be expressed as:

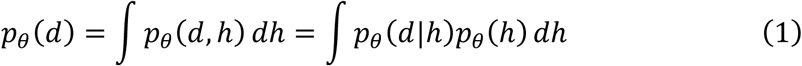

where 𝑝𝜃(𝑑, ℎ) denotes the joint distribution of the measured data 𝑑 and the latent variables ℎ. In this context, ℎ represents the unknown positions and brightnesses of the emitters that need to be localized. The integral marginalizes these latent variables to give the probability of observing the data alone. 𝑝𝜃(ℎ) is the prior distribution. The conditional probability 𝑝𝜃(𝑑|ℎ) , modeled by the physics decoder (PSF model), describes the image formation process of deterministic emitters. The parameters 𝜃 are Zernike coefficients that need to be learned to represent the aberrations (Supplementary Fig. 1a, Supplementary Note 2). The marginal likelihood is intractable due to the complexity of the imaging process and the high-dimensional, continuous nature of ℎ. The task of multiple-emitter localization can be viewed as the inverse of the imaging process, solving the posterior distribution 𝑝𝜃(ℎ|𝑑) = 𝑝𝜃(𝑑, ℎ)/𝑝𝜃(𝑑) of emitter positions ℎ given the data 𝑑, which is also intractable.

Therefore, we use a neural network to learn an approximate posterior function, denoted as 𝑞𝜙(ℎ|𝑑) , where 𝜙 represents the network’s parameters. By introducing this network encoder, we can now optimize the evidence lower bound (ELBO) on the log-likelihood of the data:

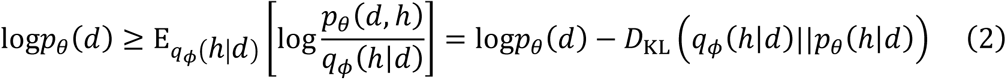

Here, the right side of the inequality is the ELBO. The equality holds true when the approximate posterior 𝑞𝜙(ℎ|𝑑) exactly matches the true posterior 𝑝𝜃(ℎ|𝑑), at which point the non-negative Kullback-Leibler (KL) divergence 𝐷KL(𝑞𝜙||𝑝𝜃) becomes zero. Therefore, our general objective is to train the network to approximate the true posterior as closely as possible, thereby maximizing the log-likelihood of the observed data by iteratively updating the PSF parameters 𝜃.

### LUNAR architecture

LUNAR employs an autoencoder architecture inspired by the VAE^28^, leveraging a probabilistic framework to interpret experimental data while accounting for inherent uncertainties. Different from traditional VAEs, which use abstract latent variables and a network-based decoder to generate data, LUNAR incorporates a learnable vectorial PSF model^52^ as its decoder. This has two benefits: (1) The data generated strictly obey the light diffraction theory. (2) The latent variables are physically interpretable, representing emitters with different positions and photons. After proper training, the network encoder can be used separately to make inferences on SMLM data, localizing molecules with uncertainty estimation.

Network encoder. The LUNAR network is comprised of three modules: the feature extraction module (FEM), the temporal attention module (TAM) and the output module (Supplementary Fig. 3). The FEM employs a U-NeXt backbone, integrating the design principles of ConvNeXt^32^ into the U-Net^31^ architecture. We implement several ConvNeXt strategies to enhance the network performance on single-molecule localization: 1) Adjusting the stage compute ratio to 1:1:3:1 in the FEM; 2) Replacing the 3×3 convolution with a 5×5 depth wise convolution combined with an inverted bottleneck; 3) Replacing ReLU with GELU as the activation function and using fewer activations.

The FEM outputs spatial features for each input frame in a batch, where each pixel in the spatial dimension corresponds to a batch-length time sequence. These sequences are then fed to the TAM, which is a Transformer block containing a masked self-attention layer and a feed-forward layer^33^. The temporal mask in the self-attention layer enables flexible configuration of the temporal context length that the network can utilize for localization, with a default setting of 7. At the end of the TAM, a temporal dropout layer is applied with a probability of 0.5 to randomly drop this temporal information during training, enhancing the network’s generalization ability. A ConvNeXt block follows to aggregate the temporal information with the initial TAM inputs. Notably, if temporal information is not utilized (i.e., using single frame for inference), the TAM will be skipped.

The output module applies several convolution-based prediction heads to output multi-channel maps of equal size to the input images. For each pixel 𝑘, it outputs: detection probability 𝑝^_𝑘_ indicating whether an emitter exists; lateral sub-pixel offsets 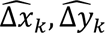 relative to the pixel center; axial distance 𝑧^_𝑘_ relative to the focal plane; brightness 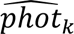; associated uncertainties 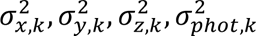; background 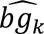. Note that the sub-pixel offsets 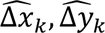, plus the position of pixel 𝑘, form the final lateral position predictions 𝑥^_𝑘_, 𝑦^_𝑘_.

Physics decoder. To reconstruct the raw SMLM data, we applied a learnable vectorial PSF model as the physics decoder for LUNAR. The model is generally formulated as:

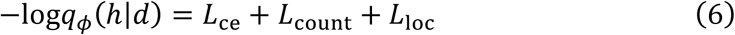

where ℱ_2𝐷_ denotes the 2D Fourier transform. 𝐴(𝜌, 𝜑) is the amplitude function with the apodization factor. 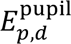 represents the contribution of the fluorescence dipole components 𝑑 = 𝑥, 𝑦, 𝑧 at the sample plane to the light filed components 𝑝 = 𝑥, 𝑦 at the pupil plane. 𝜓_pos_ is the phase shift dependent on the molecule position. 𝜓_aber_ is the phase term that accounts for both PSF engineering and unknown sample-induced aberrations, expressed as a linear sum of multiple Zernike polynomials:

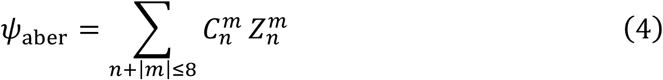

where 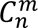 is the Zernike coefficient corresponding to the Zernike polynomial 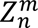, 𝑛 is the radial order, and 𝑚 is the angular frequency. In this work 21 Zernike coefficients 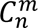 that satisfy 2 < 𝑛 + |𝑚| ≤ 8 are treated as the learnable parameters 𝜃 during the physics learning phase. More details are provided in Supplementary Note 2.

### Synchronized learning strategy

To optimize LUNAR’s model effectively, inspired by the reweighted wake-sleep^29^ and EM^30^ algorithms, we developed a synchronized learning strategy that alternates between two steps: localization learning and physics learning. Each step updates different components of the model while keeping others fixed. Below, we describe the objective functions and gradients computation for each step. Further implementation details and computational cost are provided in Supplementary Note 8.

**In localization learning step**, the physics decoder 𝑝_𝜃_(𝑑|ℎ) is fixed, while the network encoder 𝑞_𝜙_(ℎ|𝑑) learns to approximate the posterior 𝑝_𝜃_(ℎ|𝑑) . This is achieved by minimizing the localization loss, which computes the KL divergence between the posteriors under the physical model and the network, averaged over the training data generated by the current physical model:

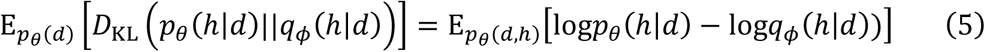

Since 𝜃 is fixed, the gradient of the above objective is estimated by evaluating ∇_𝜙_log𝑞_𝜙_(ℎ|𝑑) , where ℎ, 𝑑∼𝑝_𝜃_(ℎ, 𝑑) . This is equivalent to maximizing the log-likelihood log𝑞_𝜙_(ℎ|𝑑) . In this work, the distribution represented by 𝑞_𝜙_(ℎ|𝑑) is formulated as the joint distribution of three distributions: a Bernoulli distribution for detecting emitters in the pixels, a Gaussian distribution for counting the emitters, and a Gaussian-mixture model (GMM) for emitter parameters regression (Supplementary Note 1.2). This formulation leads to a loss function similar to that of FD-DeepLoc^6^:

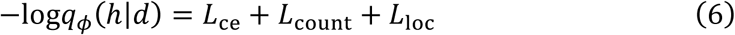

where 𝐿_ce_ computes the cross-entropy between the predicted probability map and the ground truth. 𝐿_count_ computes the negative log-likelihood of the ground truth emitter number under the predicted Gaussian distribution. 𝐿_loc_ computes the negative log-likelihood of the ground truth emitter parameters under the predicted GMM. Additionally, a background term is introduced to form the final loss for localization learning:

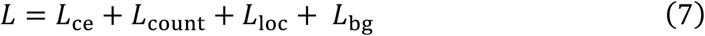

where 𝐿_bg_ is computed using the simple ℓ_2_ norm:

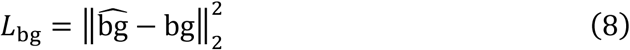

Here, 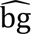 and bg represent the predicted background map and the ground truth background, respectively. This comprehensive loss function ensures that the model not only accurately localizes the molecules but also effectively models the background noise, which is crucial for physics learning.

In physics learning step, the network encoder 𝑞_𝜙_(ℎ|𝑑), which predicts the posterior about emitters’ parameters ℎ = [𝑥, 𝑦, 𝑧, 𝑝ℎ𝑜𝑡𝑜𝑛] given data 𝑑, with parameters 𝜙, is fixed. The physics decoder 𝑝_𝜃_(𝑑|ℎ) with aberration parameters 𝜃 is learnable. The objective of physics learning is given by the ELBO:

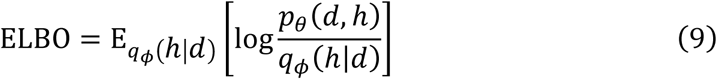

where E_𝑞_[·] represents calculating the expectation under distribution 𝑞 . The joint distribution is defined as 𝑝_𝜃_(𝑑, ℎ) = 𝑝_𝜃_(𝑑|ℎ)𝑝_𝜃_(ℎ) , where 𝑝_𝜃_(ℎ) is the prior distribution of fluorescence molecules. In this work, we make no assumptions about the molecules to be localized and set 𝑝_𝜃_(ℎ) as a uniform distribution.

Since 𝑞_𝜙_ in our framework is represented by the network encoder and is not integrable, we apply the importance sampling (IS) estimator^29^ to get an optimizable objective (whose negative is reconstruction loss):

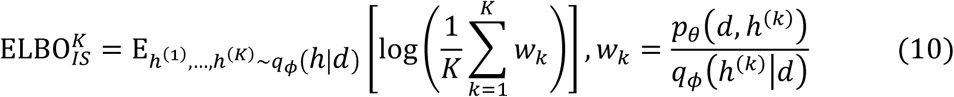

Here, ℎ^(1)^, … , ℎ^(𝐾)^ are sampled independently from the posterior 𝑞_𝜙_(ℎ|𝑑), and we typically set 𝐾 = 100 , which works well for all datasets in this work. Finally, maximizing ELBO_IS_^𝐾^ with respect to the aberration parameters 𝜃 yields the estimated gradients:

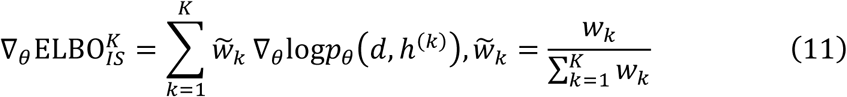

where ̃𝑤_𝑘_ is the normalized importance weight. These gradients are used to update the aberration parameters 𝜃 in the physics decoder, ensuring that the generated data remains consistent with the physical laws governing the SMLM process.

### Training data generation

For physics learning, experimental single-molecule data is used as both training data and targets. For each sampled batch (16 frames in this work), several extra frames are padded at both ends to provide the necessary temporal context for each frame to be reconstructed.

**For localization learning**, training data and targets are generated using the physics decoder. Each unit in the batch (2 used in this work) consists of a series of frames (8 used in this work) with molecules activated using a simplified photophysical model. Specifically, we use a three-state model with off, on and dark states. Transitions between these states follow simple Bernoulli distributions. This is because only a few consecutive images (typically fewer than 16) are simulated in each unit. The molecules with a predefined density in the off state are first randomly activated. The probabilities for on-state molecules surviving in the next frame or dark-state molecules transitioning back to the on-state are determined by:

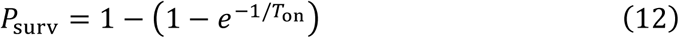

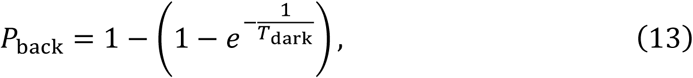

where 𝑇_on_ and 𝑇_dark_ correspond to the average durations of the on and dark states in the photophysical model of the SMLM challenge^35^. In this work, we set 𝑇_on_ = 1 frame and 𝑇_dark_ = 4 frames to account for the multiple decay paths in the more sophisticated challenge model.

For each molecule in the training data, the position and brightness are drawn from uniform distributions within predefined ranges. These ranges are estimated from the raw data or determined based on experience. Although incorporating structure prior about the sample could enhance performance, we decided not to use it to avoid any potential bias. To account for the spatially variant background in real biological specimens, we employ both a constant background and Perlin noise, which are randomly applied to each unit in the training batch. The background range is estimated from experimental data or manually defined.

For camera noise simulation, both EMCCD and sCMOS noise models can be used (**Supplementary Note 3**), which are the same as in our previous work^6^. For the EMCCD camera, the noise model includes shot noise, electron multiplication noise, and camera readout noise. For the sCMOS camera, only shot noise and readout noise are considered, as there is no electron multiplication process. Notably, the synchronized learning currently works only for noise models dominated by Poisson shot noise. If the noise involves more complicated distributions, the loss function for the physics learning should be rederived to accommodate these additional noise components.

### PSF engineering

A deformable mirror (DM140A-35-P01, Boston Micromachines) was used to modulate the PSF for large-DOF experimental imaging. Based on our previous work^38^, we optimized the actuator voltages of the deformable mirror to construct a pupil phase that minimizes the average 3D CRLB within a predefined axial range. For simulation tests, we modulated the PSF pupil using Zernike polynomials. The Zernike coefficient for the astigmatic PSF was 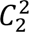 = 70 nm. The optimized Zernike coefficients for the 3 μm Tetrapod PSF were 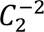 = 120 nm, 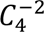 = −130 nm and 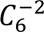 = −6 nm. For the 6 μm Tetrapod PSF, the coefficients were 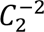 = 230 nm, 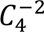 = −240 nm and 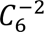 = −17 nm. All simulations were conducted at a wavelength of 670 nm.

### Microscope

In this study, we utilized a custom-built microscope, as described in our previous work^53^. Excitation light from two lasers (iBEAM-SMART-405-S, 150 mW, and iBEAM-SMART-640-S-HP, 200 mW, TOPTICA Photonics) was coupled into a single-mode fiber (P3-405BPM-FC-2, Thorlabs) via a fiber coupler (PAF2-A4A, Thorlabs). By adjusting the translation stage at the fiber output, we can change the illumination angle. The output beam was passed through a laser clean-up filter (ZET405/488/561/640xv2, Chroma) for spectral purification. Beam collimation and reshaping were achieved using a pair of lenses with focal lengths of 75 mm and 400 mm, combined with a slit (SP60, Owis) for beam shaping. The collimated beam was then reflected by a main dichroic mirror (ZT405/488/561/640rpcxt-UF2, Chroma) and directed into the objective for sample illumination. Fluorescence emitted from the sample was collected by a high-NA objective (NA 1.35, UPLSAPO 100 XS or NA 1.5, UPLSAPO 100XOHR, Olympus) and imaged through a tube lens (TTL-180-A, Thorlabs). To confine the imaging field, a slit (SP40, Owis) was placed immediately after the tube lens.

Fluorescence signals were further filtered using two bandpass filters (NF03-405/488/561/635E25 and FF01-676/37-25, Semrock) mounted on a filter wheel, effectively discriminating the emitted fluorescence from the excitation light. A 4f system was integrated into the imaging path using lenses with focal lengths of 125 mm and 75 mm. A deformable mirror (DM140A-35-P01, Boston Micromachines) positioned at the Fourier plane was employed for PSF engineering and aberration correction. To visualize the back focal plane of the objective, a lens with a focal length of 40 mm was placed before the camera. Image acquisition was carried out using an sCMOS camera (Dhyana 400BSI V3, Tucsen or ORCA-Flash4.0 V3, Hamamatsu), which provided a pixel size of 108 nm in the sample plane. Additionally, a closed-loop focus-lock control system was incorporated into the optical setup to ensure stable focus. This system utilized a 785 nm laser (iBEAM-SMART-785-S, 125 mW, TOPTICA Photonics) that was reflected off the coverslip through a dichroic mirror (FF750-SDi02, Semrock) and detected by a quadrant photodiode (SD197-23-21-041, Advanced Photonix Inc). Feedback from the photodiode was used to control a z-piezo stage (P-726.1CD, Physik Instrumente), maintaining precise focus during imaging.

### Sample preparation

#### Cell culture and induced neuronal cells generation

U2OS cells (Nup96-SNAP no. 300444, Cell Line Services) were grown in DMEM (catalog no. 10569, Gibco) containing 10% (v/v) fetal bovine serum (FBS; catalog no. 10099-141C, Gibco), 100 U/ml penicillin and 100 μg/ml streptomycin (PS; catalog no. 15140-122, Gibco) and 1× MEM NEAA (catalog no. 11140-050, Gibco). COS-7 cells (catalog no. 100040, BNCC) were grown in DMEM containing FBS and PS. Cells were cultured in a humidified atmosphere with 5% CO2 at 37 °C and passaged every two or three days. Before cell plating, high-precision 25-mm-round glass coverslips (catalog no. CG15XH, Thorlabs) were cleaned by sequentially sonicating in 1 M potassium hydroxide (KOH), Milli-Q water and ethanol, finally irradiated under ultraviolet light for 30 min. For SR imaging, U2OS and COS-7 cells were cultured on the clean coverslips for 2 days with a confluency of ∼80%.

For the generation of induced neuronal cells, mouse embryonic stem cells (mESCs) were transfected with plasmid containing Tet-on promoter driving expression of rtTA, hygromycin resistance, and Ngn2-P2A-puro. Transfected cells were selected with 200 μg/ml hygromycin (Sigma, V900372) for 3-5 days. On day 0, cells were digested and plated (1×10^5^ cells/well in 6-well plates) on poly-D-lysine/laminin-coated coverslips in serum/LIF medium. After 12 hours, the medium was replaced with N2B27, a 1:1 mixture of DMEM/F12 (Gibco, 11320033) and Neurobasal (Gibco, 21103049), supplemented with 1% N2 (Gibco, 17502048), 2% B27 (Gibco, 17504044), 1×sodium pyruvate, 1×penicillin-streptomycin, 1×NEAA, 1×GlutaMAX, and 1 mM 2-mercaptoethanol, containing 5 μM retinoic acid (Sigma, R2625) and 2 mg/ml doxycycline (TargetMol, T1687). On day 1, puromycin selection (1 mg/ml) was started for 24 hours. On day 6, Ara-C (2 mM, TargetMol, T1272) was added to inhibit neural stem cell proliferation. From day 2 onward, 50% of the medium was exchanged every 2 days, and induced neuronal cells were assayed on day 21 in most experiments.

#### Microtubules labeling

To label microtubules, COS-7 cells were first prefixed in a solution containing 0.3% (v/v) glutaraldehyde (GA) and 0.25% Triton X-100 in cytoskeleton buffer (CB; 10 mM MES, 5 mM glucose, 150 mM NaCl, 5 mM MgCl₂, 5 mM EGTA, pH 6.1), preheated to 37 °C. The prefixation step lasted for 1–2 minutes. The fixative was then removed, and cells were subsequently fixed in 2% GA (in preheated CB) for 10 minutes at 37 °C. After fixation, cells were quenched with 0.1% (w/v) sodium borohydride (NaBH₄; 0.01 g in 10 mL PBS) for 7 minutes, followed by three 5-minute washes with PBS. Cells were permeabilized in 0.1% Triton X-100 in 3% BSA for 10 minutes and washed again three times with PBS. Primary antibody staining was performed using mouse anti-β-tubulin (catalog no. T4026, Sigma, 2.4 mg/mL) diluted 1:1,000 in 3% BSA for 1 h. Cells were then washed three times with PBS for 5 minutes each. Secondary staining was carried out using DNA-PAINT kit antibodies (anti-mouse, MASSIVE-AB 3-PLEX, Massive Photonics), diluted 1:1,000 in the supplied antibody incubation buffer and incubated for 1 h. After staining, cells were washed three times with the washing buffer provided in the kit. Prior to imaging, imager strands were added at a final concentration of 1 nM, pre-diluted in the imaging buffer from the same kit.

#### Nuclear pore complex labeling

To label Nup96, U2OS-Nup96-SNAP cells were prepared as previously reported^40^. Cells were prefixed in 2.4% (w/v) paraformaldehyde (PFA) for 30s, permeabilized with 0.4% (v/v) Triton X-100 for 3 min, and subsequently fixed in 2.4% PFA for 30 min. All fixation and permeabilization steps were conducted using preheated buffers maintained at 37 °C. Following fixation, cells were quenched in 0.1 M NH_4_Cl for 5 min and washed twice with PBS. To minimize nonspecific binding, cells were blocked for 30 min using Image-iT FX Signal Enhancer (catalog no. I36933, Invitrogen). For SNAP-tag labeling, cells were incubated with a dye solution containing 1 μM SNAP-tag ligand BG-AF647 (catalog no. S9136S, New England Biolabs), 1 mM DTT (catalog no. 1111GR005, BioFroxx), and 0.5% (w/v) bovine serum albumin (BSA) in PBS for 2 h. After labeling, cells were washed 3 times with PBS for 5 min each to remove excess dyes. Finally, cells were post-fixed in 4% PFA for 10 min, washed 3 times with PBS for 3 min each, and stored at 4 °C until imaging.

#### Mitochondria labeling

Mitochondrial samples were prepared as previously described^54^. COS-7 cells were fixed in 4% PFA prepared in PBS and preheated to 37 °C for 12 min. The fixed cells were permeabilized for 3 min in permeabilization buffer containing 0.3% IGEPAL CA-630 (catalog no. I8896, Sigma), 0.05% Triton X-100, 0.1% BSA, and 1× PBS. Following permeabilization, the cells were quenched in 0.1 M NH4Cl for 5 min. The cells were then washed 3 times with PBS, each wash lasting 5 min, and subsequently blocked in 3% BSA for 60 min to minimize nonspecific binding.

For primary antibody staining, cells were incubated with rabbit anti-Tom20 (catalog no. ab78547, Abcam, 1 mg/ ml) diluted 1:1,000 in 3% BSA for 2 h. Following this step, the cells were washed 3 times with PBS for 5 min each. Secondary staining was performed using AF647-conjugated secondary antibodies (anti-rabbit, catalog no. A21245, Invitrogen, 2 mg /ml) diluted 1:2,000 in 3% BSA for 2 h. Cells were washed again 3 times with PBS, each wash lasting 5 min. Finally, the cells were post-fixed with 4% PFA for 10 min, washed 3 times with PBS for 5 min each, and stored in PBS at 4 °C.

#### Neuronal cells labeling

Neuronal samples were labeled following a previously described protocol^55^. Cultured mESC-derived neuronal cells were fixed between days 21 and 25 days in vitro using 4% PFA prepared in PBS and preheated to 37 °C. Fixation was performed for 30 min, followed by 3 washes with PBS. The cells were then permeabilized with 0.15% Triton X-100 in PBS for 10 min. After permeabilization, cells were washed 3 times with PBS for 5 min each and subsequently blocked in 3% BSA for 60 min to minimize nonspecific binding. For primary antibody labeling, the cells were incubated with mouse anti-β-spectrin II antibody (catalog no. 612563, BD Biosiences, 250 μg/ml) diluted 1:100 in 3% BSA. The incubation was performed overnight at 4 °C, followed by 3 washes with PBS, each lasting 5 min. Secondary antibody staining was conducted using AF647-conjugated goat anti-mouse IgG (catalog no. A21235, Invitrogen, 2 mg/ml) diluted 1:800 in 3% BSA. The cells were incubated with the secondary antibody for 2 h at room temperature and then washed 3 times with PBS for 5 min each. Finally, the cells were post-fixed with 4% PFA for 20 min, washed 3 times with PBS for 5 min each, and stored in PBS at 4°C until imaging.

#### Dual-color labeling of microtubules and mitochondria

COS-7 cells were first fixed in a solution containing 3% PFA and 0.1% (v/v) GA in CB (10 mM MES, 5 mM glucose, 150 mM NaCl, 5 mM MgCl_2_, 5 mM EGTA, pH 6.1) for 15 min. Cells were then quenched with 0.5% (w/v) NaBH_4_ in PBS for 7 minutes, permeabilized with 0.3% IGEPAL CA-630 (catalog no. I8896, Sigma) and 0.05% Triton X-100 for 3 min, and blocked in 3% BSA for 1h. Primary antibody staining was performed using mouse anti-β-tubulin (catalog no. T4026, Sigma, 2.4 mg/mL) and rabbit anti-Tom20 (catalog no. ab78547, Abcam, 1 mg/mL), diluted 1:600 and 1:1,000 in 3% BSA, respectively, for 2 h at room temperature. Secondary staining was carried out using CF680 (anti-mouse, catalog no. 20817, Biotium) and AF647-conjugated secondary antibodies (anti-rabbit, catalog no. A21245, Invitrogen), diluted 1:250 and 1:2,000 in 3% BSA, respectively, for 2 h. After staining, cells were washed three times with PBS for 5 min each. Finally, cells were post-fixed with 3% PFA and 0.1% (v/v) GA for 10 min, and stored in PBS at 4 °C until imaging.

Imaging Buffer. Samples were imaged in a RI-matched buffer containing 50 mM Tris-HCl (pH 8.0), 10 mM NaCl, 10% (w/v) glucose, 0.5 mg/ml glucose oxidase (catalog no. G7141, Sigma), 40 μg/ml catalase (catalog no. C100, Sigma), 35 mM cysteamine, and 28.5% (v/v) 2,2’-thiodiethanol (catalog no. 166782, Sigma). The final RI of the imaging buffer was 1.406.

Beads preparation. Fluorescent beads with a diameter of 100 nm (custom-designed, Invitrogen) were diluted 1:400,000 in Milli-Q water and vortexed for 3-5 min to ensure uniform dispersion. A 40 μl aliquot of 1 M MgCl_2_ was pipetted onto the center of a 25-mm-diameter coverslip (cleaned as described in the cell culture section) and mixed with 360 μl of the diluted bead solution. The mixture was incubated at room temperature for 5 min, followed by 3 washes with Milli-Q water to remove unbound beads. The coverslip was then stored in Milli-Q water at 4 °C in the dark until use.

## Data availability

The data supporting the findings of this study are publicly available on Zenodo (https://doi.org/10.5281/zenodo.14709467).

## Code availability

Source code of LUNAR is available at: https://github.com/Li-Lab-SUSTech/LUNAR.

## Acknowledgements

This work was supported by the National Key Research and Development Program of China (2024YFF0726003 to Y. L.), Shenzhen Medical Research Fund (B2302038 to Y. L.), National Natural Science Foundation of China (62375116 to Y. L., 623B2044 to S. F.), Key Technology Research and Development Program of Shandong Province (2021CXGC010212 to Y. L.), Shenzhen Science and Technology Innovation Program (JCYJ20220818100416036 and KQTD20200820113012029 to Y. L.), Basic and Applied Basic Research Fund of Guangdong Province (2024A1515011565 to Y. L.), China Postdoctoral Science Foundation (GZC20240651 and 2025T180788 to W. S., 2025T180225, 2025M772887, and GZC20250546 to S. F.), SUSTech Presidential Postdoctoral Fellowship (S. F.), Guangdong Basic and Applied Basic Research Foundation (2022A1515011174 to K. F.), Guangdong Provincial Key Laboratory of Advanced Biomaterials (2022B1212010003), and a startup grant from Southern University of Science and Technology.

## Author contributions

S.F. and Y.L. conceived the project and designed the experiments. Y.L. supervised the entire project. S.F. developed the methods and wrote the software. W.S., Y.F., R.W. and T.Z. tested the software. W.S. and S.F. performed the imaging experiments and analyzed the data. E.K. performed the LLS motor-PAINT data analysis. X.C. and D.M. tested the INSPR. K.F. performed the dual-color imaging experiments. S.F. and Y.L. wrote the manuscript with the input from all other authors.

## Competing interests

The authors declare no competing interests.

## Notes

### Competing Interest Statement

The authors have declared no competing interest.

### Summary of Updates

More quantitative evaluation added; author updated; main figures updated;

