## Supplementary Information for "Aberration-aware 3D localization microscopy via self-supervised neural-physics learning"

### Table of Contents

#### Supplementary Figures


| | |
|---|---|
| Supplementary Fig. 1 | LUNAR architecture |
| Supplementary Fig. 2 | Synchronized learning strategy |
| Supplementary Fig. 3 | Network architecture of LUNAR |
| Supplementary Fig. 4 | Example images of simulated emitters using Tetrapod PSFs |
| Supplementary Fig. 5 | Example images of simulated emitters using astigmatic PSFs |
| Supplementary Fig. 6 | Aberration robustness of different methods on high-density emitters |
| Supplementary Fig. 7 | Performance comparison across different emitter densities |
| Supplementary Fig. 8 | Performance comparison across different backgrounds |
| Supplementary Fig. 9 | Performance comparison across different photons |
| Supplementary Fig. 10 | Network architectures of FD-DeepLoc and DECODE |
| Supplementary Fig. 11 | Example images of simulated microtubule data |
| Supplementary Fig. 12 | Performance of LUNAR and FD-DeepLoc trained with varying temporal context lengths on simulated microtubules |
| Supplementary Fig. 13 | LUNAR exhibits stable performance for unknown PSF model |
| Supplementary Fig. 14 | Evaluation of the localization accuracy using different lengths of temporal context |
| Supplementary Fig. 15 | In situ PSF learning from sparse bead data |
| Supplementary Fig. 16 | Performance of in situ PSF learning across varying emitter densities |
| Supplementary Fig. 17 | Training background range estimation from microtubule dataset and LUNAR learned representations |
| Supplementary Fig. 18 | Super-resolution imaging of nuclear pore complexes (NPCs) using different algorithms |
| Supplementary Fig. 19 | LUNAR enables cross-type PSF correction |
| Supplementary Fig. 20 | Learned representations from whole-cell datasets acquired with a 6 μm DMO Tetrapod PSF |
| Supplementary Fig. 21 | Super-resolution imaging of neurons using different algorithms |
| Supplementary Fig. 22 | Example frames from the lattice light-sheet motor-PAINT dataset |
| Supplementary Fig. 23 | Dual-color super-resolution imaging using LUNAR |
| Supplementary Fig. 24 | Example convergence to a local minimum |


#### Supplementary Tables

| | |
|---|---|
| Supplementary Table 1 | Distinctions between LUNAR and related methods |
| **Supplementary Notes** | |
| Supplementary Note 1 | The development of the LUNAR |
| Supplementary Note 2 | Vectorial PSF model |
| Supplementary Note 3 | Test datasets simulation |
| Supplementary Note 4 | Benchmarking with other methods |
| Supplementary Note 5 | Computation of theoretical localization precision limit |
| Supplementary Note 6 | Evaluation metrics |
| Supplementary Note 7 | Data post-processing, analysis and rendering |
| Supplementary Note 8 | Implementation details about LUNAR |

#### Supplementary Figures

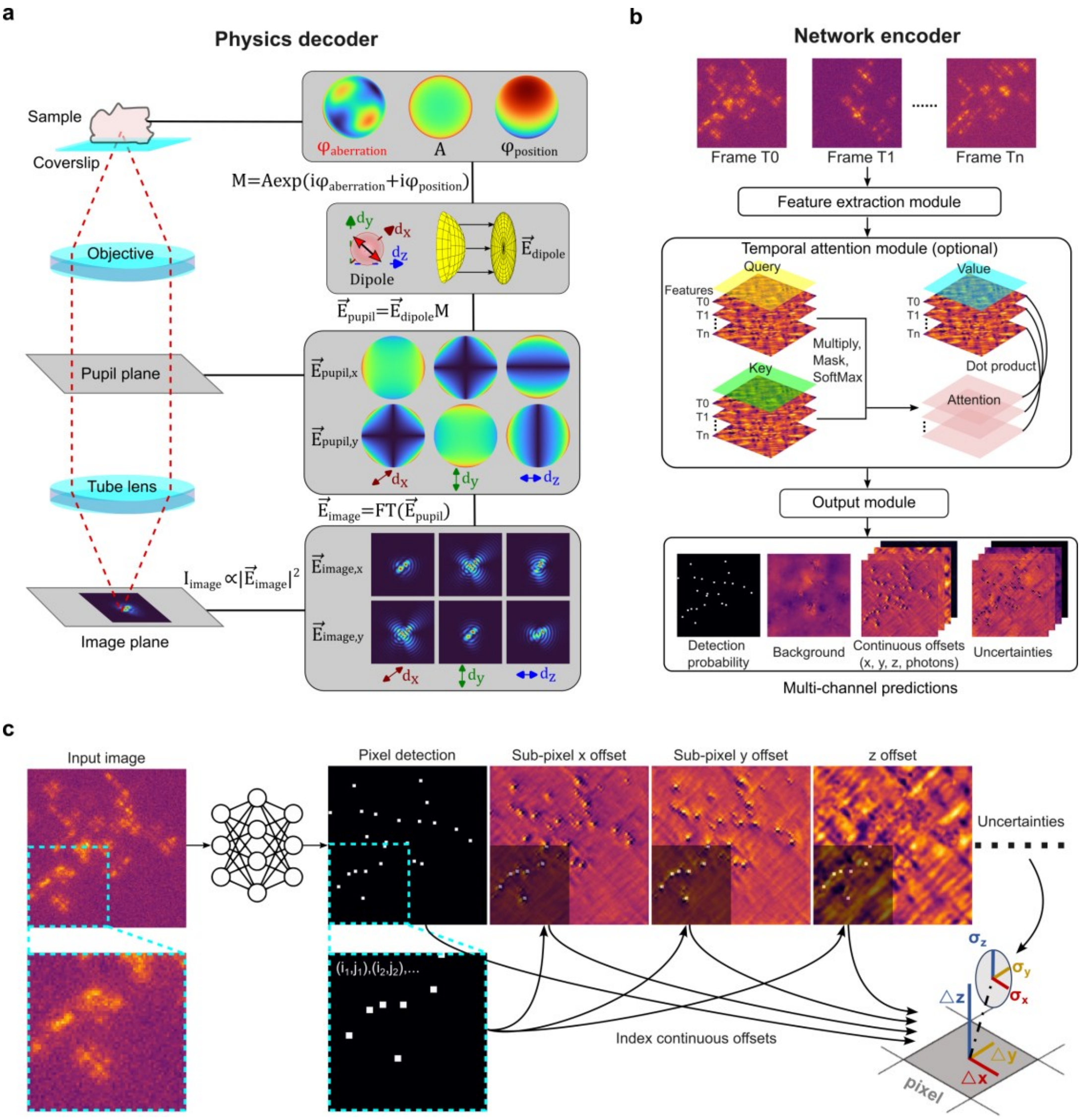


**Supplementary Fig. 1 | LUNAR architecture. a**, The physics decoder is implemented as a vectorial PSF model with a learnable pupil phase. The pupil phase is parameterized using 21 Zernike modes satisfying $2 < n + |m| \leq 8$, where $n$ is the radial order and $m$ is the angular frequency. **b**, The network encoder utilizes a feature extraction module to process each frame independently. When multiple frames are used for prediction with temporal context, an optional Transformer-based temporal attention module captures relationships between consecutive frames. The output module generates multi-channel prediction maps that encode emitter localizations and their associated uncertainties for each frame. **c**, To convert multi-channel predictions into a list of emitter positions, pixel-level detections are first identified by thresholding the probability map. These indices are then used to retrieve continuous sub-pixel offsets and uncertainty estimates.

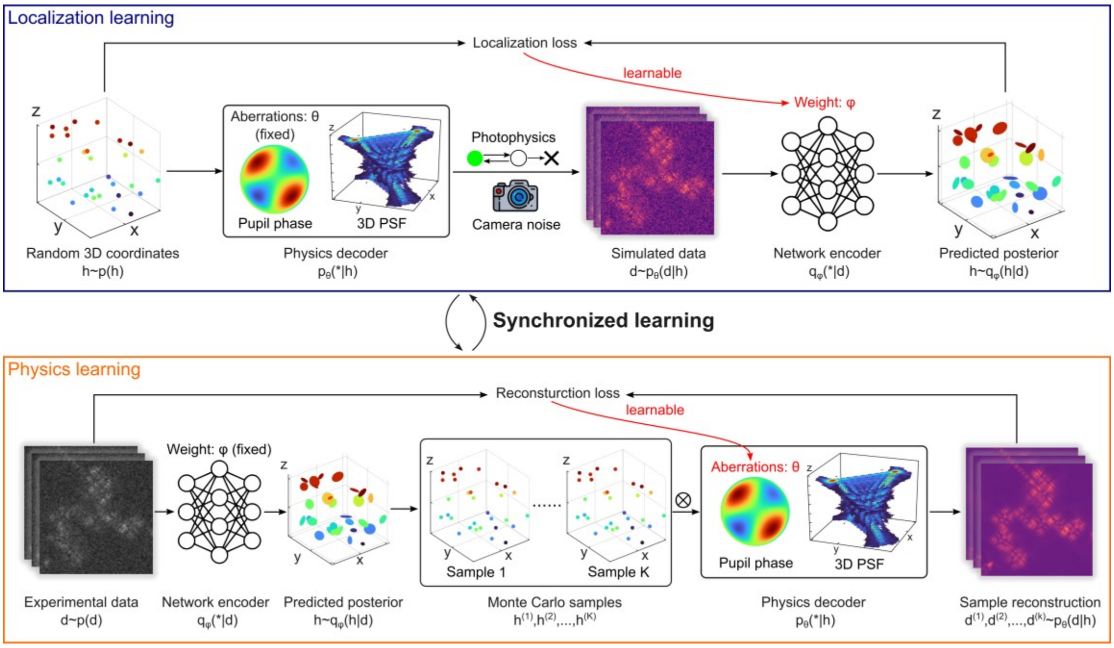


**Supplementary Fig. 2 | Synchronized learning strategy.** LUNAR jointly optimizes its network encoder and physics decoder through an alternating two-step procedure: physics learning and localization learning. During localization learning, the physics decoder is fixed, and the network encoder is updated to minimize the localization loss. During physics learning, the network encoder is fixed, and the physics decoder is updated to minimize the reconstruction loss. See **Methods** for more details.

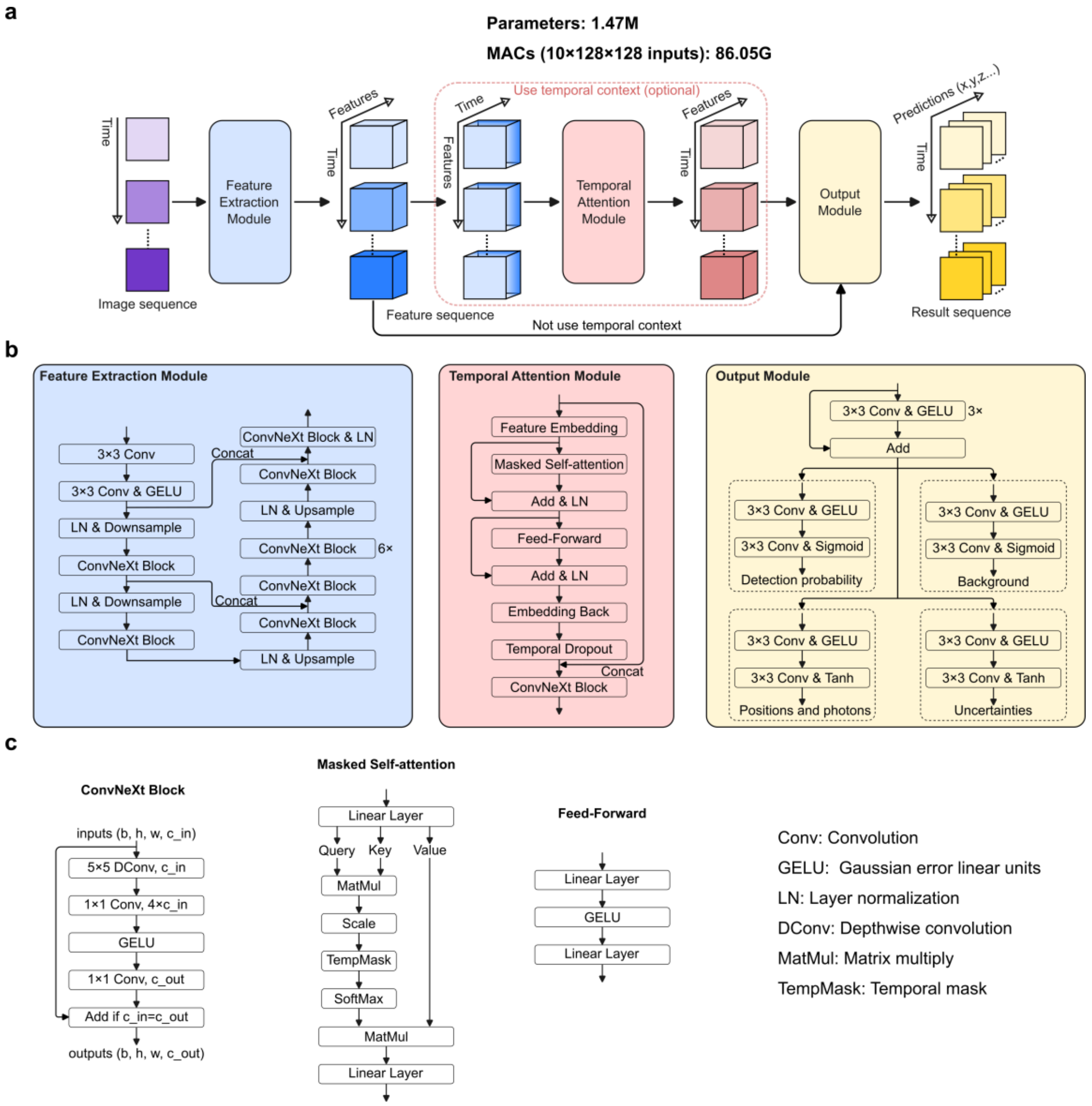


**Supplementary Fig. 3 | Network architecture of LUNAR. a**, Overview of the network design, including its number of parameters and computational workload. **b**, Detailed architectures of the feature extraction module, temporal attention module and output module. **c**, Mechanisms of the ConvNeXt block, masked self-attention and feed-forward layers, with abbreviations explained on the right.

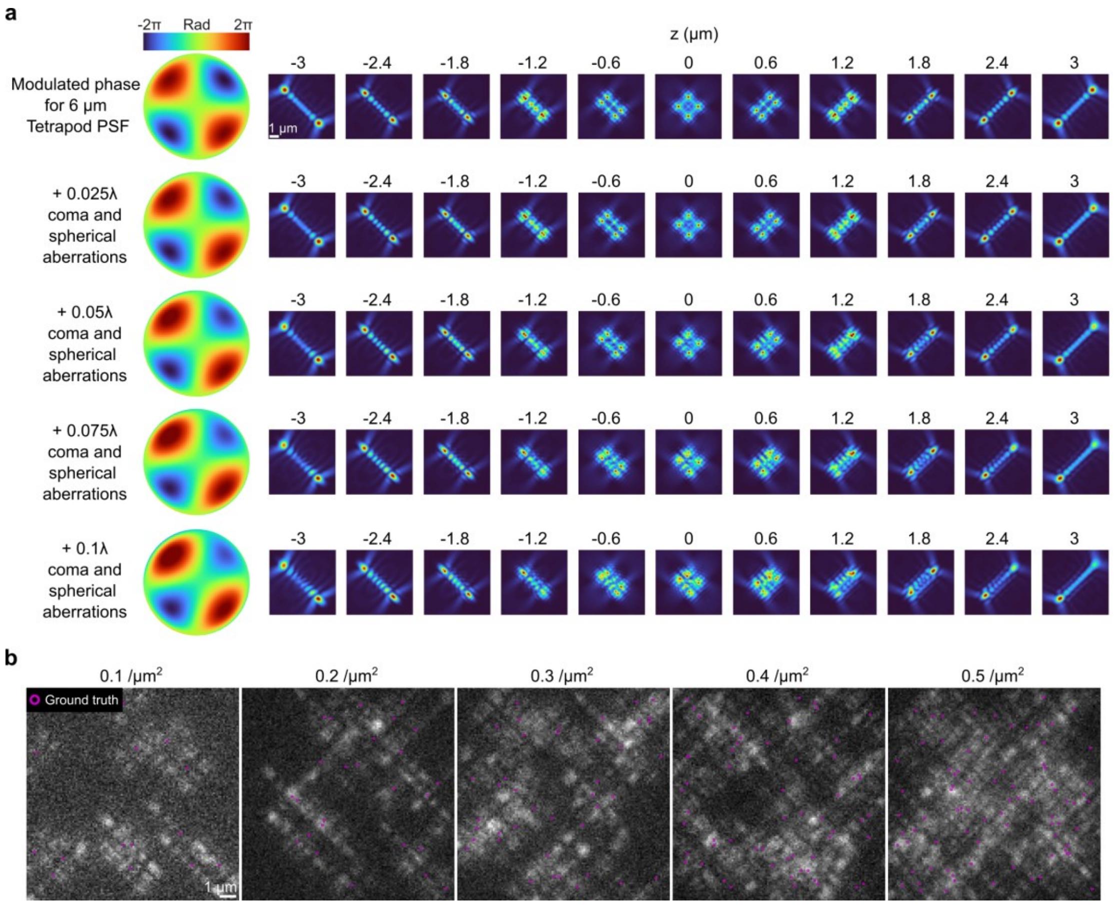


**Supplementary Fig. 4 | Example images of simulated emitters using Tetrapod PSFs.** **a**, Modulated phases and corresponding PSF images at different *z* positions for the 6 μm Tetrapod PSF, showing increasing levels of coma and spherical aberrations from top to bottom (0.025λ, 0.05λ, 0.075λ, 0.1λ). **b**, Example images of simulated datasets with varying emitter densities and an aberration level of 0.05λ. The average photon count is 5000, with a background of 20 photons. Scale bars, 1 μm (**a**, **b**).

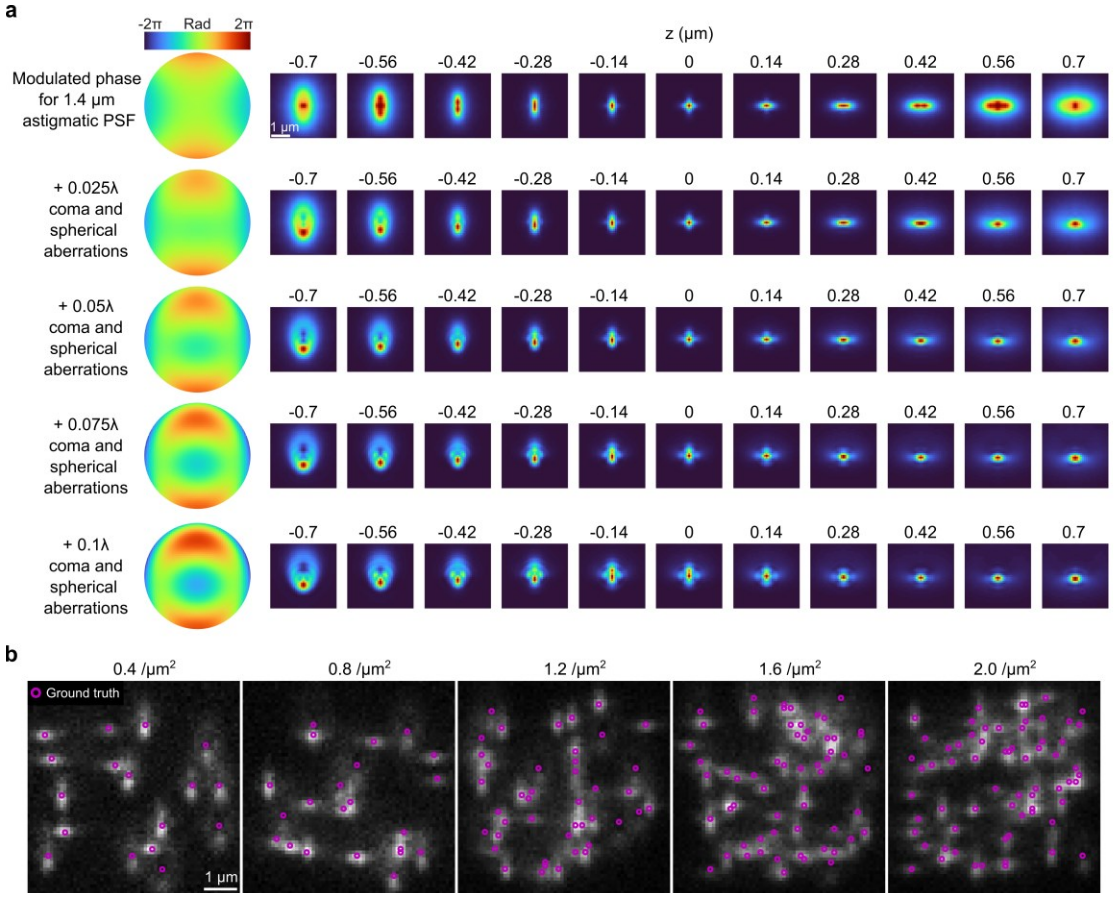


**Supplementary Fig. 5 | Example images of simulated emitters using astigmatic PSFs.** **a**, Modulated phases and corresponding PSF images at different *z* positions for the 1.4 μm astigmatic PSF, showing increasing levels of coma and spherical aberrations from top to bottom (0.025λ, 0.05λ, 0.075λ, 0.1λ). **b**, Example images of simulated datasets with varying emitter densities and an aberration level of 0.05λ. The average photon count is 5000, with a background of 20 photons. Scale bars, 1 μm (**a**, **b**).

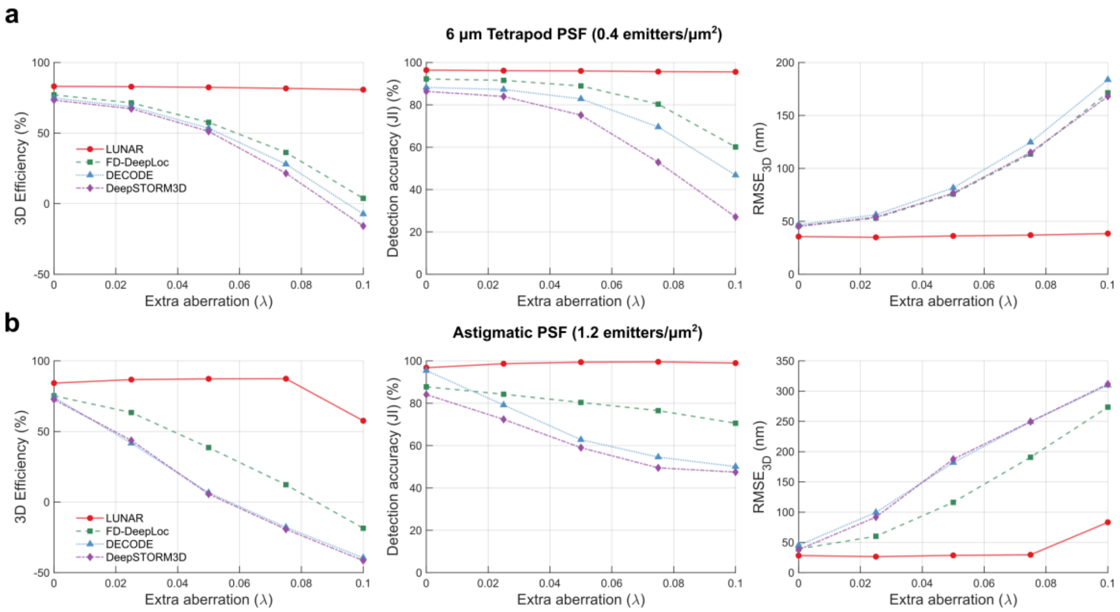


**Supplementary Fig. 6 | Aberration robustness of different methods on high-density emitters. a**, Localization performance as a function of increasing unknown aberrations for the 6 µm Tetrapod PSF and **b** astigmatic PSF, at fixed emitter densities. All methods were provided with the same PSF model that did not include the extra aberrations (0λ). FD-DeepLoc implements a robust training strategy by adding random aberrations to the training data. Performance is evaluated using three metrics (**Supplementary Note 6**): 3D efficiency (higher is better), detection accuracy (Jaccard Index, JI; higher is better), and 3D localization error ($RMSE_{3D}$; lower is better).

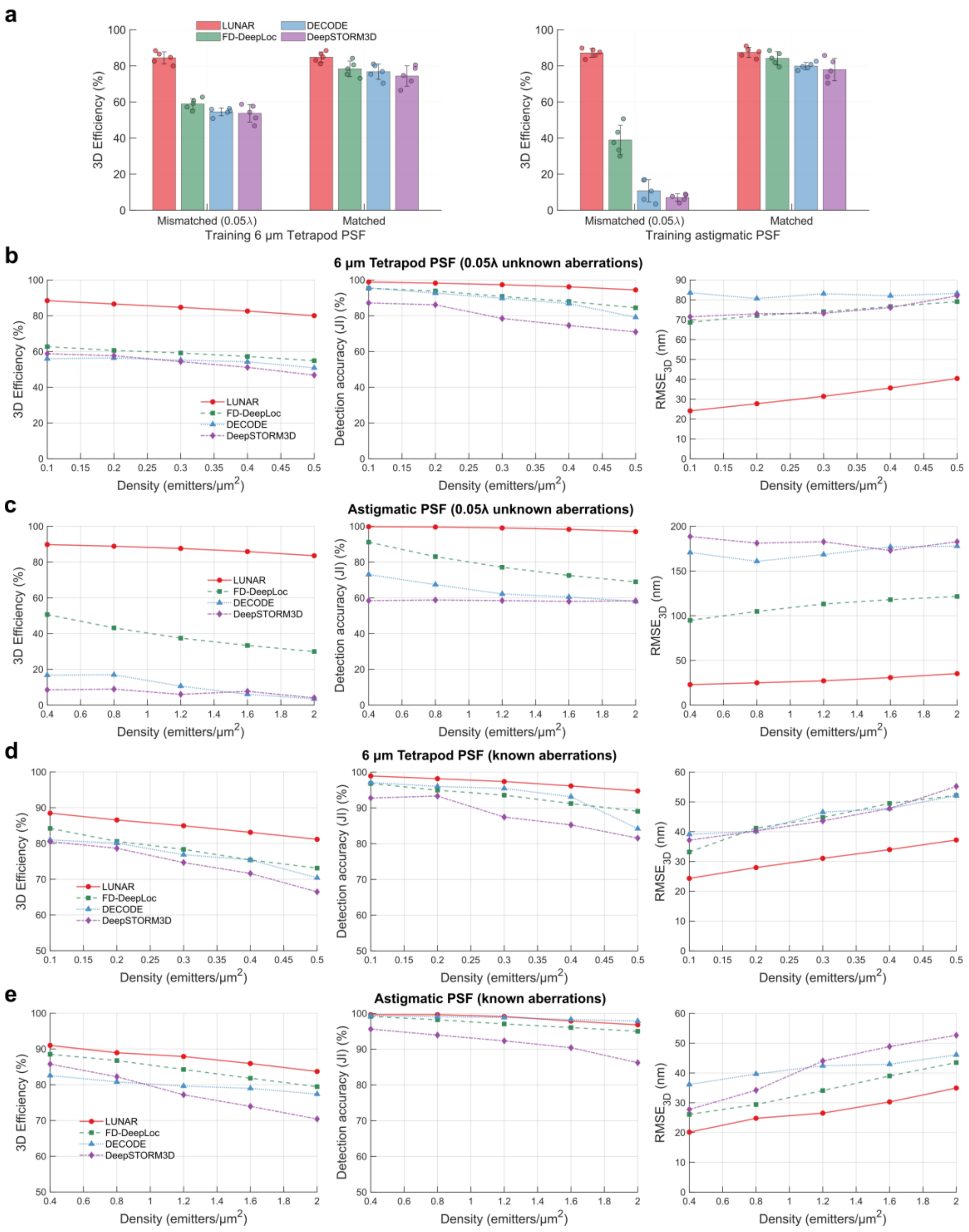


**Supplementary Fig. 7 | Performance comparison across different emitter densities.** **a**, Average 3D efficiency for different methods with and without matched PSFs (n=5). **b** and **c**, Localization performance as a function of increasing emitter density for the 6 μm Tetrapod PSF and astigmatic PSF, respectively, under a fixed level of unknown aberrations (0.05λ). All methods were provided with the same initial PSF model that did not include the additional aberrations (0λ). FD-DeepLoc implemented a robust training strategy. **d** and **e**, same as **b** and **c**, but using the ground truth PSF model.

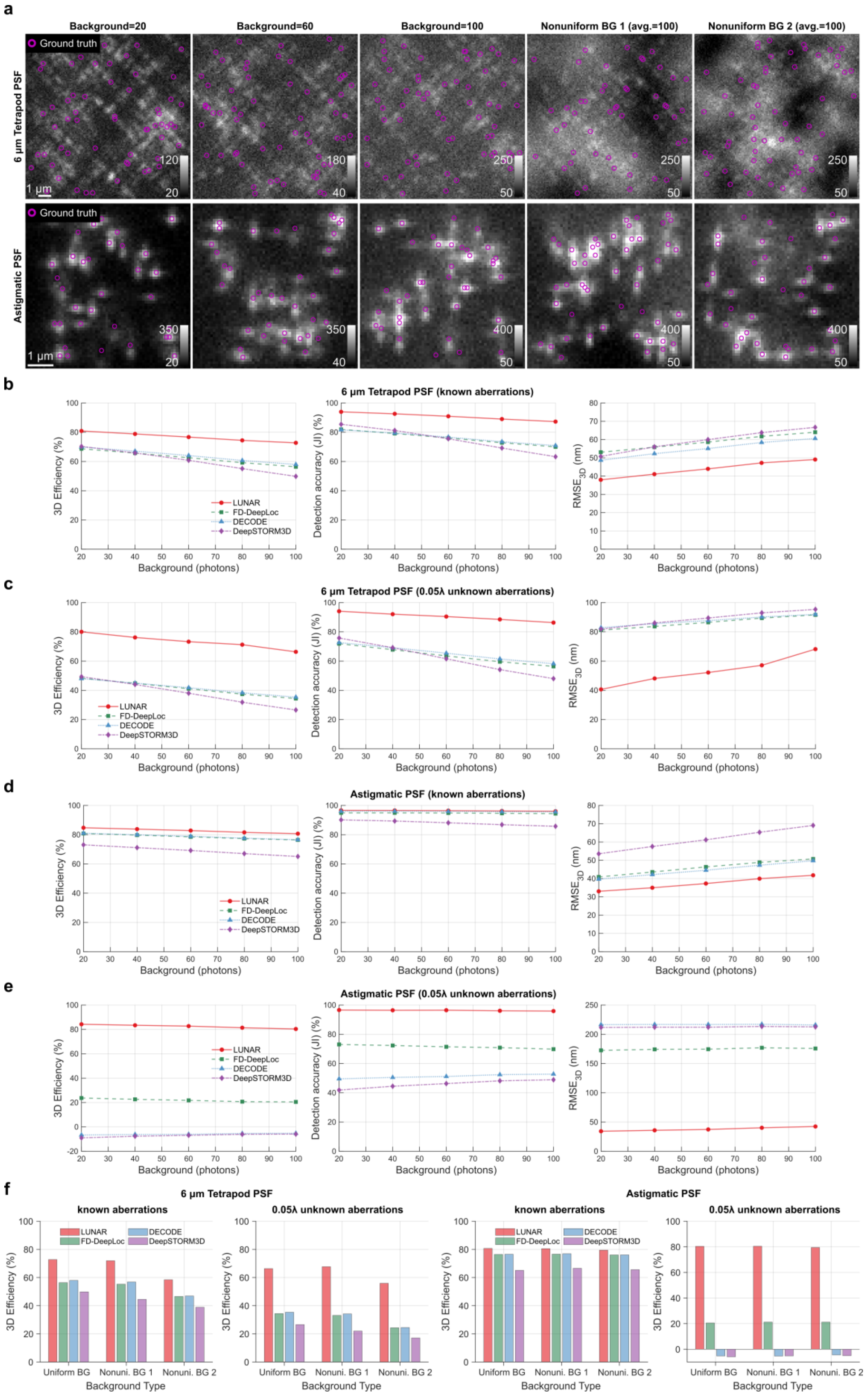


**Supplementary Fig. 8 | Performance comparison across different backgrounds. a,**

Example images of simulated datasets with varying backgrounds. The average photon count is 5000. The density is 0.4 and 1.2 emitters/μm$^2$ for 6 μm Tetrapod PSF and astigmatic PSF, respectively. **b**, Localization performance as a function of increasing backgrounds, under a fixed level of extra aberrations (0.05λ). All methods were provided with the ground truth PSF model. **c**, same as **b**, but all methods were provided with the mismatched PSF model that did not include the additional aberrations (0λ). FD-DeepLoc implemented a robust training strategy. **d** and **e**, same as **b** and **c**, but for the astigmatic PSF. All networks were trained over a photon range of [1000, 10000] and a background range of [10, 150]. **f**, Effects of structured backgrounds. Nonuniform BG 1 and 2 correspond to medium (64 pixels) and high frequencies (32 pixels) of Perlin noise added on the uniform background. Scale bars, 1 μm (**a**).

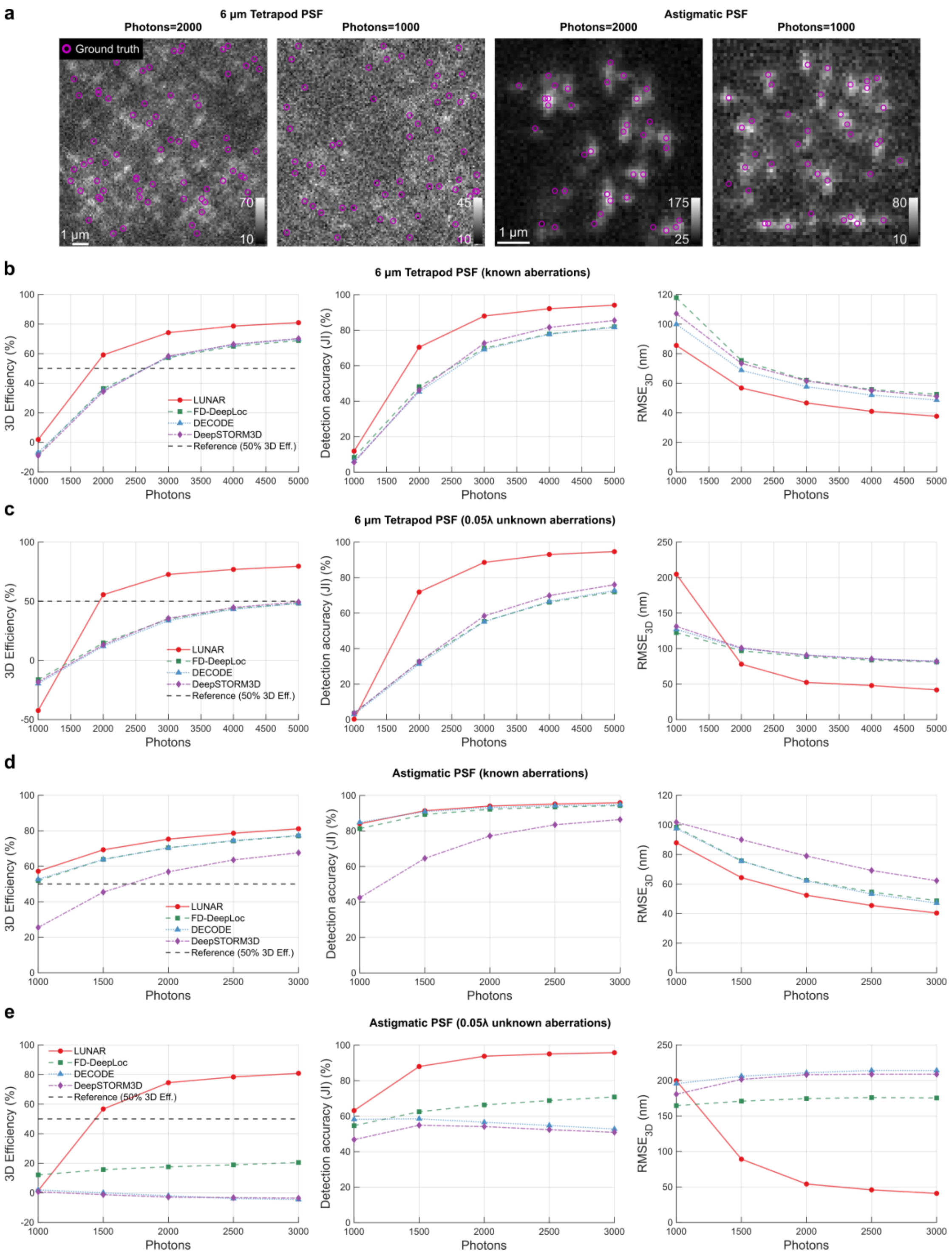


**Supplementary Fig. 9 | Performance comparison across different photons. a**, Example images of test datasets with limited photons. The background is set as a constant of 20 photons. The densities are 0.4 and 1.2 emitters/µm$^2$ for the 6 µm Tetrapod PSF and astigmatic PSF, respectively. **b**, Localization performance as a function of different photons for the 6 µm Tetrapod PSF, under a fixed level of extra aberrations (0.05λ). All methods were provided with the ground truth PSF model. **c**, same as **b**, but all methods were provided with the mismatched PSF model that did not include the additional aberrations (0λ). FD-DeepLoc implemented a robust training strategy. **d** and **e**, same as **b** and **c**, but for the astigmatic PSF. Scale bars, 1 μm (**a**).

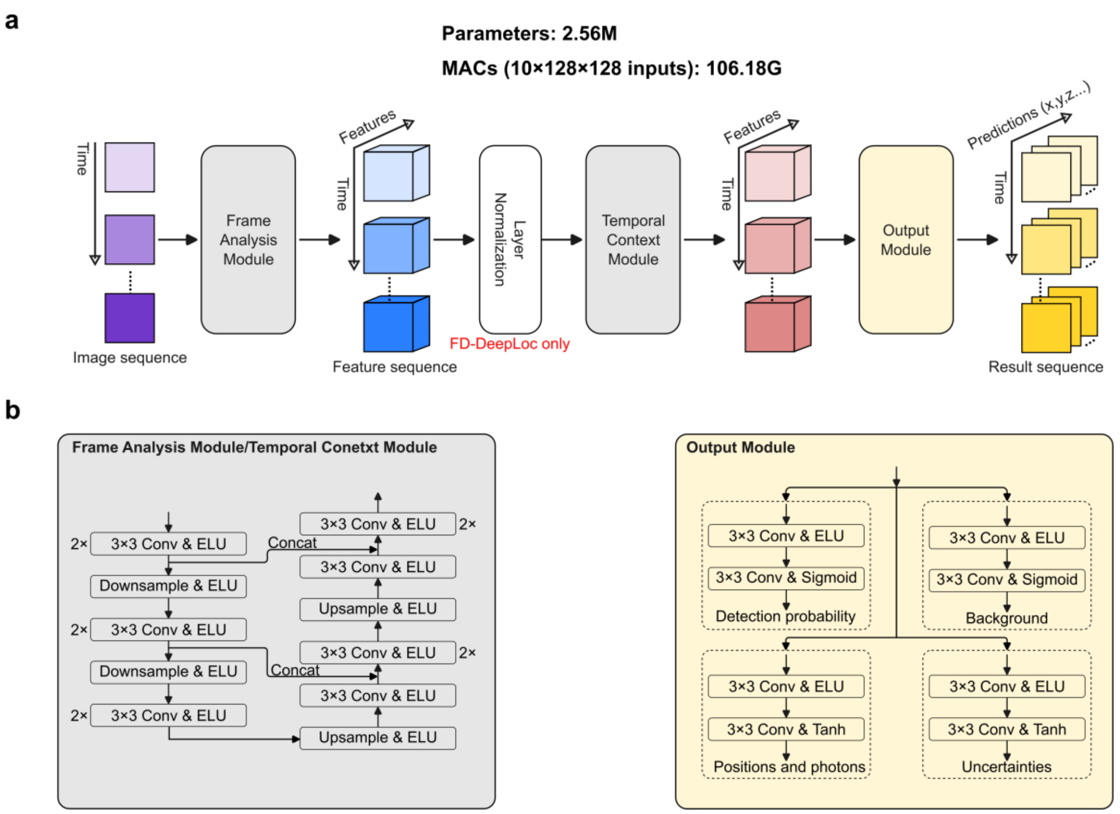


**Supplementary Fig. 10 | Network architectures of FD-DeepLoc and DECODE. a,** Overview showing that the two algorithms employ nearly identical network structures, with the exception that FD-DeepLoc includes an additional layer normalization step. **b,** Detailed architectures of the modules used in **a**. Both the frame analysis module and the temporal context module are based on the same U-Net architecture, differing only in the number of channels in the first convolution layer, which corresponds to the length of temporal context they process.

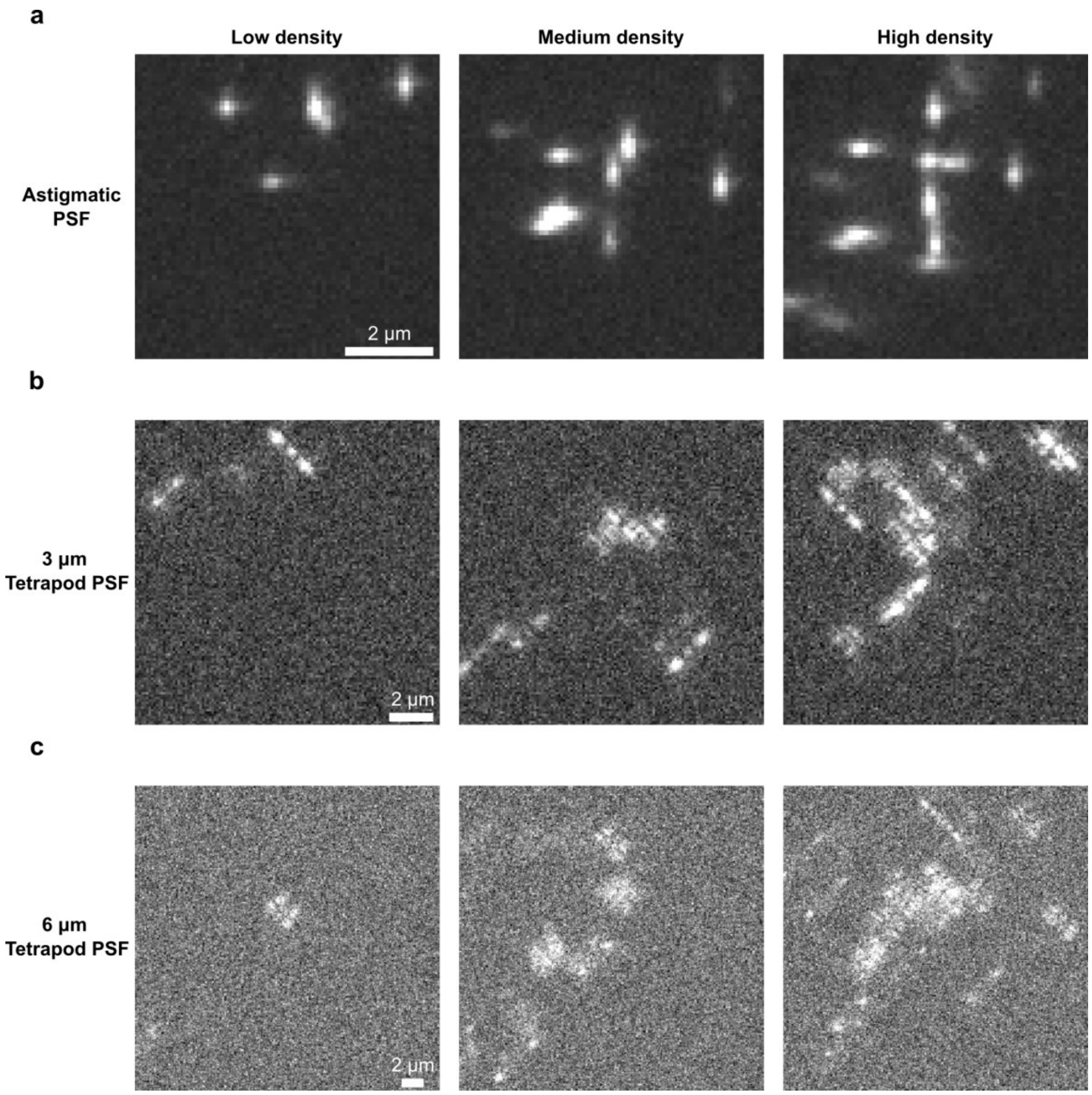


**Supplementary Fig. 11 | Example images of simulated microtubule data. a**, Example images of astigmatic PSF datasets at three density levels, corresponding to 4.5, 9.2 and 18.3 emitters per frame. The exact density values cannot be determined like uniformly distributed emitters in **Supplementary Fig. 4** and **5**, as microtubules only appear in certain area. **b** and **c**, Same as **a**, but simulated using 3 μm Tetrapod PSF and 6 μm Tetrapod PSF, respectively. Details for data simulation can be found in **Supplementary Note 3**. Scale bars, 2 μm (**a**, **b**, **c**).

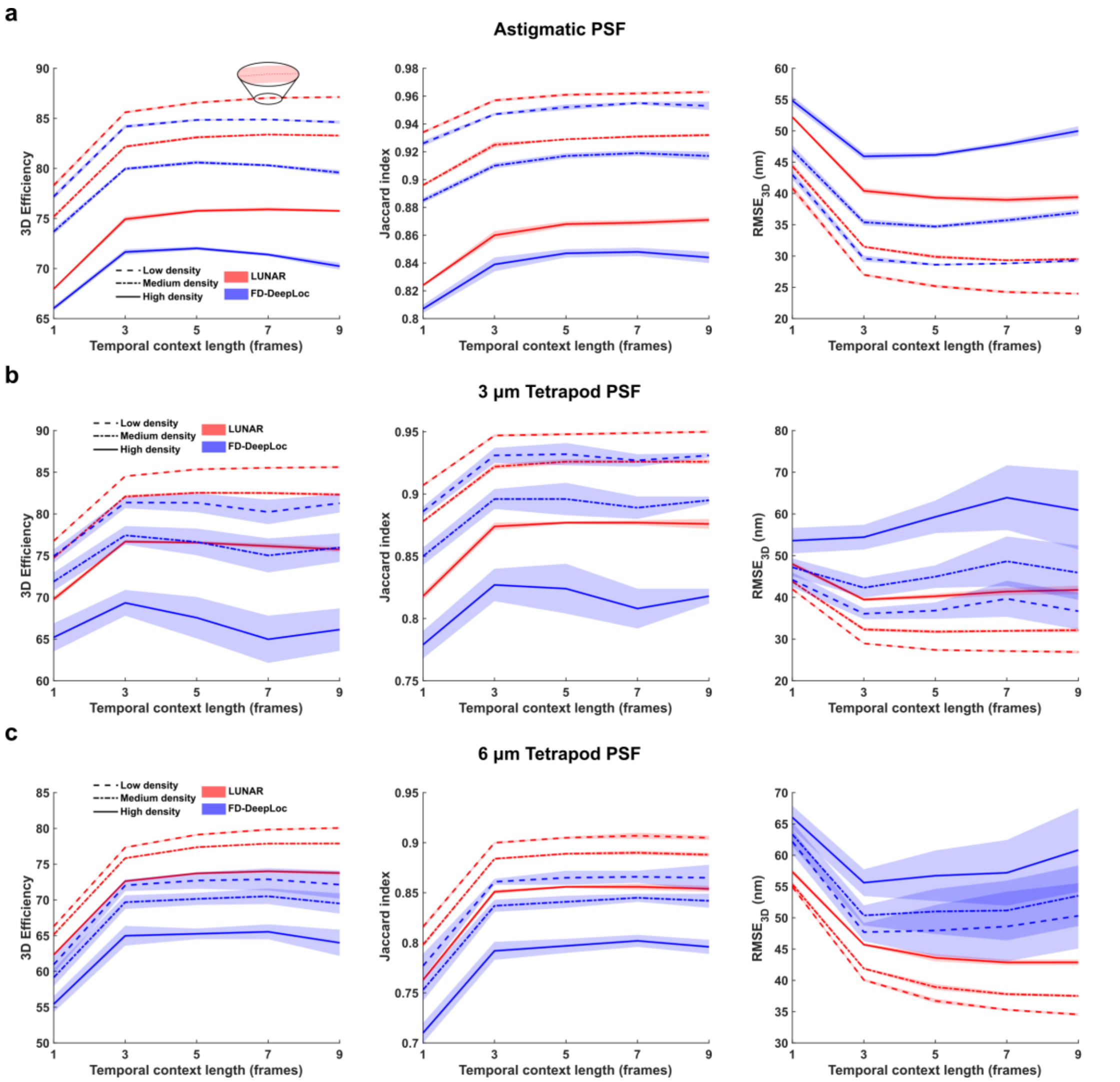


**Supplementary Fig. 12 | Performance of LUNAR and FD-DeepLoc trained with varying temporal context lengths on simulated microtubules. a**, From left to right: plots showing 3D efficiency, Jaccard index, and localization error across different temporal context lengths, evaluated on the astigmatic PSF based microtubule dataset. **b** and **c**, Same as **a**, but evaluated on the 3 μm Tetrapod PSF and 6 μm Tetrapod PSF based datasets, respectively. Both LUNAR and FD-DeepLoc were trained using ground truth PSF, estimated SNRs derived from the raw data (photon range [1000, 10000], background range [33, 66] with Perlin noise), and fixed emitter densities (0.24 emitters/μm$^2$ for astigmatic PSF, 0.06 emitters/μm$^2$ for Tetrapod PSF. The models were then tested across all densities with the same PSF type. Solid lines and shaded areas indicate mean and standard deviation of 5 repeated experiments.

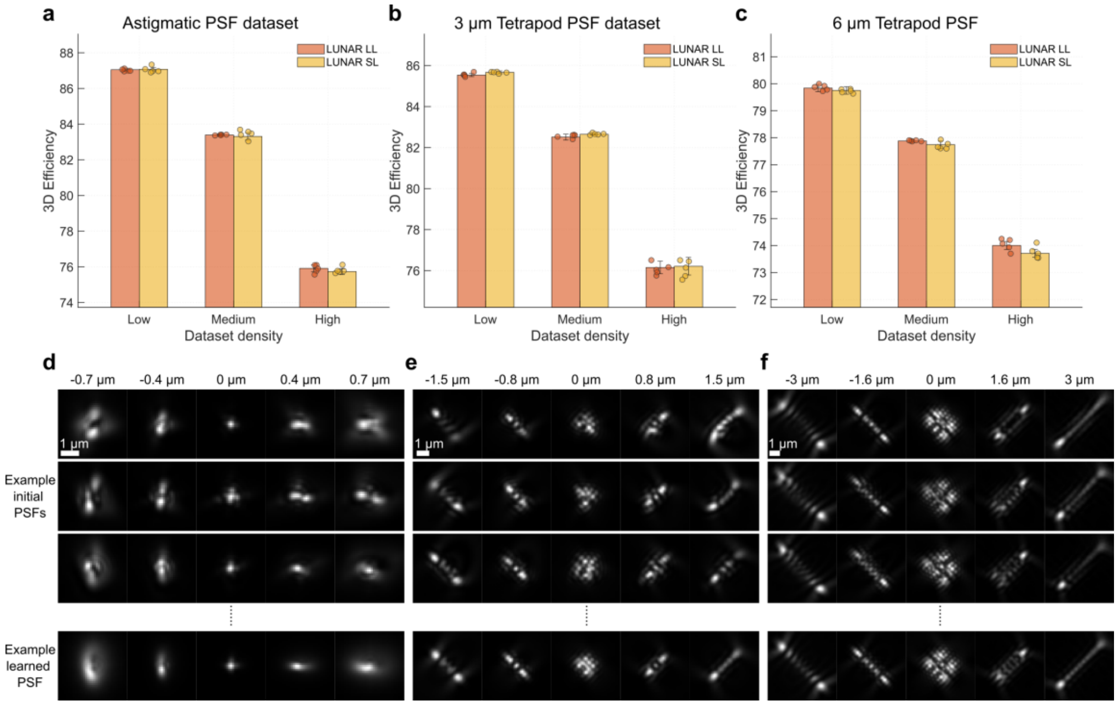


**Supplementary Fig. 13 | LUNAR exhibits stable performance for unknown PSF model. a**, Average 3D efficiency of LUNAR LL (trained with ground truth PSF) and LUNAR SL (trained with random initial PSFs) on simulated microtubule datasets with astigmatic PSF. Amplitude of each aberration was initialized by adding zero mean Gaussian noise with a standard deviation of λ/50 to the ground truth. **b** and **c**, Same as **a**, but evaluated on the 3 μm Tetrapod PSF and 6 μm Tetrapod PSF, respectively. **d**, Example randomly initialized astigmatic PSFs and the learned PSF of LUNAR SL. **e** and **f**, Same as **d**, but for 3 μm Tetrapod PSF and 6 μm Tetrapod PSF, respectively. Error bars indicate the standard deviation of 5 repeated experiments. Scale bars, 1 μm (**d**, **e**, **f**).

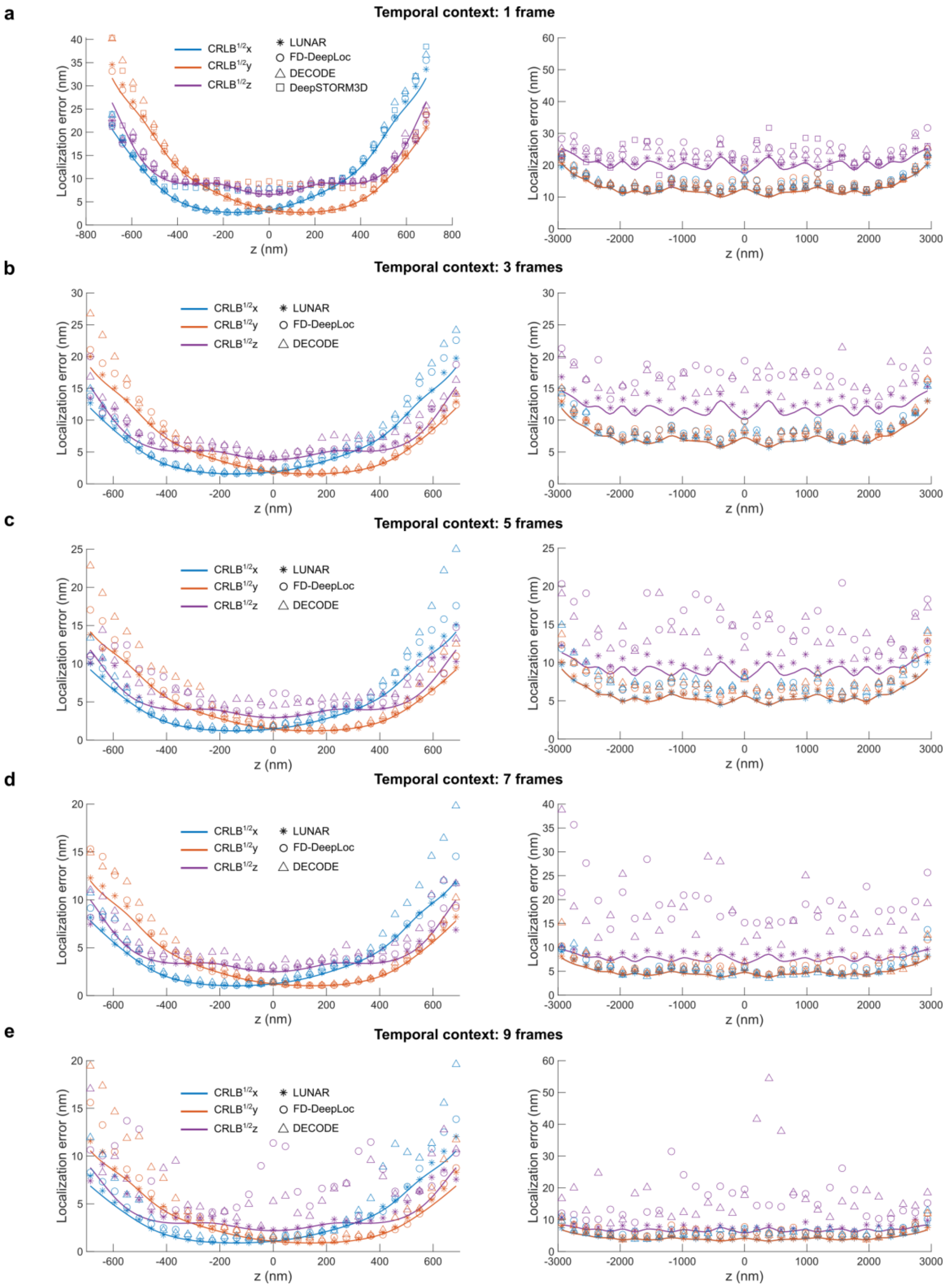


**Supplementary Fig. 14 | Evaluation of the localization accuracy using different lengths of temporal context. a**, Localization error at different z-positions analyzed by networks using no temporal context (1 frame), with the information limit (CRLB) shown for comparison. Left: astigmatic PSF. Right: 6 µm Tetrapod PSF. **b-e**, the same as **a**, but using temporal contexts of 3-, 5-, 7-, and 9-frames, respectively.

DeepSTORM3D was only evaluated with 1-frame input, as it is not designed to utilize temporal context. Single molecules were simulated with a fixed photon count of 5,500 and background of 50 photons per frame. For each of the 31 z-positions investigated, 3,000 frames were simulated. All networks except DeepSTORM3D were trained over a photon range of [1,000, 10,000] and a background range of [40, 60]. DeepSTORM3D was trained using a narrower photon range of [4,000, 7,000] to ensure stable convergence, and its normalization strategy was modified to improve generalization between training and test data. Training densities were set to 0.24 emitters/$\mu m^2$ for astigmatic PSF and 0.06 emitters/$\mu m^2$ for Tetrapod PSF.

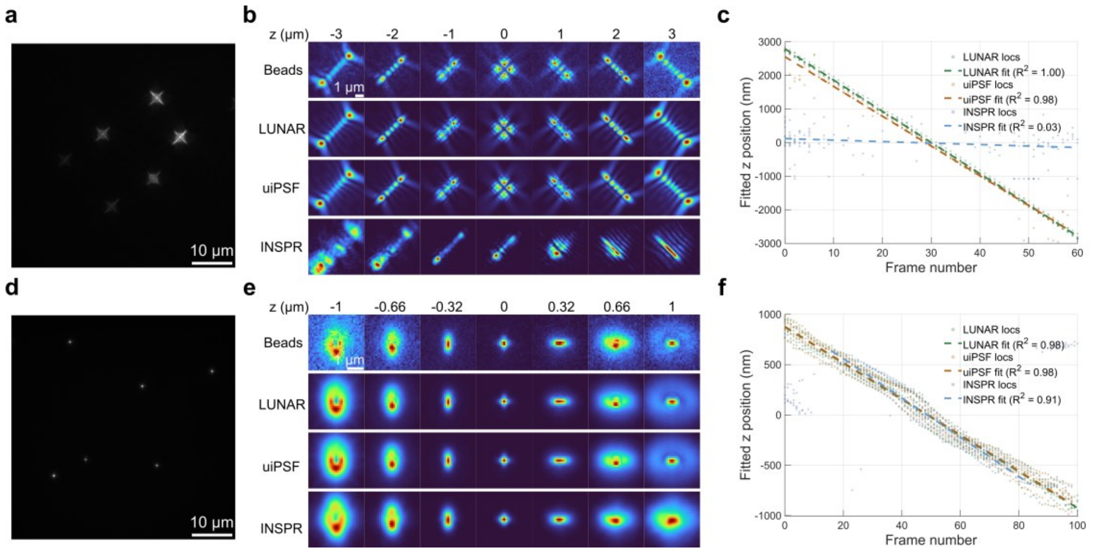


**Supplementary Fig. 15 | In situ PSF learning from sparse bead data. a**, Sum projection of a 6 μm Tetrapod PSF bead stack. **b**, Example learned PSFs at different axial positions using LUNAR, uiPSF, and INSPR, without prior knowledge of objective stage positions. LUNAR and uiPSF are based on a vectorial PSF model while INSPR is based on a scalar PSF model. **c**, Estimated z positions of beads using PSF models learned by different algorithms. Bead datasets were acquired by capturing images at each step of the objective stage movement. $R^2$ values indicate the goodness of fit of all localizations to a linear function. **d-f**, Same as **a-c** but for astigmatic PSF. For the Tetrapod PSF data, LUNAR, uiPSF, and INSPR detected 369, 209, and 139 emitters, respectively. For the astigmatic PSF data, LUNAR, uiPSF, and INSPR detected 594, 593, and 411 emitters, respectively. Scale bars, 10 μm (**a**, **d**), 1 μm (**b**, **e**).

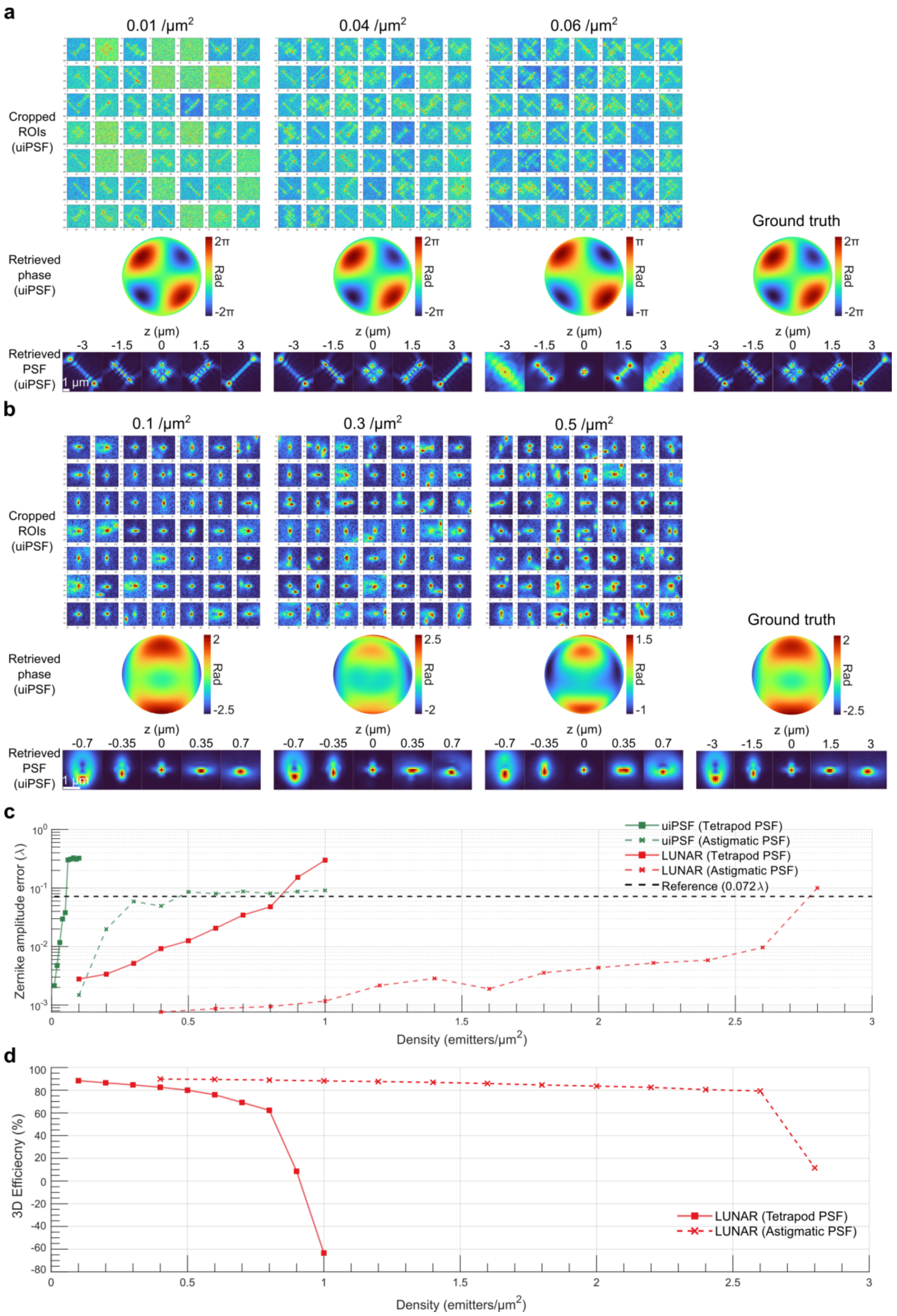


**Supplementary Fig. 16 | Performance of in situ PSF learning across varying emitter densities. a**, Example results for the 6 μm Tetrapod PSF at different emitter densities. From top to bottom: cropped regions of interest (ROIs), retrieved pupil phase, and reconstructed PSF using uiPSF. From left to right: increasing emitter density and corresponding ground truth. **b**, Same as **a**, but for the astigmatic PSF. **c**, Zernike amplitude estimation error as a function of increasing emitter density for in situ PSF learning using uiPSF and LUNAR, for both the 6 μm Tetrapod and astigmatic PSFs. **d**, 3D efficiency as a function of increasing emitter density for LUNAR with in situ PSF learning. Scale bars, 1 μm (**a**, **b**).

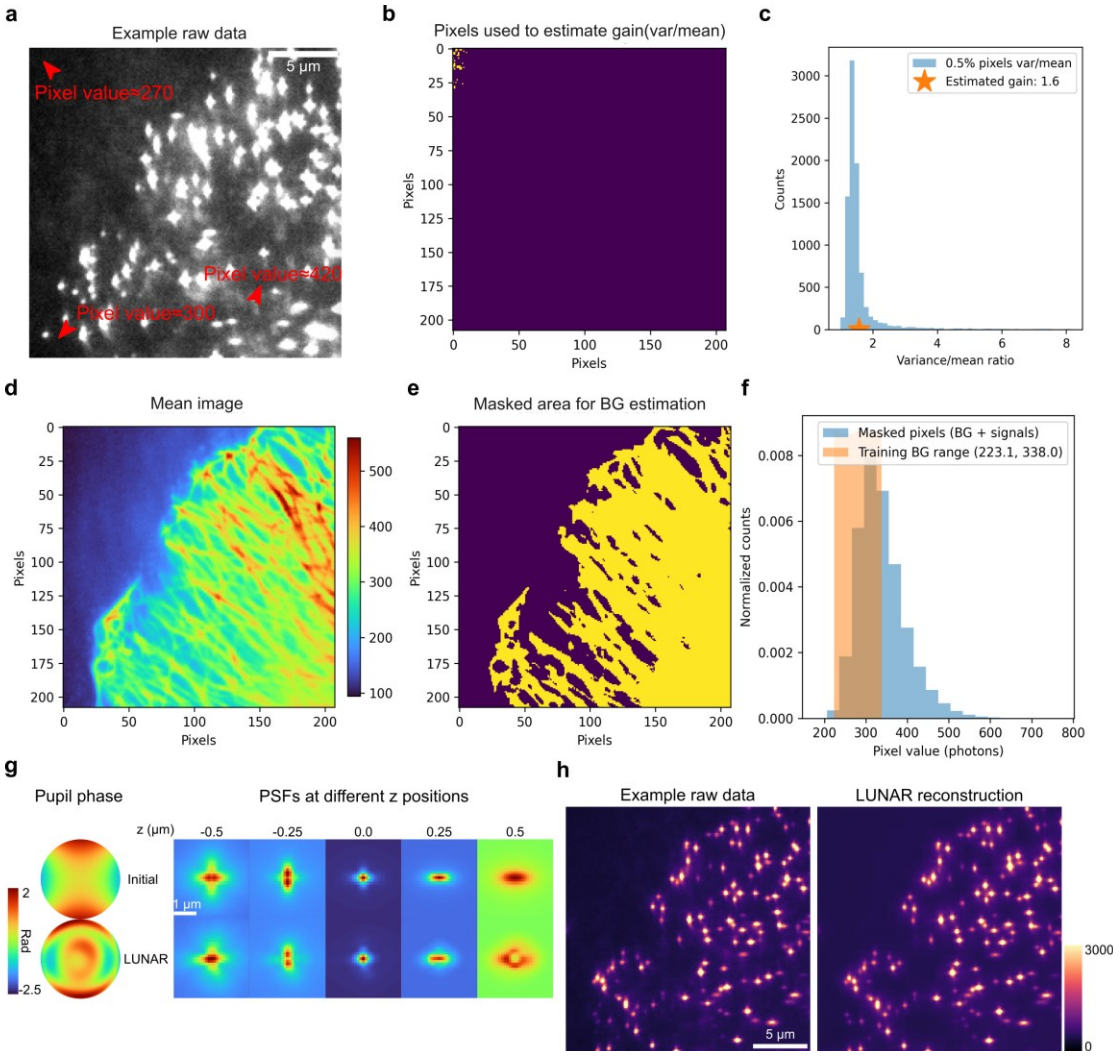


**Supplementary Fig. 17 | Training background range estimation from microtubule dataset and LUNAR learned representations. a**, Example microtubule data labeled using DNA-PAINT, showing a highly non-uniform background. **b** and **c**, An optional step to calibrate camera gain using the raw data. Briefly, the lowest 0.5% of pixel values (**b**) are selected, which are assumed to represent background regions with constant photon counts over short time intervals. These pixel values are then converted to photon units using provided camera parameters. **c**, The gain is estimated by computing the ratio of variance to mean across these pixels within short time windows. Under Poisson statistics, this ratio should equal one, so deviations are used to calibrate the gain accordingly. **d**, Average intensity projection of the microtubule dataset. **e**, Due to the non-uniform background, regions containing biological structures exhibit higher background, so we identified sample-containing areas and used them to estimate the background range for network training (**f**). **g**, Comparison between the initial PSF provided for training and the learned PSF by LUNAR. **h,** Example raw data and LUNAR's reconstruction using the learned PSF and predicted localizations. Scale bars, 5 μm (**a**, **h**), 1 μm (**g**).

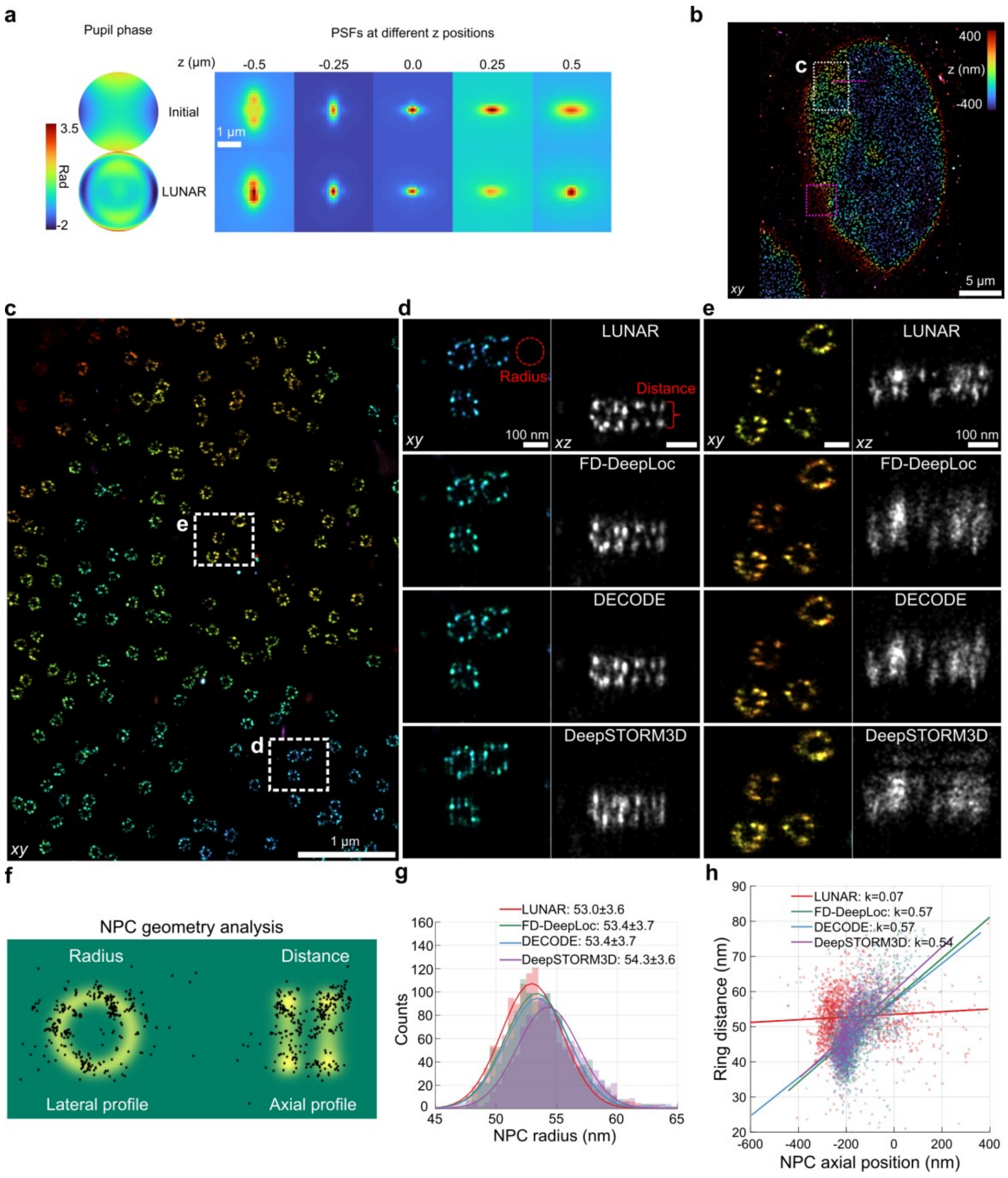


**Supplementary Fig. 18 | Super-resolution imaging of nuclear pore complexes (NPCs) using different algorithms. a,** Comparison between the initial PSF provided for training and the learned PSF by LUNAR. **b**, Overview of the super-resolution image of NPCs (Nup96-SNAP, AF647, U2OS cells) reconstructed by LUNAR. Regions indicated by magenta dashed box and line are showed in **Fig. 3e,f** in main text. **c**, Magnified views of the region denoted by the dashed box **c** in **b**. **d** and **e**, Magnified views of the regions denoted by dashed boxes in **c**, comparing top- and side-view reconstructions by different algorithms (LUNAR, FD-DeepLoc, DECODE and

DeepSTORM3D). **f**, Schematic of NPC geometry analysis. Radius and distance are fitted by a double-ring model. **g**, Histograms of fitted NPC radii based on reconstructions of different algorithms. Reported values are mean ± s.d., computed based on $n_{Method}$ sites extracted from each method's reconstruction. Sample size: $n_{LUNAR}$=1558, $n_{FD\text{-}DeepLoc}$=1449, $n_{DECODE}$=1412, $n_{DeepSTORM3D}$=1251 NPCs. Similar results were overserved in 3 independent experiments. **h**, Double-ring distance of Nup96 as a function of NPC axial position. Straight lines represent linear regression fits for each algorithm, with *k* denoting the corresponding slope. A larger *k* reflects a stronger correlation between ring distance and NPC axial position, suggesting greater distortion due to PSF mismatch. Scale bars, 5 μm (**b**), 1 μm (**a**, **c**), 100 nm (**d**, **e**).

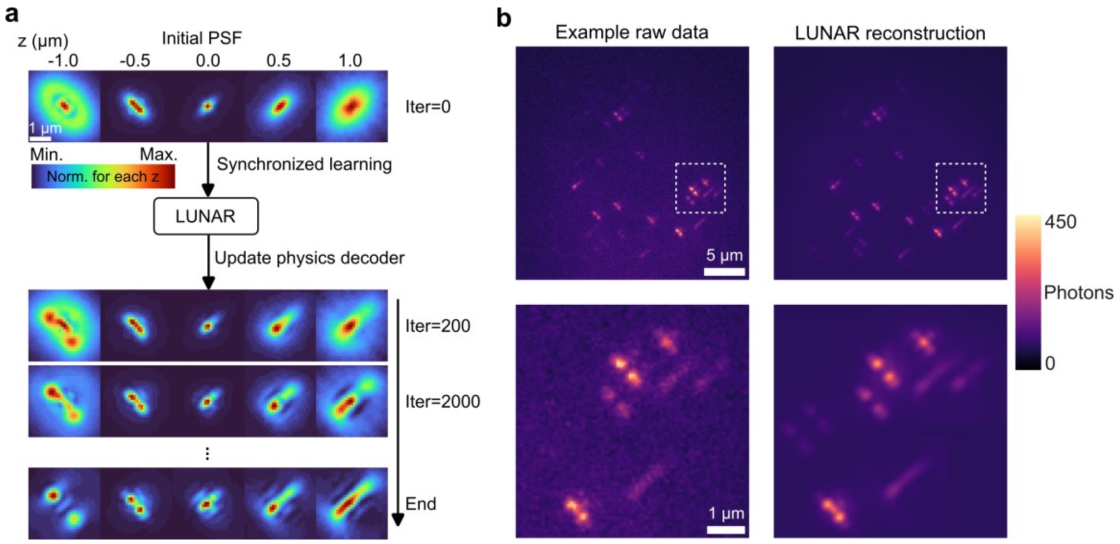


**Supplementary Fig. 19 | LUNAR enables cross-type PSF correction. a**, LUNAR's synchronized learning strategy dynamically adapts the physics decoder during training, transitioning from an initial astigmatic PSF to the DMO saddle-point PSF. **b**, Comparison between experimental single-molecule data (left) and data reconstructed using the learned PSF model and LUNAR predictions (right). Scale bars, 5 μm (**b** top), 1 μm (**a**, **b** bottom).

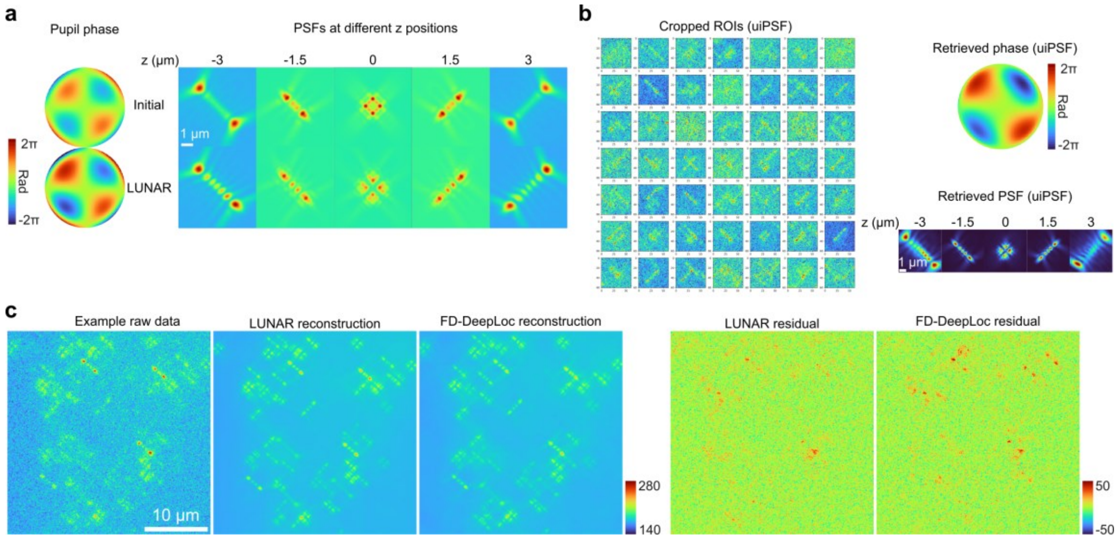


**Supplementary Fig. 20 | Learned representations from whole-cell datasets acquired with a 6 μm DMO Tetrapod PSF. a**, Comparison between the initial PSF used for training and the learned PSF by LUNAR on the NPC dataset. **b**, PSF learned by uiPSF on the NPC dataset. **c**, Example raw data of the mitochondria dataset and corresponding reconstructions by LUNAR localizations and LUNAR-learned PSF, or FD-DeepLoc localizations and bead PSF. Scale bars, 10 μm (**c**), 1 μm (**a**, **b**).

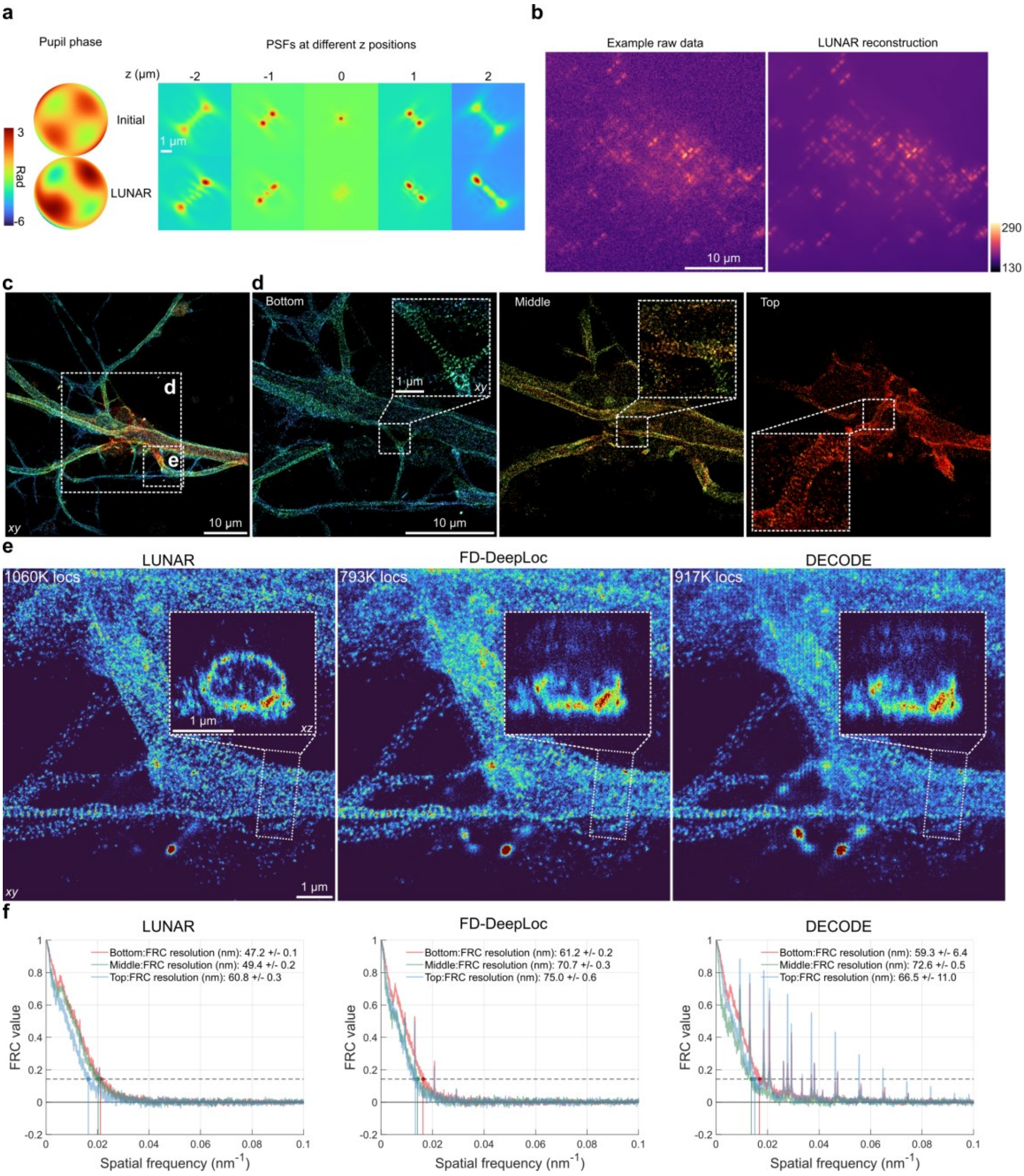


**Supplementary Fig. 21 | Super-resolution imaging of neurons using different algorithms. a**, Comparison between the initial PSF used for training and the learned PSF by LUNAR. **b**, Example raw SMLM data and corresponding reconstructions using LUNAR localizations and LUNAR-learned PSF. **c**, Overview of the super-resolution image of neurons (βII-spectrin, AF647, mESC-derived neuronal cells) reconstructed by LUNAR. **d**, Super-resolution images reconstructed using bottom (-2 to 0 μm), middle (0 to 0.8 μm), and top (0.8 to 2 μm) axial sections of the region denoted by the dashed box **d** in **c**. **e**, Magnified views of the region denoted by the dashed box **e** in **c**, reconstructed by LUNAR, FD-DeepLoc, and DECODE. Images are color-coded by localization density, with number of localizations annotated in the top left. **f**, FRC resolution analysis of the different axial sections shown in **d**. Scale bars, 10 μm (**b**, **c**, **d**), 1 μm (**d** insets, **e**).

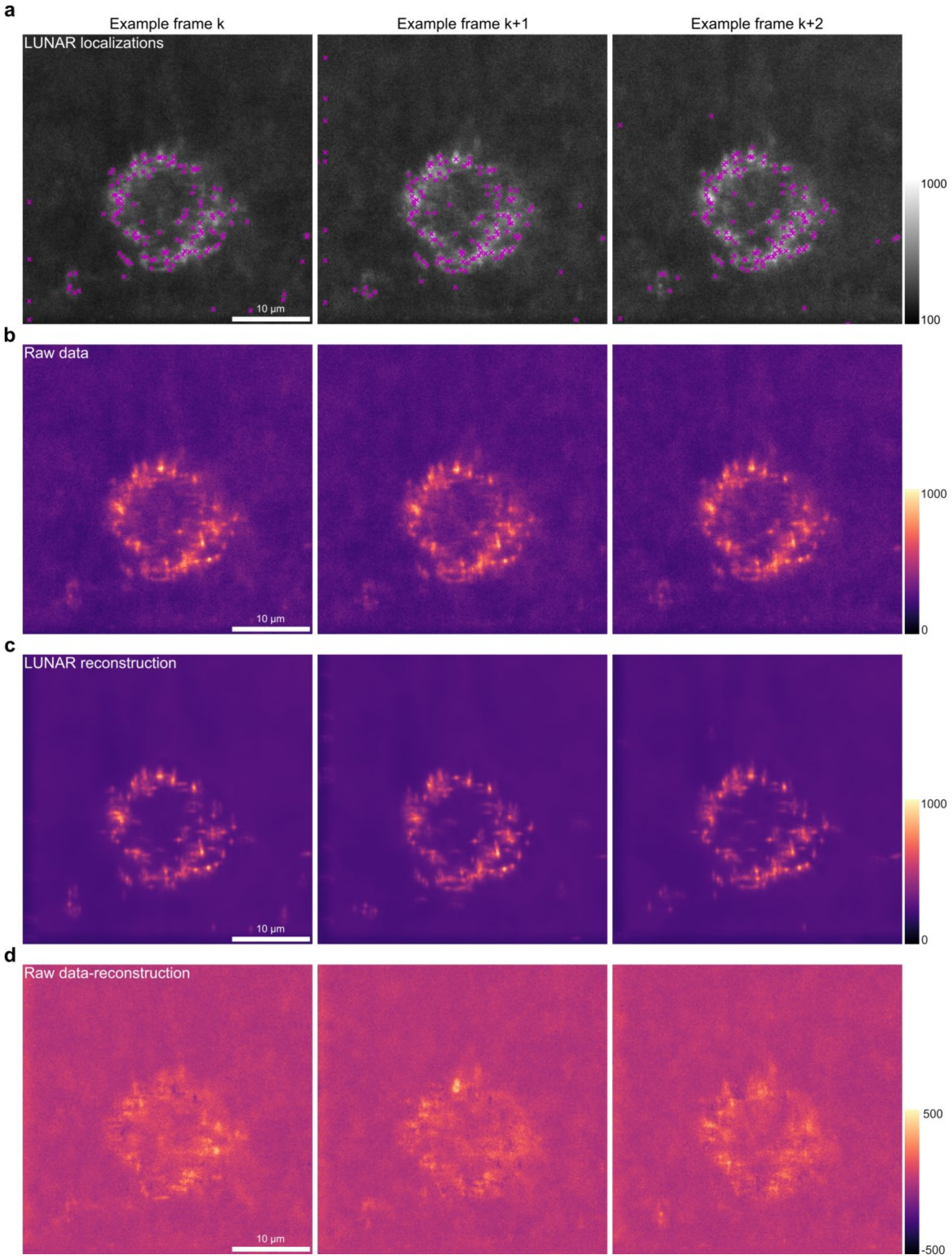


**Supplementary Fig. 22 | Example frames from the lattice light-sheet motor-PAINT dataset. a**, LUNAR localizations (magenta crosses) overlaid on the example raw frames. **b-d**, Example images for the raw data (**b**), corresponding reconstructions using LUNAR localizations and pre-calibrated PSF (**c**), and residual error maps (**d**). Scale bars, 10 µm (**a**, **b**, **c**, **d**).

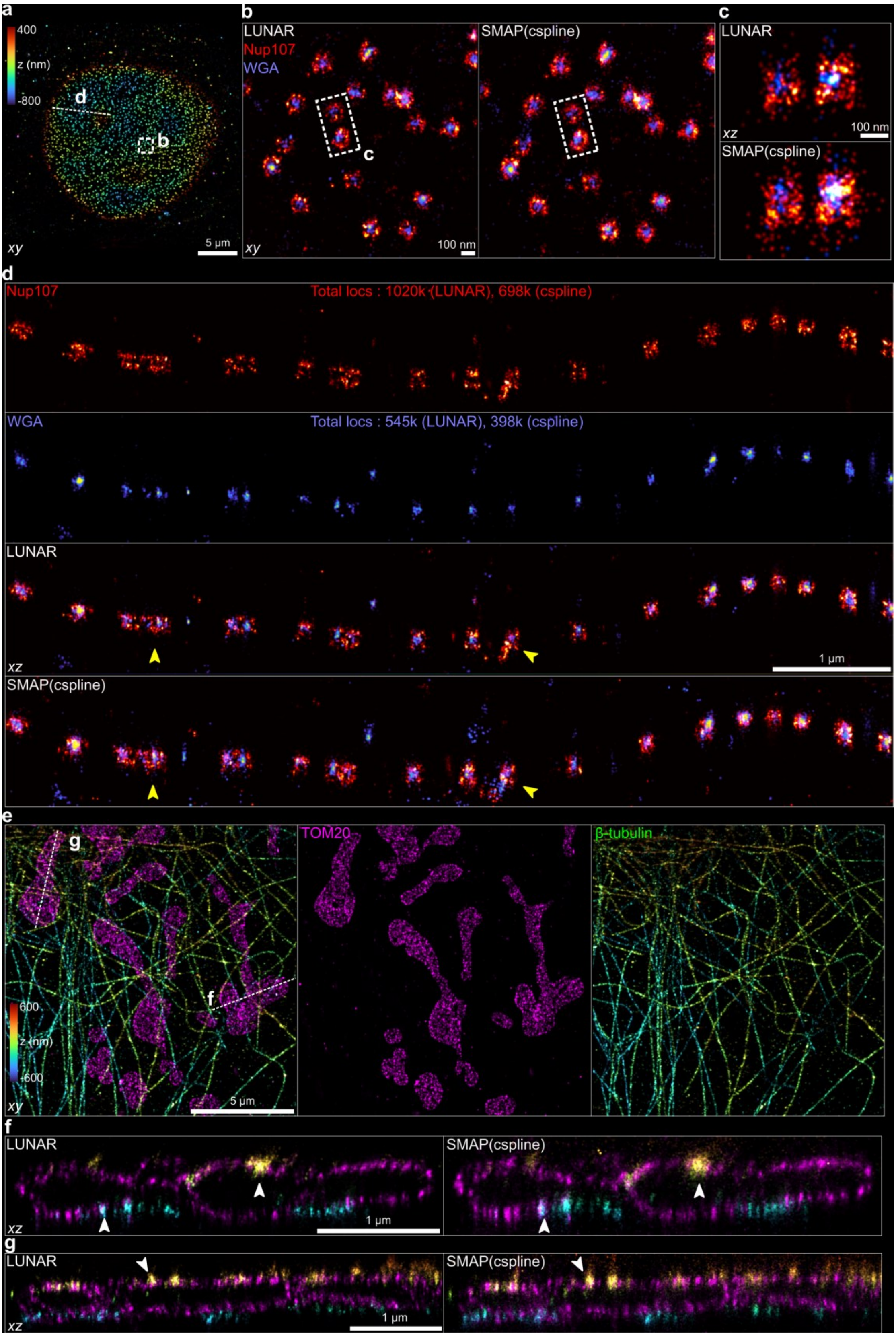


**Supplementary Fig. 23 | Dual-color super-resolution imaging using LUNAR. a,** Overview of a dual-color super-resolution image of NPCs (Nup107-SNAP-AF647, WGA-CF680, U2OS cells) reconstructed by LUNAR. The image is color-coded by z-position. **b**, Magnified views of the area denoted by the dashed box in **a**, reconstructed using LUNAR and cspline. **c**, Side-view reconstructions of the area denoted by the dashed box in **b**. **d**, 500 nm-width side-view reconstructions along the dashed line in **a**. From top to bottom are: LUNAR-reconstructed Nup107, WGA, the merged image, and the cspline-reconstructed merged image. Yellow arrows indicate structural details better resolved by LUNAR. **e**, Overview of a dual-color super-resolution image of mitochondria and microtubules reconstructed by LUNAR (Tom20-AF647, β-tubulin-CF680, COS-7 cells). **f** and **g**, 500 nm-width side-view reconstructions along the dashed lines in **e**, reconstructed by LUNAR and cspline, respectively. While arrows indicate the mitochondria-microtubule contacts. Scale bars, 5 μm (**a**, **e**), 1 μm (**d**, **f**, **g**), 100 nm (**b**, **c**).

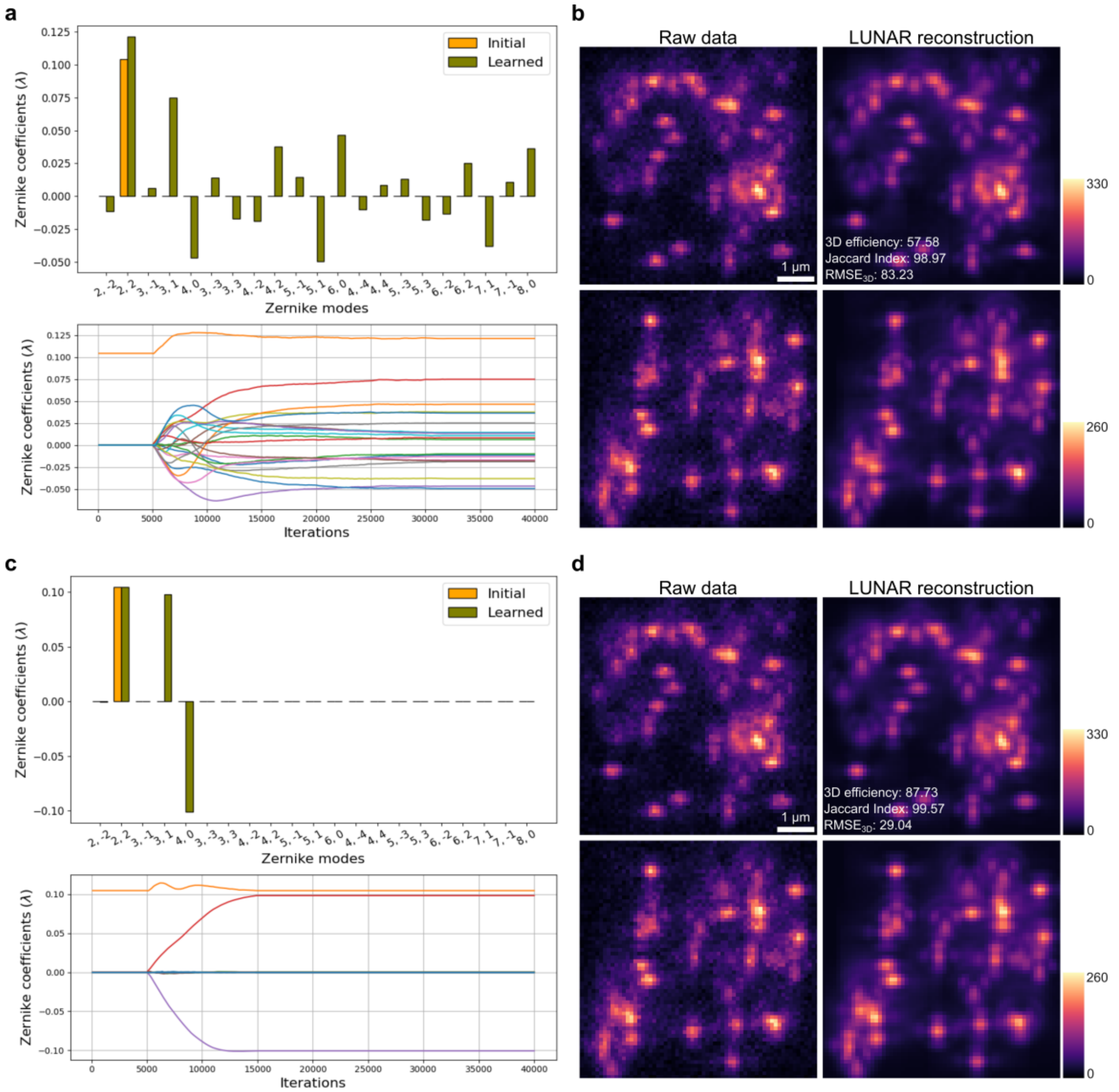


**Supplementary Fig. 24 | Example convergence to a local minimum. a**, Initial and learned Zernike coefficients for LUNAR applied to an astigmatic PSF dataset (1.2 emitters/μm$^2$, 0.1λ extra aberrations). A total of 21 Zernike coefficients were optimized during training. **b**, Example raw data frames and corresponding reconstructions by LUNAR. The model converges to a local minimum that closely reproduces the observed data. This leads to a performance drop observed in **Supplementary Fig. 6b**. **c** and **d**, Same as **a** and **b**, respectively, but only 7 Zernike coefficients were optimized during training. This strategy can help reduce ill-posedness, as practical aberrations are usually low-order. Scale bars, 1 μm (**b**, **d**).

### Supplementary Tables

| Main functions | Name | PSF learning on sparse data | PSF learning on overlapped data | Localization on overlapped data | Use temporal information | Reach temporal CLRB | Network architecture | Principle |
|---|---|---|---|---|---|---|---|---|
| Deep learning localization + PSF learning | LUNAR (this work) | **Yes** | **Yes** | **Yes** | **Yes** | **Yes** | CNN+ Transformer | Self-supervised neural-physics learning |
| Deep learning localization methods | DECODE (Nat Methods, 2021) | No | No | **Yes** | **Yes** | No | CNN | PSF-supervised simulation learning |
| | DeepSTORM3D (Nat Methods, 2020) | No | No | **Yes** | No | Not applicable | CNN | PSF-supervised simulation learning |
| | FD-DeepLoc (Nat Methods, 2023) | No | No | **Yes** | **Yes** | No | CNN | PSF-supervised simulation learning |
| In situ PSF retrieval methods | INSPR (Nat Methods, 2020) | **Yes** | No | No | No | Not applicable | Not applicable | Model based optimization |
| | uiPSF (Nat Methods, 2024) | **Yes** | No | No | No | Not applicable | Not applicable | Model based optimization |

**Supplementary Table 1 | Distinctions between LUNAR and related methods.**

**Supplementary Notes**

#### Supplementary Note 1 The development of the LUNAR

##### 1.1 Single-emitter fitting

For conventional SMLM algorithms, the localization process can be described as a parameter estimation problem. Specifically, it involves finding the emitter parameters $h$ that maximize the log likelihood of the data:

$$\underset{h}{\operatorname{argmax}} \log p_\theta(d|h) \tag{S1}$$

where $h$ corresponds to the emitter position and brightness, $d$ is the collected data of an emitter, $\theta$ represents the PSF parameters involved in the probability computation, which are typically determined in advance through bead calibration. For the most common Poisson noise in SMLM, the likelihood is given by:

$$p_\theta(d|h) = \prod_k \frac{e^{-\mu_k(h,\theta)} \mu_k(h,\theta)^{d_k}}{d_k!} \tag{S2}$$

Here, $\mu_k(h,\theta)$ is the expected number of photons in pixel $k$ from the PSF model function, $d_k$ is the measured number of photons. Normally, the execution of equation (S1) assumes that the fitted data $d$ is generated by a single emitter. Therefore, $\mu_k(h,\theta)$ is calculated by the integration of a single PSF model over the pixels:

$$\mu_k(h,\theta) = \iint_{\text{pixel}_k} \left(h_{phot} \cdot \text{PSF}(h,\theta) + bg\right) dxdy \tag{S3}$$

where $h_{phot}$ is the brightness, $bg$ is the background density. The PSF can be constructed in flexible forms, such as Gaussian functions[1], spline functions[2], and diffraction models[3]. In this work, we applied the vectorial PSF model (**Supplementary Note 2**) for its high accuracy and physical interpretability.

##### 1.2 Multiple-emitter fitting

The single-emitter assumption in conventional SMLM algorithms necessitates that blinking signals be cropped sparsely and fitted one by one, often accelerated by parallel computation. This assumption holds true in most SMLM experiments with well-controlled fluorescence properties and laser activation conditions. However, as SMLM requirements have evolved to include faster imaging or large depth-of-field (DOF) imaging, the signals in the raw data often overlap with each other. This overlap means that the number of PSF models required for the fitting process is unknown, posing significant challenges to constructing the correct model for fit:

$$\mu_k(h,\theta) = \iint_{\text{pixel}_k} \left(\sum_{n=1}^{N} h_{phot}^{n} \cdot \text{PSF}(h^n,\theta) + bg\right) dxdy \tag{S4}$$

Here, $N$ is the total number of emitters, and every pixel in the model receives photons from every potential emitter. As $N$ is unknown, it becomes hard to find the optimal solution for equation (S1).

To address this problem, multiple-emitter fitting algorithms have been developed. Early approaches generally involve fitting using several models with different number of emitters and then selecting the best one using specific criteria[4,5]. This is somewhat similar to maximum a posteriori (MAP) method. Recently, Bayesian methods using Markov Chain Monte Carlo (MCMC) have been proposed to solve multiple-emitter localization[6] and tracking problems[7]. These methods offer a principled way to handle the uncertainty in the number of emitters and their positions by estimating the posterior:

$$p_\theta(h|d) = \frac{p_\theta(d,h)}{p_\theta(d)} = \frac{p_\theta(d|h)p_\theta(h)}{p_\theta(d)} \tag{S5}$$

Here, $p_\theta(h|d)$ is the posterior distribution of all emitters' parameters $h$ given the data $d$, without sparsity requirement. It is proportional to the product of the likelihood $p_\theta(d|h)$ and priors $p_\theta(h)$. $p_\theta(d)$ is called the evidence. Deterministic estimations about the parameters $h$ could be obtained by investigating the posterior. However, Bayesian methods often suffer from significant computational burdens due to the large number of parameter sampling and rejection operations required. Additionally, the need for substantial mathematical expertise to properly implement and interpret these methods has hindered their widespread adoption.

Another approach to multiple-emitter localization is the use of deep neural networks[8,9], which combine the advantages of efficiency and accuracy through a high-quality training process. Therefore, we propose LUNAR localization learning (LUNAR LL), which utilizes a network parameterized by $\phi$ to approximate the multiple-emitter posterior estimation function. This is achieved by:

$$\underset{\phi}{\operatorname{argmin}}\ \mathrm{E}_{p_\theta(d)}\left[D_{\mathrm{KL}}\left(p_\theta(h|d), q_\phi(h|d)\right)\right] \tag{S6}$$

Here, $\phi$ is updated to minimize the Kullback-Leibler (KL) divergence between the posteriors under the PSF model parameterized by $\theta$ and the inference network. As the PSF is known and the image formation process is well understood, we can easily sample the generative model $d, h \sim p_\theta(d, h)$ to generate the training data pairs with different densities. As derived in **Methods**, equation (S6) is equivalent to minimizing the negative log likelihood $-\log q_\phi(h|d)$, which can be decomposed into three terms:

$$-\log q_\phi(h|d) = L_{\mathrm{ce}} + L_{\mathrm{count}} + L_{\mathrm{loc}} \tag{S7}$$

where:

$$L_{\mathrm{ce}} = \sum_k -[p_k \times \log(\hat{p}_k) + (1 - p_k) \times \log(1 - \hat{p}_k)] \tag{S8}$$

$$L_{\mathrm{count}} = \frac{(N - \mu_{\mathrm{count}})^2}{2\sigma_{\mathrm{count}}{}^2} + \log\left(\sqrt{2\pi}\sigma_{\mathrm{count}}\right) \tag{S9}$$

$$L_{\mathrm{loc}} = -\frac{1}{N}\sum_{n=1}^{N} \ln\left(\sum_{k=1}^{K} \frac{\hat{p}_k}{\sum_k \hat{p}_k} \cdot p_{\mathrm{Gauss}}(\mathbf{gt}_n; \boldsymbol{\mu}_k, \boldsymbol{\Sigma}_k)\right) \tag{S10}$$

Here, $L_{\mathrm{ce}}$ is the negative log likelihood of a Bernoulli distribution, where $\hat{p}_k$ represents the precited probability that an emitter exists in the pixel $k$, and $p_k$ is the binary label. $L_{\mathrm{count}}$ is the negative log likelihood of the true emitter number $N$ under the approximate Gaussian distribution of the emitter number, with mean and variance computed by $\mu_{\mathrm{count}} = \sum_k \hat{p}_k$ , $\sigma_{\mathrm{count}}{}^2 = \sum_k \hat{p}_k (1 - \hat{p}_k)$ . $L_{\mathrm{loc}}$ is the negative log likelihood of the true emitter parameters $\mathbf{gt}_n = [x_n, y_n, z_n, phot_n]$ under the predicted Gaussian-mixture model (GMM), with each component being a four-dimensional Gaussian distribution with normalized weight $\hat{p}_k / \sum_k \hat{p}_k$. $\boldsymbol{\mu}_k$ are the predicted means $[\hat{x}_k, \hat{y}_k, \hat{z}_k, \widehat{phot}_k]$ for positions and brightnesses, and $\boldsymbol{\Sigma}_k$ is the diagonal covariance matrix $\mathrm{diag}(\sigma^2_{x,k}, \sigma^2_{y,k}, \sigma^2_{z,k}, \sigma^2_{phot,k})$ representing the uncertainties in the predictions. In addition to predicting the complex parameter distributions using equation (S7), a simple $L_{\mathrm{bg}}$ term is introduced to estimate the background without uncertainty (**Methods**).

During the training process, the inference network gradually approximates a variational function that can estimate the posterior given data with an unknown number of emitters. When the number of emitters is one, the inference network should give similar results to equations (S1-S3), which has been shown to reach the theoretical minimum uncertainty (Cramér-Rao Lower Bound, CRLB). The single-molecule localization accuracy test verifies that LUNAR can also reach the CRLB, even with a longer temporal context and without manually linking signals that persist over multiple frames (**Supplementary Fig. 12**).

##### 1.3 Multiple-emitter fitting with PSF learning

Until now, whether in the single-emitter case or the multiple-emitter case, the PSF parameters $\theta$ are assumed to be known, meaning we have an explicit form of $p_\theta(d|h)$ to solve equation (S1) or equation (S6). However, when imaging deep inside the samples or with poor PSF calibration, the understanding of $p_\theta(d|h)$ may be incorrect, leading to errors with previous methods. To address this issue, existing solutions have been proposed to estimate the in-situ PSF from the abundant SMLM data[10,11]. The idea is feasible because the blinking signals of single molecules provide random views of the 3D PSF at different axial positions. Despite their randomly distributed positions, these signals inherently carry information about the shared wavefront distortion, which is parameterized by $\theta$. This naturally leads to maximizing the likelihood of the data $p_\theta(d|h)$ by optimizing the global parameters $\theta$ of the PSF:

$$\underset{\theta,h}{\operatorname{argmax}} \sum_{n=1}^{N} \log p_\theta(d^n|h^n) \tag{S11}$$

Here, $d^n$ and $h^n$ represent the measured data and local parameters of the $n$-th emitter, respectively. Typically, thousands of emitters at different axial positions are sufficient to estimate $\theta$. Notably, this method still requires that the raw data be sparse enough, as $p_\theta(d^n|h^n)$ is the likelihood function for a single-PSF model. However, in densely labeled samples or when using large-DOF PSF engineering with large PSF size (**Supplementary Fig. 4**), both isolating individual emitters and determining the number of emitters become difficult. It is challenging to further estimate PSF parameters with the overlapped single-molecule data.

To address the above challenges, we draw inspiration from the framework of the variational autoencoder[12] (VAE), which implicitly maximizes the likelihood of the observed data (without the sparsity assumption) by jointly learning an encoder $q_\phi(h|d)$ and a decoder $p_\theta(d|h)$. SMLM experiments can be viewed as a random process involving the unobserved variables $h$, which represent the parameters of multiple molecules. At specific time points, these molecules are first activated according to the prior $p_\theta(h)$, considering both the specimen and fluorophore properties. Then, these activated molecules are observed by the optical system, with the likelihood function $p_\theta(d|h)$ describing the PSF and the stochastic shot noise:

$$p_\theta(d,h) = p_\theta(d|h)p_\theta(h) \tag{S12}$$

The marginal data likelihood is given by:

$$p_\theta(d) = \int p_\theta(d|h)p_\theta(h)\,dh \tag{S13}$$

Since the above integral is intractable and cannot be analytically calculated, we turn to maximizing the evidence lower bound (ELBO) derived from Jensen's Inequality:

$$\begin{aligned}\log p_\theta(d) &= \log\int p_\theta(d,h)\frac{q_\phi(h|d)}{q_\phi(h|d)}dh \\ &= \log \mathrm{E}_{q_\phi(h|d)}\left[\frac{p_\theta(d,h)}{q_\phi(h|d)}\right] \\ &\geq \mathrm{E}_{q_\phi(h|d)}\left[\log\frac{p_\theta(d,h)}{q_\phi(h|d)}\right] = \mathrm{ELBO}\end{aligned} \tag{S14}$$

where $q_\phi(h|d)$ is an approximation to the intractable posterior $p_\theta(h|d)$, similar to LUNAR LL in the previous section. As $q_\phi(h|d)$ approaches $p_\theta(h|d)$, the lower bound becomes tighter. Instead of directly optimizing equation (S14) with respect to $\phi$ and $\theta$, different algorithms have been developed to optimize the encoder and decoder with less gradient variance[13].

In this work we draw inspirations from the reweighted wake-sleep algorithm[14] (RWS) to maximize the ELBO for its benefits in learning the network encoder. We proposed LUNAR synchronized learning (LUNAR SL), which integrates multiple-emitter fitting and PSF learning into one task and solves them simultaneously. Similar to the wake phase $p$-update and sleep phase $q$-update in RWS, LUNAR SL alternates between the physics learning and localization leaning (LUNAR LL) steps, with corresponding objectives and estimated gradients provided in **Methods**. From another perspective, LUNAR SL is also akin to the expectation-maximization[15] method, as it includes two key steps: (1) Calculate the expectation of the data likelihood under the posterior $p_{\theta^t}(h|d)$; (2) Maximize the expectation to update the generative parameters $\theta^{t+1}$. The former corresponds to training the network encoder based on the current PSF model and sampling the predicted distributions, while the latter corresponds to optimizing the PSF parameters.

#### Supplementary Note 2 Vectorial PSF model

To accurately describe the response of an optical system equipped with a high numerical aperture (NA) objective, a vectorial PSF model[3] that accounts for light polarization and refractive index mismatch is used in this work. As mentioned in the **Methods**, the model can be written as:

$$\mathrm{PSF} \propto \sum_{p=x,y} \sum_{d=x,y,z} \left|E_{p,d}^{\mathrm{img}}\right|^2, E_{p,d}^{\mathrm{img}} = \mathcal{F}_{2D}\left\{A(\rho,\varphi) E_{p,d}^{\mathrm{pupil}} e^{i\psi_{\mathrm{pos}}} e^{i\psi_{\mathrm{aber}}}\right\} \tag{S15}$$

where $A(\rho,\varphi) = \frac{(n_3{}^2-\rho^2NA^2)^{1/4}}{(n_1{}^2-\rho^2NA^2)^{1/2}}$ is the amplitude function with the apodization factor. $(\rho,\varphi)$ is the normalized polar coordinate in the pupil plane, with $\rho = 1$ corresponding to the limiting aperture angle $NA/n_3$. $NA$ is the numerical aperture, $n_1$, $n_2$ and $n_3$ are the refractive indices of the sample, cover glass and immersion oil, respectively. $\psi_{\mathrm{aber}}$ is the learnable phase term. $\psi_{\mathrm{pos}}$ is the position-dependent phase shift:

$$\psi_{\mathrm{pos}} = \frac{2\pi}{\lambda}\left( \begin{array}{c} NAx_0\rho\, cos\,\varphi + NAy_0\rho\, sin\,\varphi + \\ n_1 z_0 \sqrt{1-(\frac{\rho NA}{n_1})^2} - n_3 z_{\mathrm{nom}} \sqrt{1-(\frac{\rho NA}{n_3})^2} \end{array} \right) \tag{S16}$$

where $\lambda$ is the wavelength, $(x_0, y_0, z_0)$ is the molecule position, and $z_0$ represents the molecule's depth away from the cover glass. $z_{\mathrm{nom}}$ is the distance between the nominal focal plane and the cover glass. $E_{p,d}^{\mathrm{pupil}}$ represents the contribution of each fluorescent dipole component to the pupil field components. This can be calculated using the transformation matrix $\boldsymbol{M}$:

$$\boldsymbol{E}^{\mathrm{pupil}} = \boldsymbol{M} \cdot \boldsymbol{E}^{\mathrm{dipole}} \tag{S17}$$

Here, $\boldsymbol{E}^{\mathrm{dipole}}$ is the electric vector of the fluorescent dipole. Since most fluorescent probes often rotate freely, the dipole components are set with equal strength to mimic the random orientation case. To account for the effects of refractive index mismatch on different polarization components, $\boldsymbol{E}^{\mathrm{dipole}}$ is decomposed into p-polarized and s-polarized components and calculated separately, multiplied by their respective Fresnel transmission coefficients $t_p$ and $t_s$. Therefore, $\boldsymbol{M}$ is written as follows to include this process:

$$\boldsymbol{M}=\begin{bmatrix} t_p cos\theta_1 cos^2\varphi + t_s sin^2\varphi & cos\varphi sin\varphi(t_p cos\theta_1 - t_s) & -t_p sin\theta_1 cos\varphi \\ cos\varphi sin\varphi(t_p cos\theta_1 - t_s) & t_p cos\theta_1 sin^2\varphi + t_s cos^2\varphi & -t_p sin\theta_1 sin\varphi \\ 0 & 0 & 0 \end{bmatrix} \quad \text{(S18)}$$

The rows and columns of matrix $\boldsymbol{M}$ correspond to the pupil field components and the fluorescent dipole components, respectively. The third row is set to 0 as the z-component of the pupil field is negligible. The total Fresnel transmission coefficients for the p- and s-polarized light through multiple mediums are given by:

$$t_{k=p,s} = t_{k,1-2} \times t_{k,2-3} \quad \text{(S19)}$$

Taking the interface between the sample and the cover glass as an example, the Fresnel transmission coefficients are calculated as follows:

$$t_{p,1-2} = \frac{2n_1 \cos\theta_1}{n_1 \cos\theta_2 + n_2 \cos\theta_1}, t_{s,1-2} = \frac{2n_1 \cos\theta_1}{n_1 \cos\theta_1 + n_2 \cos\theta_2} \quad \text{(S20)}$$

For the numerical implementation of equation (S15), the Chirp *z*-transform is used to replace the 2D Fourier transform, allowing the decoupling of the relationship between sampling points in the imaging space and Fourier space. Optionally, an extra 2D Gaussian blurring can be applied to account for further system imperfections.

**Supplementary Note 3 Test datasets simulation**

**Random emitters for localization test**. To evaluate aberration robustness of localization, we simulated datasets of randomly distributed emitters with different densities (**Supplementary Fig. 4** and **5**). The simulations used an ideal camera model incorporating only Poisson noise, with image sizes of 64 × 64 pixels for astigmatic PSFs and 128 × 128 pixels for Tetrapod PSFs. A 5-pixel margin was left to avoid too many incomplete PSF patterns at image borders.

Since algorithms may leverage local temporal context to enhance localization accuracy, we modeled temporal blinking events using a simplified photophysical model adapted from DECODE[8], with an average on time of two frames. Specifically, for each emitter, the initial activation time was sampled from a continuous uniform distribution $\mathrm{U}(t_{start}, t_{end})$, and the on time followed an exponential distribution $\mathrm{E}(\theta)$ with $\theta = 2$. The signal intensity per blinking event in a single frame was sampled from a uniform distribution $\mathrm{U}(4000, 6000)$, reulsting in an average of 5000 photons. A constant background of 20 photons was added to each pixel.

In addition to the Zernike coefficients used to modulate the 3D PSF (astigmatic PSF: $C_2^2 = 70$ nm; 6 μm Tetrapod PSF: $C_2^{-2} = 230$ nm, $C_4^{-2} = -240$ nm, $C_6^{-2} = -17$ nm (wavelength $\lambda = 670$ nm)), we introduced additional coma ($C_3^1$) and spherical ($C_4^0$) aberrations to mimic unknown sample-induced aberrations at four levels: $0.025\lambda$ (16.75 nm), $0.05\lambda$ (33.5 nm), $0.075\lambda$ (50.25 nm), and $0.1\lambda$ (67 nm).

**Random emitters for in situ PSF learning test.** To investigate the maximum emitter density at which existing methods and LUNAR can accurately learn the in situ PSF, we simulated random emitter datasets using the same pipeline as described in the previous section. Briefly, we selected the 6 μm Tetrapod PSF and the astigmatic PSF with extra 0.05λ coma and spherical aberrations. We generated data across a wide range of emitter densities. For the Tetrapod PSF, we tested uiPSF at densities ranging from 0.01 to 0.1 emitters/μm², and LUNAR at higher densities from 0.1 to 1.0 emitters/μm². For the astigmatic PSF, uiPSF was evaluated from 0.1 to 1.0 emitters/μm², while LUNAR was tested from 0.4 to 2.4 emitters/μm². These ranges were chosen to systematically assess the breakdown point of each method, which was defined as the density at which the

Euclidean distance between the estimated and ground truth Zernike amplitude vectors exceeded 0.072λ[3].

**Simulated microtubules**. To quantitatively evaluate LUNAR's generalization across different PSFs and imaging conditions, we simulated a series of SMLM datasets of microtubules based on three PSFs: astigmatic PSF, 3 μm Tetrapod PSF, and 6 μm Tetrapod PSF. The initial structure for these simulations was taken from the molecule positions of the microtubule dataset MT0.N1.HD in the SMLM challenge[16]. This initial structure was duplicated multiple times with random rotations and 3D translations to create a denser distribution of microtubules within the same field of view. The original coordinates, with an approximate axial range of 1.4 μm for the astigmatic PSF, was scaled by factors of 2 and 4 to match the 3 μm and 6 μm Tetrapod PSFs, respectively.

Once the molecule positions were established, we utilized the photophysical model from the SMLM challenge[16] to determine the timing and occurrence of blinking events, which is different from the training data generation. The average durations of the on, dark and bleach states were set to 3, 2.5, and 1.5 frames, respectively. For all blinking events, the photon count was sampled from a broad uniform distribution [500, 10500], and a constant background of 50 photons was applied to all datasets. Three different densities (low, medium, and high) were simulated for each PSF to reflect a range of experimental conditions. To further mimic real-world scenarios where unknown aberrations are present, we introduced varying levels of coma and spherical aberrations to these datasets. Specifically, the aberrations denoted by Zernike coefficients $C_3^{-1}$, $C_3^1$ and $C_4^0$ were set to −20, 20, and −10 nm for the astigmatic PSF; −20, 20, and −50 nm for the 3 μm Tetrapod PSF; −30, 30, and −70 nm for the 6 μm Tetrapod PSF, respectively. The wavelength for all simulations was set as 670 nm.

**Noise simulation.** The simulation process began with the convolution of molecule positions with the respective PSFs to generate noise-free images. To introduce realistic noise, we employed the camera model specified in the SMLM challenge[16]. For the electron-multiplying charge-coupled device (EMCCD) camera, the pixel values $d_{\mathrm{ADU}}$ are given by:

$$d_{\mathrm{ADU}} = \frac{\mu_{\mathrm{o}e}}{e_{\mathrm{ADU}}} + \mathrm{BL} \tag{S21}$$

where

$$\mu_{\mathrm{o}e} = \mathrm{Gamma}\left(\mu_{\mathrm{i}e}, EM_{\mathrm{gain}}\right) + \mathrm{Gauss}(0, \sigma_R) \tag{S22}$$

and

$$\mu_{\mathrm{i}e} = \mathrm{Poisson}\left(\mathrm{QE} \cdot \mu_{\mathrm{phot}} + c\right) \tag{S23}$$

In these equations, $\mu_{\mathrm{phot}}$ represents the expected number of photons received by the camera pixel, calculated using the imaging formation model described in equation (S3). $\mathrm{QE}$ is the quantum efficiency, and $c$ is the spurious charge measured in electrons. Shot noise is introduced via a Poisson distribution to sample the input electrons $\mu_{\mathrm{i}e}$. This signal then undergoes an electron multiplication process, characterized by a Gamma distribution with the electron-multiplying gain $EM_{\mathrm{gain}}$ as the scale parameter. Readout noise is added as a zero-mean Gaussian distribution with a standard deviation $\sigma_R$. Finally, the output electrons $\mu_{\mathrm{o}e}$ are converted into pixel values $d_{\mathrm{ADU}}$ using the analog-to-digital conversion factor $e_{\mathrm{ADU}}$ and the camera baseline $\mathrm{BL}$. For the scientific complementary metal-oxide semiconductor (sCMOS) camera, the simulation does not include an electron multiplication step. Instead, only Poisson shot noise and Gaussian readout noise are considered in the calculation of the final pixel values. In this work, to simulate the microtubule datasets, we used the sCMOS model (ORCA-Flash4.0 V3, HAMAMATSU) with the following parameters provided by the manufacture: $\mathrm{QE} = 0.81, c = 0.002, \sigma_R = 1.61, e_{\mathrm{ADU}} = 0.47$, and $\mathrm{BL} = 100$.

**Supplementary Note 4 Benchmarking with other methods**

LUNAR is a method capable of simultaneously performing high-density localization and in situ PSF learning. To evaluate its performance, we conducted comparative benchmarks focusing on these two key capabilities.

**High-density localization.** We compared LUNAR with three state-of-the-art (SOTA) deep learning-based localization methods: FD-DeepLoc[17], DECODE[8], and DeepSTORM3D[9], all of which have demonstrated strong performance on high-density data. Among these, DeepSTORM3D and DECODE are designed for shift-invariant systems, while FD-DeepLoc extends DECODE to spatially variant systems with modifications to enhance performance. All three rely on PSF-supervised 3D localization learning (**Fig. 1b**). For fair comparison, we used their open-source codes and implemented them within our unified framework AI4Loc (Li-Lab-SUSTech/LUNAR), where LUNAR was also implemented. This ensured that all algorithms used the same data simulator for training and evaluation, minimizing inconsistencies due to implementation differences.

For network training, LUNAR, FD-DeepLoc, and DECODE were trained for 40,000 iterations using 16 randomly simulated images per iteration. To implement FD-DeepLoc's robust training strategy, Gaussian noise with a standard deviation of $\lambda/100$ was added to the Zernike coefficients of the PSF model during training data generation. We found DeepSTORM3D required more iterations and greater GPU memory, so it was trained for 100,000 iterations using the maximum number of images that could fit into 24 GB of GPU memory (NVIDIA GeForce RTX 4090), corresponding to 16 images with 64×64 pixels (astigmatic PSFs) or 6 images with 128×128 pixels (Tetrapod PSFs). All other training parameters including optimizers, learning rates, and learning rate schedulers were kept consistent with those reported in the original publications.

For post-processing, LUNAR, FD-DeepLoc, and DECODE employed multi-channel prediction maps that include a probability channel to identify emitters. A common threshold of 0.7 was applied for all three methods. DeepSTORM3D uses an upsampled volumetric grid to represent emitter positions. We used a four-fold lateral up-sampling and a 120-channel z-discretization. For simulated data with a pixel size of

100 nm, this resulted in output voxel sizes of 25×25×11.67 nm for astigmatic PSFs and 25×25×50 nm for 6 μm Tetrapod PSFs. Since DeepSTORM3D includes two hyperparameters that balance detection accuracy and localization error, we performed a grid search over threshold values [5, 10, 20, 30, 40, 80] and peak finding radii [2, 4, 5, 6, 8, 10] for each trained model. The parameter combination that maximized 3D localization efficiency on the validation set was selected for testing. Final results were then reported using the fixed post-processing settings.

**In situ PSF learning.** We compared LUNAR with two in situ PSF modeling methods: uiPSF[11] and INSPR[10]. Both methods rely on extracting isolated emitter images to build a PSF library, which is then used to iteratively refine the PSF model using either maximum likelihood estimation or a modified Gerchberg–Saxton algorithm. Because the accuracy of these approaches depends heavily on the quality of emitter detection and cropping, we tested multiple combinations of peak finding and segmentation parameters for each method and selected the configuration that yielded the best performance.

For the Tetrapod PSF, the parameters used in INSPR were: box size 71, distance threshold 8, and initial and segmentation thresholds both set to 25 to allow for more sensitive emitter detection. For uiPSF, we used an ROI size of 61, a blur kernel size of 24, and a maximum kernel size of 25. For the astigmatic PSF, INSPR was run with a box size of 40, distance threshold of 4, and initial and segmentation thresholds again set to 25. The uiPSF parameters were adjusted based on sample type: an ROI size of 31 was used for sparse beads and 23 for random emitters, along with a blur kernel size of 2 and a maximum kernel size of 3. All other parameters not specific to the optical system were kept at their default values.

**Supplementary Note 5 Computation of theoretical localization precision limit**

The Cramér-Rao Lower Bound (CRLB) provides a theoretical limit on the variance of any unbiased estimator[18], which can be utilized to quantify the localization precision in SMLM. The CRLB is calculated by the inverse of the Fisher information matrix:

$$var(\hat{h}_i) \geq \mathrm{CRLB}_{h_k} = [I(h)^{-1}]_{ii} \tag{S24}$$

where $\hat{h}_i$ represents the estimator of the molecule parameter $h_i$, and in this work, $h = [x, y, z, \mathrm{photon}, \mathrm{bg}]$. $I(h)$ is a 5×5 Fisher information matrix, with each element calculated as the expectation of the partial derivatives of the log likelihood with respect to the parameters:

$$I_{ij}(h) = \mathrm{E}\left[\frac{\partial \log p(d|h)}{\partial h_i}\frac{\partial \log p(d|h)}{\partial h_j}\right] \tag{S25}$$

where $p(d|h)$ denotes the likelihood of a Poisson process, as defined in equation (S2). By solving the above equation and applying the Stirling approximation ($\ln n! \approx n\ln n - n$), we obtain:

$$I_{ij}(h) = \sum_k \frac{1}{\mu_k}\frac{\partial \mu_k}{\partial h_i}\frac{\partial \mu_k}{\partial h_j} \tag{S26}$$

where $\mu_k$ is the expected number of photons in pixel $k$ from the PSF model function. The partial derivatives of the vectorial PSF model, as described in **Supplementary Note 2**, are given by:

$$\frac{\partial u_k}{\partial h_{bg}} = 1 \tag{S27}$$

$$\frac{\partial u_k}{\partial h_{phot}} = \mathrm{PSF}_k \tag{S28}$$

$$\frac{\partial u_k}{\partial h_{x,y,z}} = \frac{h_{phot}}{C}\sum_{p=x,y}\sum_{d=x,y,z}\mathcal{R}\{E_{p,d}^{\mathrm{img}^*}\frac{\partial E_{p,d}^{\mathrm{img}}}{\partial h_{x,y,z}}\} \tag{S29}$$

where $C$ is a normalization factor, $\mathcal{R}$ represents taking the real part. Substituting equation (S15) into the derivative we can get:

$$\frac{\partial E_{p,d}^{img}}{\partial h_{x,y,z}} = \mathcal{F}_{2D}\left\{ik_{x,y,z}A(\rho,\varphi)E_{p,d}^{\mathrm{pupil}}e^{i\psi_{\mathrm{pos}}}e^{i\psi_{\mathrm{aber}}}\right\} \tag{S30}$$

with the wavevector $k_{x,y,z}$ expressed as:

$$\frac{2\pi}{\lambda}\left(NA\rho\, cos\, \varphi\,, NA\rho\, sin\, \varphi\,, \sqrt{n^2-\rho^2 NA^2}\right) \tag{S31}$$

The physical meanings of the other terms have been defined in **Supplementary Note 2**. To calculate the multi-frame CRLB for a single emitter with constant brightness, we divide the single-frame CRLB by the square root of the temporal context length.

**Supplementary Note 6 Evaluation metrics**

To evaluate the performance on the simulated dataset with known ground truth, we utilized the Jaccard Index (JI), the Root Mean Squared Error (RMSE), and 3D efficiency as evaluation metrics. The Jaccard Index assesses the algorithm's ability to accurately identify blinking emitters while minimizing false predictions:

$$\mathrm{JI} = \frac{\mathrm{TP}}{\mathrm{TP} + \mathrm{FP} + \mathrm{FN}} \tag{S32}$$

where $\mathrm{TP}$ (True Positive) is the number of predicted emitters that can be matched to the ground truth emitters. $\mathrm{FP}$ (False Positive) is the number of predicted emitters that do not correspond to any ground truth emitters. $\mathrm{FN}$ (False Negative) is the number of ground truth emitters that are not identified by the predictions. The matching thresholds for lateral and axial displacements are set to 250 nm and 500 nm, respectively. To further quantify the precision of the true positive predictions, the RMSE measures the error between the predicted emitter positions and the ground truth positions:

$$\mathrm{RMSE_{3D}} = \sqrt{\frac{1}{\mathrm{TP}}\sum_{i}^{\mathrm{TP}}(\hat{x}_i - x_i^{\mathrm{GT}})^2 + (\hat{y}_i - y_i^{\mathrm{GT}})^2 + (\hat{z}_i - z_i^{\mathrm{GT}})^2} \tag{S33}$$

where $\hat{x}_i, \hat{y}_i, \hat{z}_i$ are the coordinates of the predicted true positive emitter $i$, and $x_i^{\mathrm{GT}}, y_i^{\mathrm{GT}}, z_i^{\mathrm{GT}}$ are the corresponding ground truth coordinates. Efficiency is a single metric that reflects both the software's ability to find emitters and to precisely localize them. It is given by ref.[16]:

$$\mathrm{Efficiency} = 100 - \sqrt{(100 - 100 \times \mathrm{JI})^2 + \alpha^2 \mathrm{RMSE}^2} \tag{S34}$$

Here, lateral and axial efficiencies are calculated with $\alpha = 1$ and $\alpha = 0.5$, respectively. The 3D efficiency is calculated as the average of the lateral and axial efficiencies.

**Supplementary Note 7 Data post-processing, analysis and rendering**

**Drift correction.** In SMLM experiments, data acquisition can extend from several minutes to tens of minutes, during which drift between the sample and imaging optics may introduce reconstruction artifacts. To address this issue, we employed the drift correction plugin in SMAP[19] software, which utilizes the redundant cross-correlation algorithm[20]. The drift correction was conducted with a spatial resolution of 10 nm and a time step size of 1/10 of the total frames.

**Image rendering.** SMAP software was used to render all super-resolution images in this work. To filter bad localizations, we normally set a detection probability threshold $\hat{p}_k$ of 0.7. Temporal grouping was not applied to average the positions of molecules detected multiple times in adjacent frames, as the investigated networks already incorporate temporal context for localization. Vutara SRX software (Bruker) was utilized for generating 3D super-resolution videos.

**Lateral offset correction.** The output representations of LUNAR, FD-DeepLoc and DECODE contain two lateral sub-pixel offsets $\widehat{\Delta x_k}, \widehat{\Delta y_k}$ relative to the pixel center for each detected emitter, with corresponding uncertainties $\sigma_{x,k}^2, \sigma_{y,k}^2$. Under challenging imaging conditions such as high emitter densities, large defocus, and low signal-to-noise ratios, the predicted Gaussian distribution within the pixel can become very flat (large uncertainties), with means $\widehat{\Delta x_k}, \widehat{\Delta y_k}$ concentrated near zero (the pixel center). This bias scales with the predicted uncertainties and can introduce grid artifacts if localization uncertainties are ignored during image reconstruction[8]. To solve this, we applied a post-processing correction to the lateral offsets $\widehat{\Delta x_k}, \widehat{\Delta y_k}$ without affecting localization accuracy. Specifically, all localizations were divided into 20 bins based on their lateral uncertainties $\sigma_{x,k}^2, \sigma_{y,k}^2$. For each bin's localizations, histogram equalization was applied to lateral sub-pixel offsets $\widehat{\Delta x_k}, \widehat{\Delta y_k}$ individually.

**NPC geometric analysis.** For automatic NPC geometric analysis, we used the pipeline provided in the reference[21], implemented as a series of plugins in SMAP. The analysis pipeline includes several steps, and we normally used them with default settings. Specifically, the rendered super-resolution NPC image was first convolved with a

Gaussian ring kernel to identify local maxima NPC candidates using a cutoff value of 0.06. Then these candidates were passed through a two-step clean-up process: (1) Localizations of each candidate were fitted to a circle and discarded if the fitted radius was too small (< 40 nm) or too large (> 70 nm); (2) Remaining candidates were re-fitted using a circle with a fixed radius of 55 nm. Candidates were rejected if more than 30% of localizations were within 45 nm or if more than 75% of localizations were more than 65 nm away from the NPC center, compared to the localizations on the ring. After segmentation, the filtered NPC candidates were fitted using a circular model, with center position and radius as free parameters.

**Supplementary Note 8 Implementation details about LUNAR**

**Localization learning (LL).** LUNAR LL utilizes training data simulated online. During each iteration, the data simulator generates a batch of training samples according to a predefined shape $[batch\,size, context\,size, height, width]$. Each sample in the batch contains a sequence of frames generated using a simple photophysical model. For networks set with the temporal context length $n$, additional $n-1$ frames are simulated per sample to provide temporal context for the middle frames, on which localization loss is computed exclusively. For example, given the batch shape of $[1,3,128,128]$ and the temporal context length of 3, the network receives a batch shaped as $[1,5,128,128]$, and outputs predictions for the middle 3 frames of each sample (**Fig. 2j**). In this work we normally used a batch size of 2, a context size of 8 and an image size of 128×128 for training. The AdamW optimizer[22] was used for network parameters updating, with an initial learning rate of $6\times10^{-4}$ and a weight decay of 0.05. The cosine annealing schedule was applied, ensuring the learning rate decays smoothly over the training period.

**Synchronized learning (SL).** LUNAR SL is trained by alternating between the localization learning and physics learning iterations. Given that the network encoder may poorly approximate the posterior function at the beginning, we initiated the synchronized learning with a "warm-up" phase consisting of 5,000 localization learning iterations. This allows the network to stabilize before introducing physics learning. Then we conducted the localization learning and physics learning in an interleaved manner: for every $interval$ updates of the network encoder, we performed a single update of the physics decoder. In this work we used the $interval$ of 2. The optimizer and learning rate schedule used for localization learning is the same as the above. For the physics learning, the physics decoder was updated using the Adam optimizer[23] with an initial average learning rate of $1\times10^{-2}$, which was multiplied by 0.9 every 1,000 iterations.

To address the typically non-uniform z-distribution of emitters in raw SMLM data, we designed an adaptive ROI selection strategy to construct a balanced ROI library for physics learning. Specifically, every 5,000 localization learning iterations, the current network is used to analyze the raw data in an ROI-wise manner, and the resulting localizations are stored for each ROI. The ROI library is then updated greedily, preferentially selecting both sparse ROIs (containing isolated emitters) and dense ROIs (with multiple overlapping emitters), to achieve a more uniform distribution of emitters across the axial range. This ensures robust learning of the in situ PSF model.

**Model inference.** We developed a data analyzer to perform inference on experimental SMLM data using trained models. The process begins with the analyzer inspecting the entire data folder and generating a reading plan to process the data. Users can specify the block size (1 GB used in this work) to manage CPU memory usage when handling large datasets. Upon loading a data block into CPU memory, we pad it with extra frames at both ends and split it into batches with temporal overlaps. The padding and overlap lengths are determined by the network's temporal context length, ensuring each frame within a batch has a multi-frame context. For cases where image sizes are too large or GPU memory is limited, the analyzer employs a divide-and-conquer strategy. It splits large images into smaller patches with controlled overlaps. To avoid duplication and ensure consistency, localizations that fall within the overlapped areas of these patches are discarded.

To obtain localization coordinates from multi-channel prediction maps, we utilized the same method as used in previous works[17]. Briefly, local maximum detection was performed on the predicted probability channel to identify emitter candidates: local maxima with $\hat{p}_k > 0.3$ or non-maxima pixels with $\hat{p}_k > 0.6$. The probability values of each candidate and its four adjacent pixels were summed to form a new probability map $\hat{p}_k^{new}$. A threshold of 0.7 was then applied to $\hat{p}_k^{new}$ to generate a binary map, indicating whether a pixel contains an emitter. Using this binary map, we indexed the 3D positions $\hat{x}_k, \hat{y}_k, \hat{z}_k$ and photon counts $\widehat{phot}_k$, along with their associated uncertainties $\sigma_{x,k}^2, \sigma_{y,k}^2, \sigma_{z,k}^2, \sigma_{phot,k}^2$, to compile the final localization list.

**Computational cost**. The LUNAR learning and inference were performed on a workstation equipped with an Intel Core i9-13900K processor (128 GB RAM) and an NVIDIA GeForce RTX 4090 graphics card (24 GB memory). For LUNAR LL with a

7-frame temporal context, training for 40,000 iterations using a batch size of 2, context size of 8, and image size of 128×128 took approximately 2.4 hours and utilized about 17 GB of GPU memory. LUNAR SL with the same settings required approximately 6.5 hours and 23 GB of GPU memory due to the incorporation of physics learning iterations. Inference on the demo-3 experimental dataset (100,000×296×296, 18 GB) using the LUNAR network with a batch size of 32 took approximately 16 minutes.

**Potential ill-posedness**. The joint estimation of the PSF and positions of overlapping emitters can be ill-posed under extreme conditions, e.g., the sample is too flat to contain axial structures, emitter density is very high, or the PSF changes weakly with defocus (**Supplementary Fig. 5a**). In such cases, the captured data provides limited information for PSF learning. This information deficiency explains LUNAR's performance drop under astigmatic PSF with 0.1λ aberrations (**Supplementary Fig. 6b**), where the model converges to a local minimum (**Supplementary Fig. 24**). This ill-posedness can be mitigated by incorporating appropriate priors, such as restricting the learned aberrations to lower-order modes or collecting raw data from multiple planes[10,24]. But in general, LUNAR demonstrates strong stability and robustness in practice, with convergence failures being rare in our experiments.
